# SpatialJEPA: Graph-context distillation enables spatially grounded single-cell representations across molecular assays

**DOI:** 10.64898/2026.07.21.739810

**Authors:** Dylan Mann-Krzisnik, Yue Li

## Abstract

Spatial multiomics links chromatin, transcriptomic, and metabolic states to tissue organization, but large single-cell atlases typically lack spatial coordinates and matched metabolite measurements. We introduce SpatialJEPA, a framework inspired by joint-embedding predictive architectures that distills tissue-neighborhood context into molecular representations usable without coordinates. A teacher receives spatial multiomic profiles and their neighborhood graph, whereas a student predicts its latent representations from the same profiles with neighborhood connections masked. With MultiGATE for spatial RNA–ATAC integration, the student preserves cross-modal matching and spatially organized programs. Applied without retraining to independent dissociated 10x Multiome nuclei, it supports more consistent spatial-reference placement of paired RNA and ATAC profiles than non-spatial baselines. Extending SpatialJEPA to a graph-augmented SpatialMETA backbone for spatial transcriptome–metabolome data, the student preserves broad tissue organization in sections from held-out mice. A source-defined dopamine-associated component transfers to RNA–ATAC targets lacking measured metabolites, recovering a concordant striatal transcriptional program and chromatin weights enriched for D2 medium spiny neuron regulatory elements. Finally, we apply a model trained on human spatial RNA–MSI data to 714,534 caudate-head nuclei from 42 donors in the Seattle Alzheimer’s Disease Brain Cell Atlas. A component with dopamine among its strongest inferred metabolite weights distinguishes medium spiny neurons from oligodendrocytes, consistent with caudate gray–white organization in Xenium tissue. By using spatial multiomics as supervision rather than only as a mapping reference, SpatialJEPA carries tissue context into representations of atlas-scale dissociated cells, a step toward spatially grounded multimodal models of cellular state.

## 1 Introduction

Single-cell multiomics has expanded the study of gene regulatory programs and functional cell states by measuring complementary molecular layers within the same cells (Wang et al. 2026). Spatial multiomics adds tissue context to these measurements, linking transcriptomic, chromatin, and metabolic states to local organization and microenvironment (Zhang et al. 2023; Vicari et al. 2024). This context is particularly important in the brain, where molecular programs are organized across anatomical compartments, cellular neighborhoods, and disease-associated microenvironments. In Alzheimer’s disease, for example, amyloid plaques are associated with local transcriptional programs and spatially patterned microglial responses (Keren-Shaul et al. 2017; Wood et al. 2022). Spatial metabolomics provides a complementary view of biochemical state by directly resolving metabolite distributions within tissue (Vicari et al. 2024). Yet these assays remain more difficult to scale than dissociated single-cell sequencing. Large atlases are therefore dominated by dissociated cells that lack spatial context, whereas spatial multiomic and metabolomic studies typically profile fewer samples and tissue sections (The Tabula Sapiens Consortium 2022; Li et al. 2021). As a result, datasets with the broadest cohort coverage often lack the spatial and metabolic information needed to connect molecular state to tissue organization and biochemical function.

Existing computational approaches address complementary parts of this problem. Spatial multiomic methods integrate molecular modalities within spatially resolved tissue: MultiGATE models spatial RNA–ATAC data (Miao et al. 2025), SpatialGlue uses graph-based integration across spatial omics (Long et al. 2024), and SpatialMETA jointly models spatial transcriptomic and metabolomic measurements (Tian et al. 2025). Methods such as MultiGATE use spatial neighborhood graphs to incorporate tissue context during training and inference, but this context is unavailable when embedding dissociated cells without coordinates. SpatialMETA instead uses coordinates to co-register transcriptomic and metabolomic measurements; its standard encoder does not model spatial neighborhoods. A second family of methods bridges spatial and dissociated transcriptomic data. Tangram maps single-cell RNA-seq profiles onto a spatial reference (Biancalani et al. 2021), gimVI jointly models single-cell and spatial transcriptomics to impute genes not measured by the spatial assay (Lopez et al. 2019), and ENVI uses COVET neighborhood representations to infer spatial context for dissociated cells (Haviv et al. 2025). ENVI is closest to our goal, but focuses on transcriptomic neighborhood reconstruction rather than transferring representations learned by spatial multiomic models. Together, these approaches integrate spatial modalities, map dissociated cells onto spatial references, or infer aspects of spatial context, but do not provide a general mechanism for distilling spatial multiomic representations into an encoder that operates without coordinates across modalities such as RNA, chromatin accessibility, and metabolites.

Joint-embedding predictive architectures (JEPAs) offer a natural route to such transfer. Motivated by predictive world models, JEPAs learn representations by predicting target representations in latent space rather than reconstructing the input (Assran et al. 2023). Contemporaneous work has begun adapting this principle to single-cell and spatial biology. GeneJEPA predicts latent representations of masked gene sets from visible expression context (Litman et al. 2025), and TERRA uses JEPA-based self-supervision over cells and their spatial neighborhoods to learn gene-, cell-, and neighborhood-level representations from spatial transcriptomics (Birk et al. 2026a). In TERRA, however, spatial neighborhoods remain part of the input from which representations are constructed. Neither approach addresses the complementary problem considered here: transferring context learned from spatial data into a model that operates when spatial information is unavailable. Here, we introduce SpatialJEPA, a teacher–student framework that uses JEPA-style latent prediction to transfer spatial context into non-spatial molecular representations. A teacher with access to the tissue graph supervises a student that learns to predict the teacher’s latent representation from the same molecular profiles with neighborhood connections masked, allowing the trained student to operate without coordinates. Because the framework does not depend on a particular multiomic backbone, we instantiate it with MultiGATE for RNA–ATAC integration and a graph-augmented SpatialMETA for transcriptome–metabolome integration.

In spatial RNA–ATAC data, the SpatialJEPA student preserves cross-modal integration, spatially organized molecular programs, and recoverable positional information without a spatial graph at inference. It also transfers without retraining to dissociated 10x Multiome nuclei, recovering spatial teacher-defined domains more faithfully than non-spatial baselines. In spatial transcriptome–metabolome data, graph-context distillation organizes measured dopamine in the source latent space and preserves broad tissue organization in sections from held-out animals. A dopamine-associated component defined on the metabolomic source recovers an independently derived dopamine-linked transcriptional program in RNA–ATAC targets lacking measured metabolites, while its chromatin weights are enriched for D2 medium spiny neuron (MSN) cis-regulatory elements. Finally, we transfer a human striatal model to single-nucleus RNA-seq from the caudate head in the Seattle Alzheimer’s Disease Brain Cell Atlas (SEA-AD) (Hawrylycz et al. 2024; Kana et al. 2026). Despite the absence of metabolomic measurements in SEA-AD, the transferred representation recovers a striatal transcriptome–metabolome component in which dopamine is among the strongest metabolite weights and canonical MSN genes such as *PDE10A* and *ANO3* weigh positively. This component separates MSN from oligodendrocyte states, consistent with caudate gray–white organization. Together, these results demonstrate how spatial multiomics can provide supervision for non-spatial single-cell representations, connecting molecular states to tissue organization even when spatial coordinates and metabolite measurements are unavailable.

## 2 Results

### 2.1 SpatialJEPA overview

In SpatialJEPA, a teacher encoder is trained on paired spatial multiomic data with the full spatial graph, whereas a student encoder receives the same molecular profiles with a trivial self-only identity graph, depriving the student of neighboring tissue context (Fig. 1a). After each teacher update, the frozen teacher generates modality-specific embeddings and the student is updated to predict these embeddings under a mean squared error objective (Methods, *Graph-context distillation*). The student therefore learns a map from dissociated molecular profiles to a representation shaped by spatial context, and can be applied to non-spatial data using an identity graph at inference time. Notably, SpatialJEPA is agnostic to which spatial multiomic model is used as the teacher (Fig. 1b). Once the teacher and student models are trained, samples from single-cell data and “dissociated” graph-masked spatial data can be co-embedded within the student model’s latent space (Fig. 1c). In this co-embedded space, we use optimal transport (OT) (Peyré and Cuturi 2020) to learn a transport plan from single-cell data to spatial data, and transport the former via barycentric transport to better align with spatial data. Following this alignment, each transported single-cell profile is assigned the coordinates of its nearest spatial reference profile in the co-embedded space for downstream spatial mapping (Methods, *Zero-shot transfer and optimal-transport alignment*). These coordinates are used only for this post-hoc lookup, not as inputs to the student encoder. Another key feature of SpatialJEPA’s latent space distillation is that the student model can forego a decoder during training, while still indirectly optimizing for decodability via the teacher’s training objective when the latter relies on a reconstruction loss. This feature allows fitting interpretable student decoders in a *post-hoc* fashion and then mapping the learned decoder features onto spatial coordinates (Fig. 1d). For interpretability, we primarily use a partial least-squares (PLS) regression model (Tsuchiya et al. 2022) as a *post-hoc* student decoder, which projects the latent embedding onto a small number of components and relates those components back to measured molecular features (Methods, *Post-hoc PLS decoders*).

**Figure 1:**
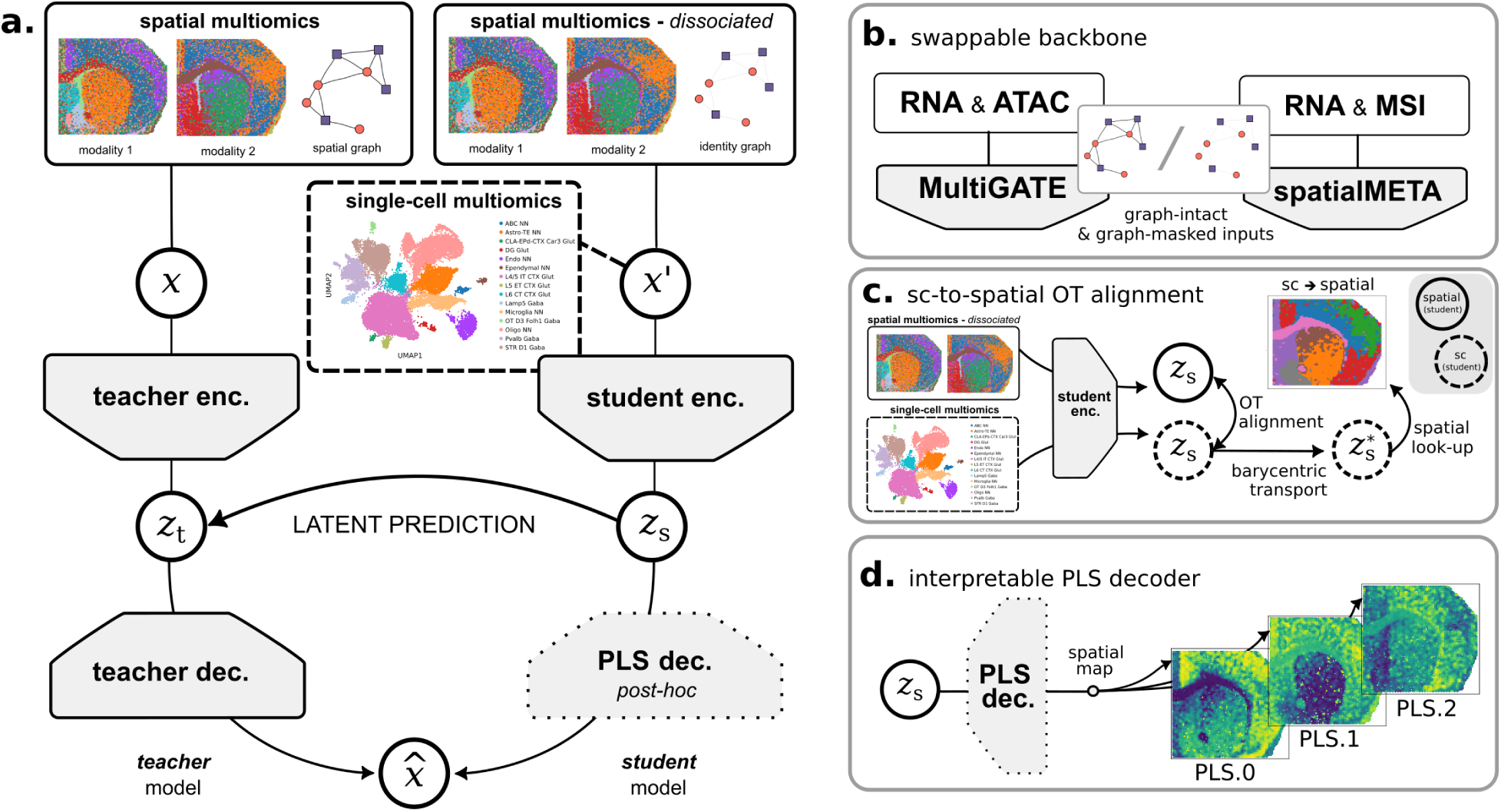
Overview of SpatialJEPA. **(a)** Graph-context distillation. A teacher encoder receives the spatial source data **X** with its full tissue-neighborhood graph; a student encoder receives **X***^′^*, the same molecular profiles with the graph replaced by a self-only identity graph that mimics dissociation. The student is trained to predict the teacher’s latent representation **z***_t_* from its own **z***_s_*. The trained student can be applied zero-shot to non-spatial single-cell multiomics (dashed box). A partial least squares (PLS) decoder is fitted *post-hoc* to the student embedding and is not part of the distillation objective (dotted box). **(b)** Each teacher–student pair shares the same backbone architecture, differing only in the graph supplied to the encoder. The backbone is selected according to the input modalities: MultiGATE for RNA–ATAC and a modified SpatialMETA for RNA–MSI. The main condition is that the encoder can operate on either graph-intact or graph-masked inputs to enable teacher-student distillation. **(c)** Optimal transport (OT) alignment. Single-cell (sc, dashed) and “dissociated” graph-masked spatial data (solid) are both passed through the trained student encoder, yielding student embeddings **z_s_** for both. OT is applied to compute a coupling between the datasets, after which the single-cell embeddings are mapped onto the spatial data embeddings using barycentric transport, becoming *z^*^_s_*. Finally, each transported cell inherits the coordinates of its matched spatial location by look-up of the closest spatial data embedding. **(d)** A *post-hoc* PLS decoder projects the student embedding onto a small set of interpretable components, which can then be related back to molecular features and mapped spatially.

In summary, SpatialJEPA allows spatial multiomics to supervise coordinate-free representations, while retaining separate downstream approaches for tissue localization and molecular interpretation.

### 2.2 Graph-context distillation preserves cross-modal integration and spatial context

Paired acquisition of scRNA-seq and scATAC-seq represents one of the most abundant single-cell multiomic modalities (Liang et al. 2023). As such, training SpatialJEPA on spatial RNA–ATAC and applying it to single-cell RNA–ATAC presents one of the most promising avenues for graph-context distillation. Following cross-dataset feature alignment (Methods, *RNA–ATAC feature harmonization and graph construction*), we trained teacher and student models on the postnatal day 22 (P22) mouse brain spatial RNA–ATAC dataset from Zhang et al. (2023) (Fig. 2a) and performed inference using the student on a dissociated 10x Multiome mouse brain sample (10x Genomics 2023). On the spatial RNA–ATAC training data, both the teacher (Fig. 2b) and the graph-masked student (Fig. 2c) successfully integrate the modalities in the latent space. Importantly, the student’s latent representation closely mimics the teacher’s in terms of UMAP geometry and alignment with reference RNA and ATAC cluster annotations. The student also integrates RNA and ATAC profiles from the dissociated 10x Multiome target without encoder retraining (Fig. 2d). Because the 10x Multiome data do not contain a curated cell-type annotation, we used CellTypist (Domínguez Conde et al. 2022) to predict their cell types based on the target RNA view; they serve as a common evaluation annotation across models rather than as ground truth.

**Figure 2:**
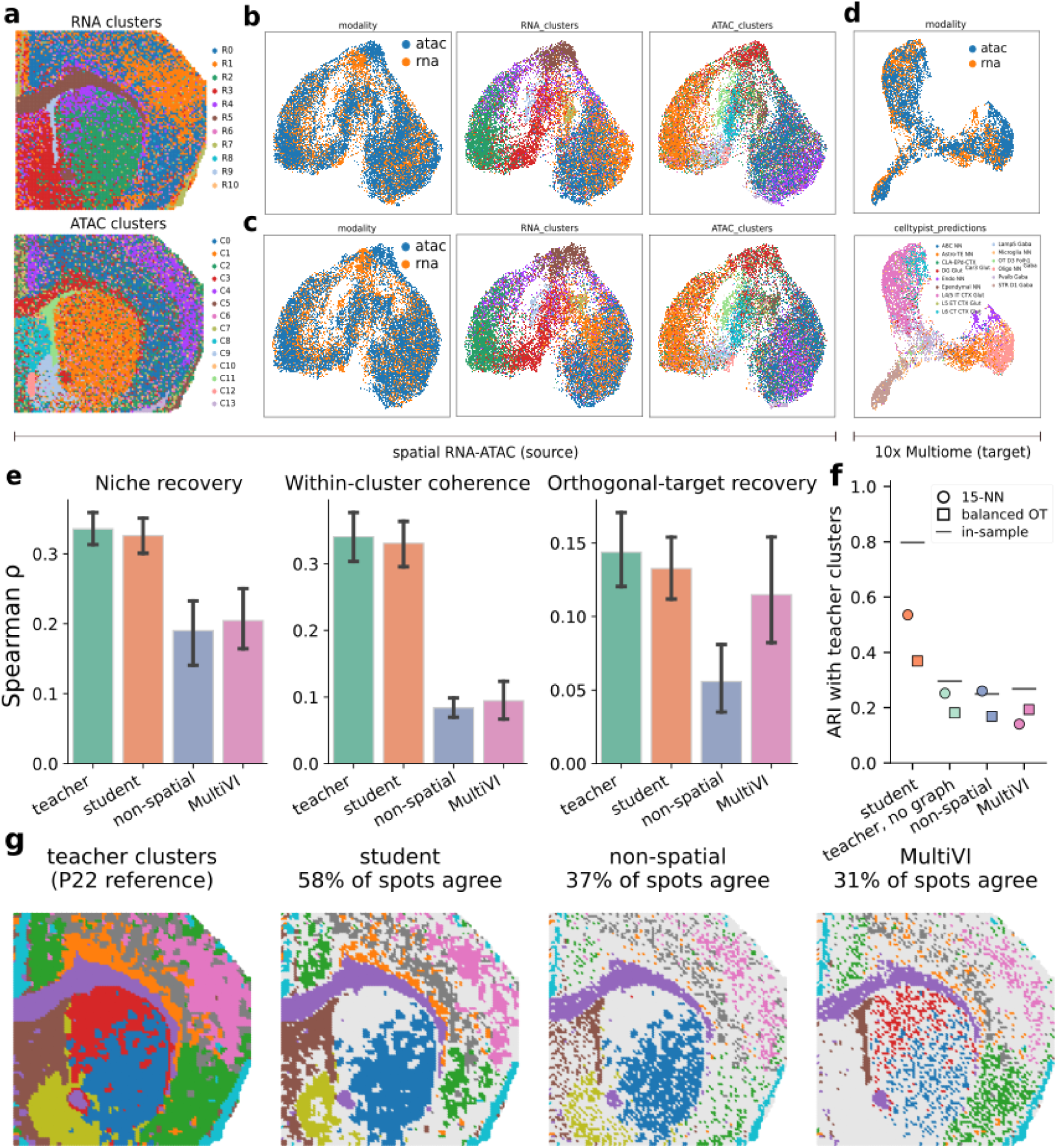
Source and target integration under graph-context distillation. **(a)** Spatial mouse RNA–ATAC data with RNA (top) and ATAC (bottom) cluster annotations from Zhang et al. (2023). **(b,c)** UMAPs of spatial-source embeddings from the teacher **(b)** and graph-masked student **(c)**, colored by modality (left), RNA cluster (middle), and ATAC cluster (right). **(d)** Zero-shot student embeddings of held-out 10x Multiome nuclei, colored by modality (top) and CellTypist-predicted cell type (bottom). Cell-type predictions are not ground-truth annotations. **(e)** Comparison of models on multimodal spatial metrics, quantified by Spearman correlation. Conventional integration metrics are reported in Supplemental Fig. S1. **(f)** Recovery of spatial teacher-defined P22 domains from held-out Multiome nuclei. Leiden clusters computed independently in each model’s Multiome embedding were assigned to P22 spots by 15-nearest-neighbor (circles) or balanced optimal-transport (squares) coupling. Agreement with the teacher domains is measured by adjusted Rand index (ARI), averaged over *K* = 5, 10, and 20 clusters. Horizontal dashes indicate in-sample P22 scores; “Teacher, no graph” denotes the teacher encoder evaluated with an identity graph. **(g)** Spatial distribution of teacher-defined P22 domains (*K* = 10) and their recovery from held-out Multiome nuclei under 15-nearest-neighbor coupling. Multiome clusters were matched one-to-one to teacher domains by maximal overlap; colored spots indicate agreement and gray indicates disagreement.

Conventional integration benchmarks (Supplemental Methods, *Conventional integration metrics*; Supplemental Fig. S1) quantify cross-modal pairing (1-FOSCTTM), modality mixing (iLISI), and preservation of annotated structure (biological conservation). On the spatial source, where all models are evaluated in-sample, the teacher and graph-masked student performed comparably, and both outperformed the non-spatial MultiGATE ablation, which shares the student’s architecture and identity graph but lacks teacher supervision. MultiVI (Ashuach et al. 2023) matched the SpatialJEPA models on pairing and mixed modalities more strongly, but preserved less annotated structure. On the dissociated target, on which no encoder was trained, the zero-shot student achieved the highest pairing and biological conservation, whereas source-trained MultiVI mixed modalities most strongly. OT alignment increased the student’s modality mixing but reduced its pairing and biological conservation.

The above three integration and clustering metrics assess whether paired RNA and ATAC profiles occupy a common latent space and whether annotated biological structure is preserved. Because none of them depends on tissue coordinates, however, strong integration does not by itself establish that the student has inherited the teacher’s representation of local tissue context. We therefore complemented them with three spatially informed metrics whose evaluated quantities are defined using physical neighborhoods or distances (Supplemental Methods, *Spatial-structure metrics*). Briefly, “niche recovery” predicts the cluster composition of each spot’s fifteen nearest physical neighbors from the joint embedding; “within-cluster coherence” correlates embedding distance with physical distance among spot pairs belonging to the same transcriptomic or chromatin cluster; and “orthogonal-target recovery” predicts the neighbor-averaged ATAC profile from the RNA-only embedding (Fig. 2e). Under spatial-block cross-validation on the source RNA–ATAC data, the student matches the teacher on all three metrics, whereas the non-spatial ablation retains roughly half the teacher’s niche recovery and a quarter of its within-cluster coherence. MultiVI yielded niche-recovery and coherence scores similar to those of the non-spatial ablation, while its orthogonal-target recovery was lower than the student’s, although their confidence intervals overlapped. The comparatively strong orthogonal-target recovery of MultiVI is consistent with this metric also capturing cross-modal integration, in addition to spatial context.

Relative position was also recoverable from the graph-masked student on the spatial RNA–ATAC source. Across 40 random half-splits of spots into query and reference sets, nearest-neighbor retrieval in the student latent placed 52% of queries within two spot widths of their true position, compared with 69% for the teacher and less than 1% for MultiVI and the non-spatial ablation (Supplemental Section S2; Supplemental Figs. S2, S3). Because all source spots were used for training, this measures positional information retained by the representation rather than generalization to unseen tissue.

Because the preceding benchmarks used source spots seen during training, we next assessed whether the learned spatial structure transfers to independent dissociated 10x Multiome nuclei. We clustered the nuclei in each model’s embedding, coupled their clusters to P22 spots, and measured agreement with Leiden domains defined from the spatial teacher’s P22 embedding (Supplemental Methods, *Placement of held-out Multiome nuclei on the spatial source*). The teacher domains also showed modest concordance with the published P22 RNA and ATAC annotations (ARI 0.29 and 0.21), comparable to the concordance between the published annotations themselves (0.23). Under 15-nearest-neighbor coupling, the student recovered the teacher domains with an ARI of 0.54, compared with 0.25 for the teacher evaluated without its graph, 0.26 for the non-spatial ablation, and 0.14 for MultiVI. With balanced optimal transport, the student achieved an ARI of 0.37, versus 0.17–0.19 for the other models (Fig. 2f). At *K* = 10, 58% of P22 spots received the matching teacher domain through the student, compared with 37% for the non-spatial ablation and 31% for MultiVI (Fig. 2g). Scored against the published RNA and ATAC annotations instead, the same placed clusters (*K* = 10) agreed to a similar extent across models (mean ARI 0.29 for the student versus 0.17–0.27 for the other models under 15-nearest-neighbor coupling, and 0.17 versus 0.18–0.19 under optimal transport; Supplemental Fig. S4), indicating that the student’s advantage is most pronounced when evaluated against spatially informed teacher domains rather than published modality-specific annotations. Together, these comparisons indicate that graph-context distillation carries the teacher’s spatial domain structure into representations of previously unseen dissociated nuclei, beyond what is recovered by graph-free teacher inference or non-spatial integration alone.

### 2.3 Transferred embeddings recover striatal and myelin programs in non-spatial single-nucleus data

Having shown that the student recovers teacher-defined spatial domains in held-out 10x Multiome nuclei (Fig. 2f,g), we next asked whether these representations also recover spatially organized transcriptomic and chromatin-accessibility programs. The spatial RNA–ATAC source was re-encoded with the student using the identity graph, while the same frozen student was applied zero-shot to the 10x Multiome mouse brain target data. We then aligned the target embeddings to the source by OT (Methods, *Zero-shot transfer and optimal-transport alignment*). Specifically, barycentric projection places target cells in the source latent frame, after which each cell inherits the coordinates of its nearest source spot. After barycentric OT alignment, Leiden clustering of the target representation, performed separately from the source, recovers a partition that closely matches the source tissue organization in both spatial-coordinate and UMAP views (Fig. 3a). Across genes significant in both source and target, per-Leiden Spearman correlations of differential-expression significance range from 0.16 to 0.46 (mean 0.37), indicating broadly concordant differential-expression patterns among matched Leiden clusters in the spatial source and aligned target (Fig. 3b; Supplemental Methods, *Marker-gene scores and source–target gene-rank concordance*).

**Figure 3:**
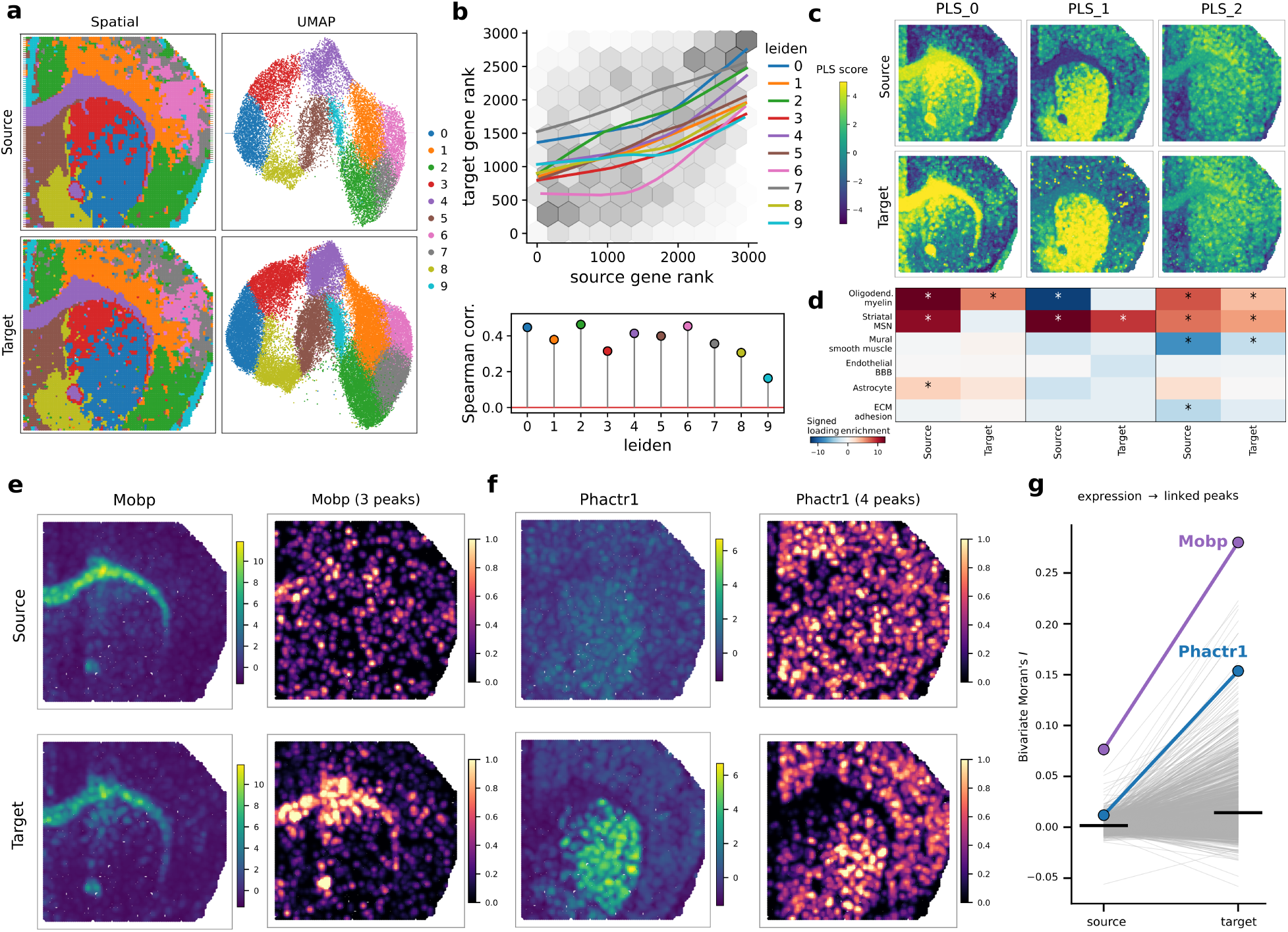
Source–target alignment of spatial and single-nucleus data. Target spatial views are OT-aligned. **(a)** Source spatial coordinates and UMAP (top row) alongside the OT-aligned target representation (bottom row), with Leiden clusters computed separately in source and target and target cluster labels matched to their source counterparts. **(b)** Source–target concordance of genes ranked by differential-expression significance, with higher ranks indicating stronger differential expression. Grayscale hexbins show overall concordance, colored traces show cluster-specific trends, and the lower panel reports per-Leiden Spearman correlations. **(c)** Spatial maps of the first three RNA PLS components (PLS_0–2), from models fitted separately in source and target from the OT-aligned embeddings to RNA features, on a shared color scale. The ATAC components are shown in Supplemental Fig. S5. **(d)** Signed enrichment of RNA PLS_0–2 gene weights for six curated cell-type and tissue-program gene sets, in source and target components of the same index after sign alignment; asterisks mark FDR *q <* 0.05. **(e)** The oligodendrocyte marker *Mobp* in source (top) and aligned target (bottom): RNA expression (left), averaged per spot and smoothed with a Gaussian kernel of one spot width, on one color scale for source and target; and accessibility of its three attention-linked peaks (right), smoothed in the same way and scaled to the pooled 99th percentile. **(f)** The same for *Phactr1* and its four linked peaks. **(g)** Bivariate Moran’s *I* between each gene’s expression and its linked-peak accessibility in source and aligned target, for the 2,163 shared genes with at least one retained gene–peak link. Gray lines denote genes, black bars means, and *Mobp* and *Phactr1* are highlighted.

To interpret the molecular structure encoded by these student representations, we fitted 9-component PLS models from the embeddings to measured RNA expression and ATAC accessibility, independently in source and target (Supplemental Methods, *PLS decoding of RNA–ATAC embeddings*). The resulting components provide interpretable molecular axes for identifying the genes and peaks represented in the student latent space. The spatial maps of the leading components show how these programs are distributed (Fig. 3c; ATAC components in Supplemental Fig. S5). In the source, PLS 0 spans broad regions of both white matter and striatum, whereas in the aligned target these programs are more distinctly resolved across PLS 0 and PLS 1. Consequently, the molecular signal represented by a single broad source component can be distributed across multiple, more sharply differentiated target components. Gene-set enrichment supports this interpretation (Fig. 3d). In the target, PLS 0 prominently captures the oligodendrocyte/myelin program, while PLS 1 resolves the striatal medium spiny neuron (MSN) program; PLS 2 captures contributions from both programs. The underlying myelin and striatal marker genes retain their spatial localization across source and aligned target, and their component weights follow the redistribution described above (Supplemental Fig. S6).

We next asked whether the student-derived target representation also recovers spatially organized regulatory structure at individual loci. For each gene with at least one gene–peak link retained from MultiGATE’s attention (Supplemental Methods, *Gene–peak link extraction and PLS weighting*), we measured the spatial coupling between its expression and linked-peak accessibility using bivariate Moran’s *I* on a common spatial lattice (Supplemental Methods, *Spatial coupling of expression and linked-peak accessibility*). Across the 2,163 genes, this coupling was stronger in the aligned target than in the source, with mean *I* increasing from 0.002 to 0.014. Spatial coupling increased for 67% of genes (Fig. 3g). Part of this increase may reflect the averaging of multiple target nuclei assigned to the same spatial location, which can reduce noise and sharpen spatial patterns. These results indicate that the transferred representation preserves spatially organized gene–peak associations and reveals stronger spatial coherence in the aligned target, supporting the recovery of regulatory structure from dissociated single-nucleus data.

The recovery of spatially coherent regulatory structure is illustrated by two genes with contrasting expression patterns. For *Mobp*, RNA expression already delineates similar white-matter structures in source and target, yet its linked-peak accessibility becomes more spatially coherent after transfer (Fig. 3e). Accordingly, bivariate Moran’s *I* increases from 0.076 to 0.280 (Fig. 3g). By contrast, *Phactr1* exhibits a relatively diffuse expression pattern in the source but resolves a distinct striatal pattern in the aligned target, accompanied by more spatially concentrated linked-peak accessibility (Fig. 3f). Its bivariate Moran’s *I* similarly increases from 0.012 to 0.154 (Fig. 3g). These examples illustrate stronger spatial gene–peak coupling after transfer, both when RNA localization is already conserved and when it becomes more clearly resolved in the aligned target.

Together, these results show that spatial-context distillation enables the student to recover coherent molecular and regulatory programs when transferred to an independent single-nucleus dataset lacking spatial measurements.

### 2.4 SpatialJEPA for spatial transcriptome–metabolome distillation

Whereas spatial RNA–ATAC is useful for investigating regulation of gene expression in different tissue contexts, it lacks the molecular layers needed to study the effect of downstream metabolites and neurotransmitters on brain tissue biology. On the other hand, spatial transcriptomics can be combined with mass spectrometry imaging (MSI) to study the relation between gene expression and such molecules, giving rise to spatial RNA–MSI data. For example, the spatial RNA–MSI data of Vicari et al. (2024) include samples from Parkinson’s disease (PD) mouse models that combine 10x Visium with MALDI-MSI, including an MSI modality that measures neurotransmitter abundance. We thus investigated whether SpatialJEPA can perform graph-context distillation on these spatial RNA–MSI data. Because SpatialJEPA can admit different spatial multiomic models as the backbone for training teacher and student models (Fig. 1b), we replaced the MultiGATE RNA–ATAC backbone with SpatialMETA (Tian et al. 2025), a conditional variational autoencoder for cross-modal integration of spatial RNA–MSI data. Importantly, although SpatialMETA uses spatial coordinates upstream to co-register transcriptomic and metabolomic measurements at matched tissue spots, its original neural encoder processes those paired spots independently without a spatial-neighborhood graph. Because SpatialJEPA requires a teacher representation that explicitly incorporates spatial context for its distillation objective, we replaced SpatialMETA’s fully connected encoder layers with graph-convolution layers that propagate information between neighboring spots while preserving the remainder of the backbone architecture (Methods, *Spatial and cross-modal extensions of SpatialMETA*).

Of note, the PD mouse model from Vicari et al. (2024) is characterized by a unilateral 6-hydroxydopamine lesion that depletes dopaminergic input to the striatum in one hemisphere, for which intact-versus-lesioned annotations are available (Fig. 4a; Methods, *Datasets and preprocessing*). This asymmetry is visible in the measured dopamine of the spatial RNA–MSI section as a spatially structured depletion that differs between hemispheres (Fig. 4b). This marked pattern can be used as a clear spatial signal on which to assess SpatialJEPA’s ability to distill graph context from mouse brain tissue. To establish a reference signature of this spatial pattern that is independent of SpatialJEPA, we decomposed the PD mouse model’s spatial transcriptome by non-negative matrix factorization (NMF) (Lee and Seung 1999) and ranked the resulting programs by bivariate Moran’s *I* against *measured* dopamine on the Visium spatial coordinates. Across NMF ranks 1–29, component 22 of the rank-29 decomposition showed the highest bivariate Moran’s *I* with measured dopamine (*I* = 0.478) and co-localized with dopamine in a unilateral striatal domain (Fig. 4c; Supplemental Methods, *NMF dopamine program*). Rank 29 was the upper bound of the tested range. NMF component 22 therefore provides a gene-loading ranking for a striatal transcriptional program spatially associated with measured dopamine, which we use as an external reference in subsequent analyses.

**Figure 4:**
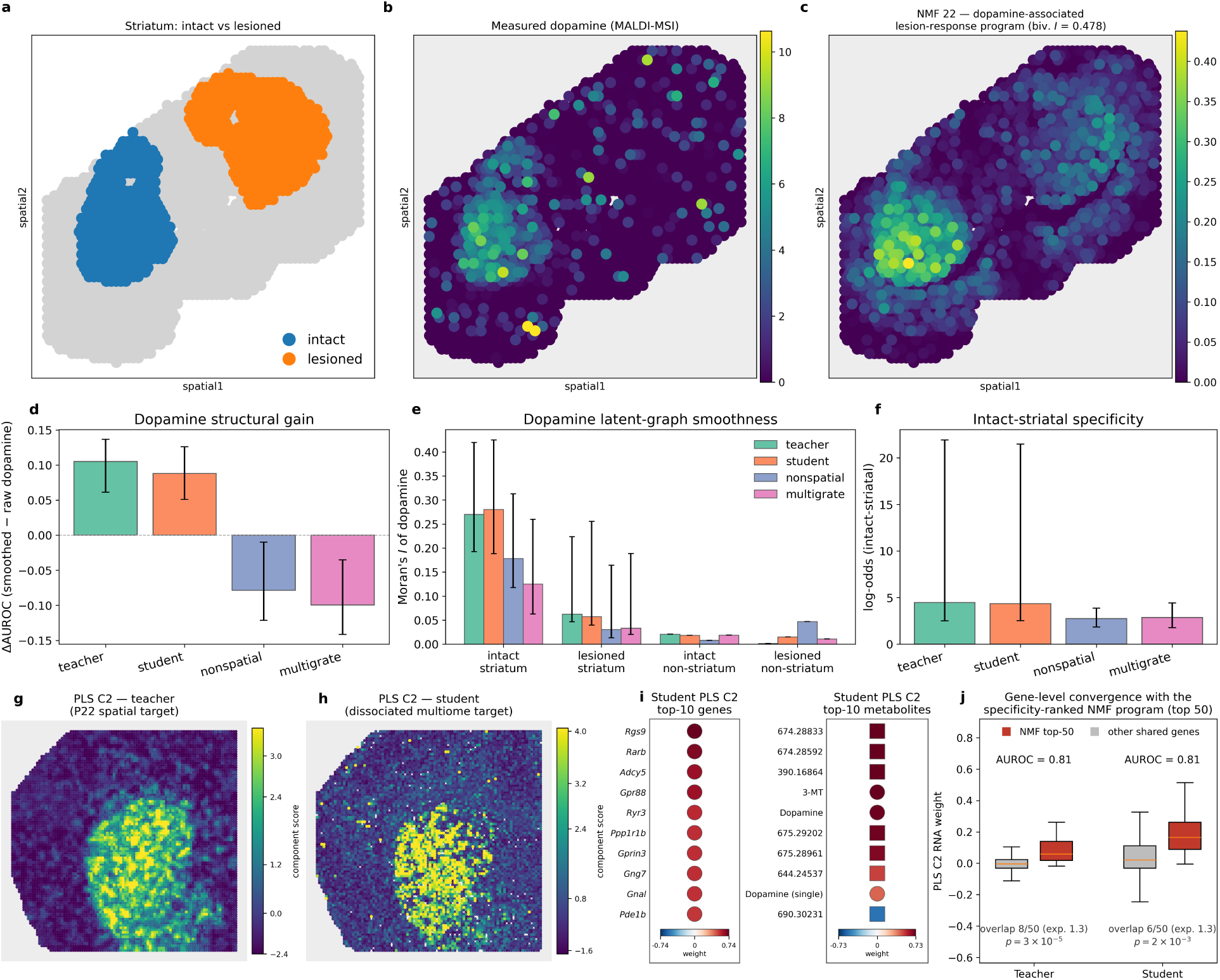
SpatialJEPA transfers a measured dopamine-associated lesion-response program. **(a)** The intact-versus-lesioned striatal axis in the PD mouse model from the spatial RNA–MSI dataset. **(b)** Measured dopamine (MALDI-MSI) depleted in one hemisphere by the unilateral 6-OHDA lesion. **(c)** NMF component 22, the gene-expression program most spatially co-varying with dopamine (bivariate Moran’s *I* = 0.478), used throughout as a reference dopamine-lesion gene signature. **(d–f)** Within-source embedding benchmark on the same spatial RNA–MSI section, comparing the spatial teacher, graph-masked student, non-spatial ablation and Multigrate; model legend in panel (e). Error bars are within-section spatial-block bootstrap intervals. **(d)** Structural gain, quantifying the change in intact–lesioned AUROC after neighbor smoothing on the latent graph. **(e)** Latent-graph Moran’s *I* by region and lesion status. **(f)** Log-odds ratio of intact-striatal membership among dopamine-high spots, relative to the section-wide base rate. **(g)** Teacher scores of the dopamine component C2 on the P22 spatial target and **(h)** student scores of C2 on the non-spatial 10x Multiome target. Both show the same component, defined once on the spatial RNA–MSI source and read out through the teacher and student RNA latents, respectively. Refer to Fig. 1c for spatial mapping procedure of dissociated data. **(i)** Top-10 gene and metabolite weights of the *student* C2 on Multiome data, with metabolite weights recovered in the nuclei by optimal-transport pairing with the source spots; marker size corresponds to absolute weight, marker color to the signed weight, and marker shape distinguishes annotated features (circle) from unannotated *m/z* MSI peaks (square). **(j)** Gene-level comparison between C2 and the NMF-derived signature shown in panel (c), showing teacher (P22) and student (Multiome) C2 RNA weights for the 50 genes most specific to NMF component 22 versus other shared genes, and the overlap between the NMF and C2 top-50 gene sets.

#### Spatial distillation sharpens the lesion-associated organization of dopamine in the latent space

After training teacher and student SpatialMETA models on the spatial RNA–MSI source data with SpatialJEPA, we first quantified how spatial distillation shapes the representation of this dataset. We calculated benchmarks that leverage the unilateral distribution of measured dopamine in the PD mouse model (Fig. 4a,b). Three metrics were used: structural gain (Fig. 4d), latent-graph smoothness (Fig. 4e), and intact-striatal enrichment of dopamine-high regions (Fig. 4f); all are defined in Supplemental Methods, *Dopamine-embedding benchmark*. Based on these metrics, we compared the spatial teacher, the graph-masked student, a non-spatial ablation, and Multigrate (Litinetskaya et al. 2025) as an external non-spatial multimodal-integration reference.

The decisive metric is structural gain (Fig. 4d), which asks whether neighbor-based smoothing sharpens or degrades lesion contrast. Whereas the teacher (0.11) and student (0.09) latent graphs sharpen intact–lesioned separation, with confidence intervals entirely above zero, the non-spatial ablation and Multigrate score negatively (*−*0.08 and *−*0.10, respectively), thus degrading the lesion contrast. Latent-graph smoothness corroborates the mechanism in terms of Moran’s *I* values (Fig. 4e). In addition to all models showing smoother dopamine signals on the latent graph in intact striatum compared to the other tissue compartments, latent dopamine signals are smoother for the teacher (0.27) and student (0.28) compared to non-spatial (0.18) and Multigrate (0.13). These observations are consistent with dopamine being depleted and spatially unstructured after lesioning. Moreover, intact-striatal specificity shows that all four models preferentially associate dopamine-high spots with intact striatum (Fig. 4f), with higher log-odds ratios for the teacher (4.5) and student (4.3) than for the non-spatial ablation (2.7) and Multigrate (2.9).

Taken together, these results indicate a consistent spatial advantage in how dopamine is organized in the SpatialJEPA teacher and student representations, clearest in terms of structural gain with directional support from smoothness and specificity. Because the analysis is based on one tissue section and relies on a within-section spatial-block bootstrap, the marginal intervals for Moran’s *I* and specificity should be interpreted as suggestive rather than strictly inferential. On held-out tissue, the student retains broad striatal organization, although recovery of the finer lesion-specific dopamine contrast is more limited (Supplemental Section S6; Supplemental Figs. S7, S8).

#### The dopamine component generalizes across spatial and dissociated multiomic targets

Having established that spatial distillation organizes dopamine in the source representation, we next asked whether the dopamine-associated program itself transfers to targets that were never metabolomically profiled. We evaluated this transfer in the two RNA–ATAC datasets introduced above: the spatial P22 dataset and the non-spatial 10x Multiome dataset. To obtain a common set of molecular components across these targets, we first defined shared components on the spatial RNA–MSI source using the leading singular vectors of the correlations between the student’s RNA-derived latent dimensions and measured metabolites. This decomposition used source data only, without selecting components based on their association with dopamine (Methods, *Shared PLS components*). We then evaluated the same components in both targets, using the trained teacher for P22 and the graph-masked student for Multiome, which lacks the spatial coordinates required by the teacher. Neither encoder was fitted on the target datasets. Gene and ATAC peak weights were computed as correlations with the component scores in each target, whereas metabolite weights were estimated through optimal-transport pairing with the source spots (Methods, *Metabolite weights in targets without MSI*).

Of the six components, C2 carries the largest dopamine weight in the source (0.63) and is, by construction, the same dopamine component in both targets. Mapped back spatially into tissue, the teacher component concentrates on the striatum (Fig. 4g; teacher-imputed dopamine in Supplemental Fig. S9). The student component, displayed over the mapped coordinates of the Multiome target, recovers the same striatal domain (Fig. 4h). Its weights recover striatal MSN biology together with dopamine and its metabolite 3-methoxytyramine (3-MT): the strongest gene weights, all positive, are *Rgs9*, *Rarb*, *Adcy5*, *Gpr88*, *Ryr3*, *Ppp1r1b*, *Gprin3*, *Gng7*, *Gnal*, and *Pde1b*, and the top recovered metabolite weights pair 3-MT and dopamine with a second dopamine annotation and seven unannotated *m/z* channels, one of them weighing negatively (Fig. 4i).

We then compared the teacher and student C2 gene weights with the NMF-derived reference signature from Fig. 4c (Supplemental Methods, *Convergence of NMF and PLS dopamine programs*). The 50 genes most specific to the NMF program carry higher C2 weights than the remaining shared genes, with a rank-based AUROC of 0.81 for the teacher (P22 spatial) and 0.81 for the student (dissociated Multiome) (Fig. 4j). The two top-50 sets share 8 genes for the teacher and 6 for the student (1.3 expected), including the striatal genes *Pde1b*, *Pcp4*, *Arpp21*, *Tac1*, and *Foxp2*.

Together, the NMF program independently associated with measured dopamine and the source-defined C2 component recover a concordant striatal transcriptional signature across the spatial RNA–MSI source, spatial P22 RNA–ATAC, and dissociated 10x Multiome datasets. An RNA-only latent yields a similar dopamine-associated component (Supplemental Fig. S10), indicating that this program is anchored in striatal gene expression, while source metabolomics provides the association used to identify and transfer the dopamine component. Applied to 5,000 random latent directions, the optimal-transport procedure used for target metabolite weights reproduced each direction’s source metabolite weights, and this agreement disappeared when the target latent was randomly rotated before pairing (Supplemental Fig. S11).

The dopamine-associated C2 component also carries a regulatory signal in the dissociated Multiome target. Its positively weighted chromatin peaks were enriched for D2-MSN cis-regulatory elements (NES = 3.18, FDR *<* 0.001), with leading-edge MEIS, PBX, and AP-1 motifs and RNA–ATAC-concordant support for EGR1 (Supplemental Section S7.5; Supplemental Figs. S12, S13). When motif accessibility was mapped onto the spatial RNA–MSI source through the student representation, these signals concentrated in the striatum (AUROC 0.68–0.82, Supplemental Fig. S14).

Together, these findings show that measured metabolites in the spatial source can anchor the interpretation of striatal transcriptional and cis-regulatory programs recovered in RNA–ATAC targets, even when those targets lack metabolomic measurements.

### 2.5 Graph-free transfer recovers striatal organization in human single-nucleus RNA-seq

Having established transfer of a dopamine-associated program across mouse spatial and dissociated multiomic datasets, we next asked whether a spatial transcriptome–metabolome representation could be transferred to a large human single-nucleus atlas in which metabolomics was not measured. We therefore trained SpatialJEPA with a SpatialMETA backbone on spatial RNA–MSI data from human PD striatum (Vicari et al. 2024) and applied the trained SpatialJEPA student to SEAAD caudate-head (CaH) snRNA-seq (Hawrylycz et al. 2024; Kana et al. 2026), embedding 714,534 nuclei from 42 donors (Methods, *Human transfer to SEA-AD caudate head*). Because the CaH is itself part of the striatum, this provides an anatomically matched but disease- and assay-distinct setting in which to test graph-free transfer of the human spatial RNA–MSI representation.

We defined five shared components from the human spatial RNA–MSI source and focused on C1, the component with the largest dopamine weight (Fig. 5a; Methods, *Shared PLS components*). To account for differences in metabolite abundance among the three source sections, we centered source RNA, MSI, and latent representations within each section before calculating feature correlations and defining the components. For transfer to CaH nuclei, we separately paired the nuclei with each source section through optimal transport and averaged the resulting metabolite estimates across sections. In a leave-one-section-out validation, C1 metabolite weights recovered by optimal transport from the other two source sections agreed with weights calculated from measured MSI in the held-out section (*r* = 0.69–0.90; Supplemental Fig. S15a; Supplemental Section S8). C1 combines positive recovered metabolite weights, including dopamine and serotonin, with positive RNA weights for canonical striatal genes including *ANO3*, *PDE10A*, and *GRM1*, together with the neuronal genes *SORCS3* and *CDH8*. Conversely, *TMEM144*, *DOCK5*, *MBP*, *QKI*, and *CLMN* are among the strongest negative RNA weights, including the glial-associated genes *MBP* and *QKI* (Fig. 5a). A second component, C2, separates astrocytes and ependymal cells from other subclasses and has similarly recoverable metabolite weights (Supplemental Figs. S16, S17).

**Figure 5:**
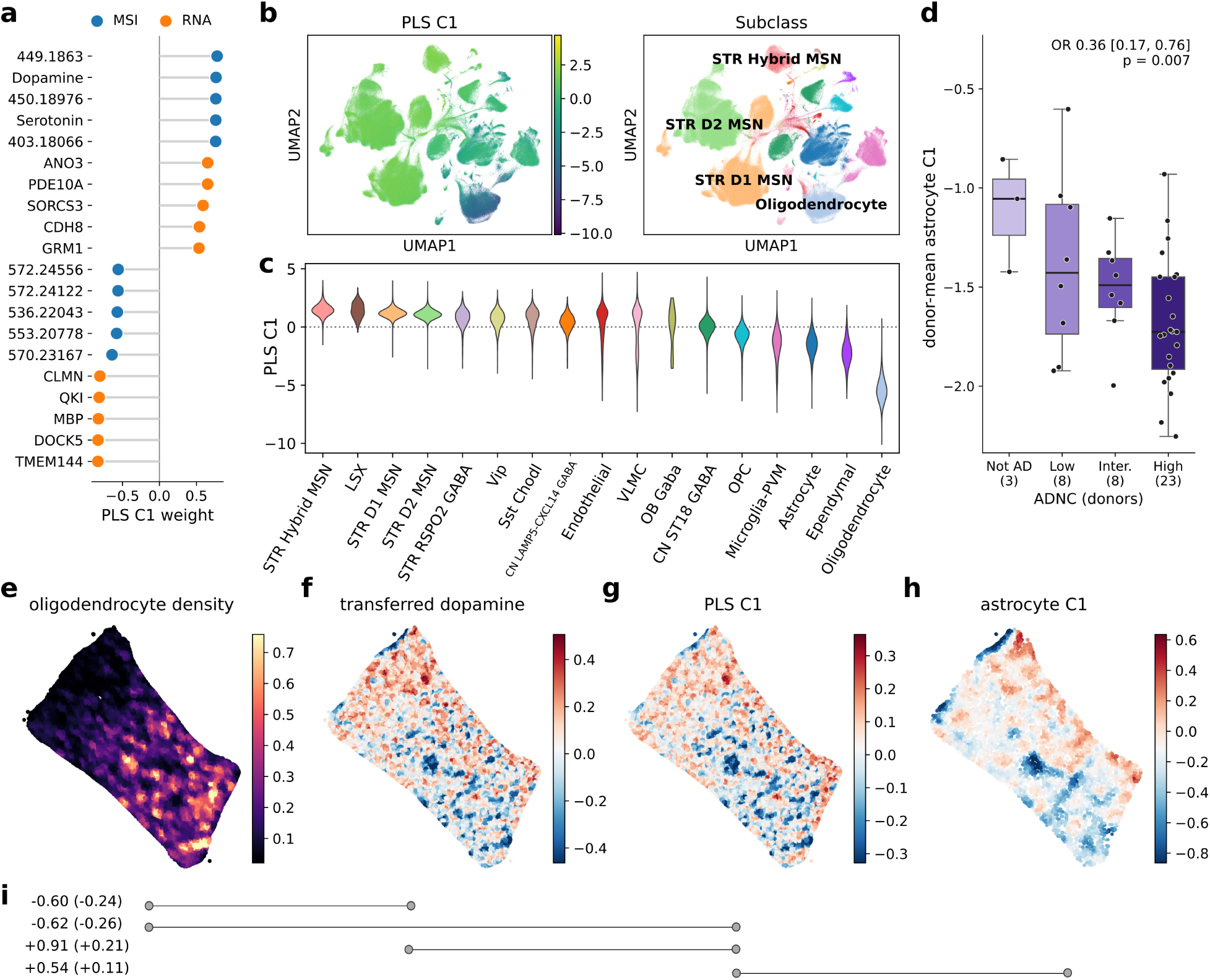
Graph-free transfer of a spatial transcriptome–metabolome representation to human caudate-head single-nucleus RNA-seq. **(a)** Top positive and negative weights of the dopamine-associated PLS component C1 across RNA and metabolites after transfer to SEA-AD caudate-head snRNA-seq; metabolite weights are recovered in the nuclei by optimal-transport pairing with the source spots. **(b)** RNA UMAPs colored by C1 score (left) and cell subclass (right); labels highlight striatal medium spiny neuron (MSN) and oligodendrocyte populations. **(c)** Distribution of C1 scores across cell subclasses, ordered by descending mean. **(d)** Donor-mean astrocyte C1 by AD neuropathologic change (ADNC; donors per category in parentheses), with its covariate-adjusted odds ratio per standard deviation. **(e)** Local oligodendrocyte density in a SEA-AD caudate-head Xenium section, used as a spatial proxy for white-matter-rich tissue. **(f)** Dopamine transferred to the snRNA-seq nuclei and carried to the Xenium cells through shared genes, averaged over 30 nearest spatial neighbors. **(g)** RNA-projected C1 scores, averaged over the same spatial neighborhoods. **(h)** C1 within astrocytes, carried from the snRNA-seq astrocytes through the Xenium panel genes and averaged over the 30 nearest astrocytes. **(i)** Pairwise bivariate Moran’s *I* among (e–g), and between (g) and (h) over the astrocytes; smoothed (unsmoothed) values.

Consistent with the C1 weights, the RNA-derived UMAP shows that high C1 scores concentrate within the region occupied by striatal medium spiny neurons (MSNs), whereas the oligodendrocyte population occupies the opposite, C1-negative end of the embedding (Fig. 5b). This organization is also evident when C1 is examined directly across annotated cell subclasses. Hybrid MSNs and LSX have the highest C1 distributions, closely followed by D1 and D2 MSNs; other neuronal subclasses take intermediate scores, while microglia, astrocytes, ependymal cells, and oligodendrocytes show the most strongly negative scores, with oligodendrocytes well below all other subclasses (Fig. 5c). C1 therefore defines a prominent MSN–oligodendrocyte axis after transfer to dissociated human nuclei. This axis is anatomically consistent with the organization of the human caudate, in which MSN-rich gray matter contrasts with oligodendrocyte-dense white-matter tracts (Kana et al. 2026).

We next examined associations between C1 and neuropathology separately within cell sub-classes: the three MSN subclasses at the positive end of C1 and the three most abundant glial subclasses (astrocytes, oligodendrocytes, and microglia), each tested against Alzheimer’s disease neuropathologic change (ADNC) (Kana et al. 2026), three continuous pseudo-progression scores of caudate pathology, and dementia (Methods, *C1 within subclasses and AD neuropathology*). The strongest association was in astrocytes, where donors with more advanced ADNC had lower mean C1 (odds ratio 0.36 per standard deviation, 95% confidence interval 0.17–0.76, *p* = 0.007; 42 donors; Fig. 5d); the three MSN subclasses showed weaker associations in the same direction (odds ratios 0.51–0.55, *p* = 0.06–0.10). This association is of particular interest given previous evidence that protoplasmic astrocyte abundance increases with tau pathology in the human caudate (Kana et al. 2026). Although the astrocyte association was nominally significant, it did not survive correction for multiple testing across the 30 tests (false discovery rate 0.22).

We next asked whether the transferred quantities underlying this axis could also be represented in spatially resolved human caudate tissue. In a SEA-AD CaH Xenium section, we computed local oligodendrocyte density as a proxy for white-matter-rich tissue, providing a spatial reference for the cellular organization observed in the snRNA-seq analysis (Fig. 5e). We then transferred the dopamine estimates from the snRNA-seq nuclei to Xenium cells based on the expression of genes shared between the two assays (Fig. 5f), and projected each cell onto C1 using its RNA weights (Fig. 5g). Both transferred quantities are shown after averaging each cell with its 30 nearest spatial neighbors. Transferred dopamine and projected C1 scores were positively associated spatially, as expected given their shared RNA-based transfer. Both showed negative spatial associations with local oligodendrocyte density, consistent with their enrichment in MSN-rich rather than oligodendrocyte-rich regions (Fig. 5i). Motivated by the association between astrocyte C1 and ADNC (Fig. 5d), we examined the spatial distribution of C1 among Xenium astrocytes. The RNA-weight projection used in Fig. 5g separates cell types but poorly captures C1 variation within astrocytes (donor-mean correlation *r* = 0.15 in the snRNA-seq astrocytes). We therefore fitted a ridge regression to predict C1 from the 464 Xenium panel genes using snRNA-seq astrocytes (*r* = 0.94 for the means of held-out donors), then applied it to Xenium astrocytes (Fig. 5h; Supplemental Methods, *Astrocyte C1 on the Xenium section*). The resulting astrocyte C1 map was spatially associated with projected C1 (bivariate Moran’s *I* = 0.54; 0.11 unsmoothed; Fig. 5i) and was lowest in oligodendrocyte-dense tissue and along the ventricular wall.

Taken together, these results show that a spatial transcriptome–metabolome representation learned in human striatum can be transferred to large non-spatial single-nucleus datasets while retaining biologically interpretable striatal organization. In SEA-AD CaH, the transferred representation resolves a coherent MSN–oligodendrocyte axis and carries a corresponding metabolic signal into nuclei for which no metabolomic measurements were available. More broadly, this establishes a route for using spatial multiomic datasets as privileged references to augment large dissociated cell atlases with tissue- and metabolism-informed latent structure, and to project those transferred representations back into independently profiled spatial tissue when spatial transcriptomic data are available.

## 3 Discussion

Single-cell and spatial multiomics provide complementary views of cellular state, but most large-scale single-cell atlases lack the tissue context that shapes molecular organization. Here, we introduce SpatialJEPA, a framework that uses spatial multiomics as training-time supervision to learn molecular representations that operate without coordinates. Across spatial RNA–ATAC and transcriptome–metabolome settings, graph-context distillation preserves aspects of tissue organization and multimodal molecular structure, including when applied to independent dissociated nuclei and tissue sections from held-out animals. In particular, the recovery of spatially informed representations in previously unseen nuclei supports the transfer of learned tissue context beyond the training data. We further show that measured spatial metabolites can anchor the interpretation of transcriptional and cis-regulatory programs in datasets lacking metabolomic measurements. Applied to human caudate-head nuclei, the transferred representation recovers an interpretable MSN–oligodendrocyte axis with dopamine associations derived from the spatial source. Together, these findings demonstrate how spatial multiomics can provide supervision for multimodal representations of cellular state, extending its role beyond that of a reference for downstream spatial mapping.

Spatial information can refine what is learned about cellular state because tissue organization, cellular neighborhoods, and intercellular interactions provide biological constraints that are not observable from dissociated molecular profiles alone (Jena et al. 2024). Recent work supports this broader view of spatial context as a source of representation-level information: spatially informed transcriptomic models can generalize to dissociated single-cell measurements (Sun et al. 2024), while ST-JEPA shows that JEPA-style latent prediction can exploit spatial neighborhood structure to learn biologically informative tissue representations (Birk et al. 2026b). Together with our results, these studies suggest that spatial context can serve not only as a property to be reconstructed, but also as an additional source of supervision for learning molecular representations of cellular state.

Spatial context may also provide a common organizing layer for integrating complementary molecular measurements that are not jointly observed. SpatialEx, for example, uses spatially registered histology as a common anchor for predicting and diagonally integrating distinct omics measurements across tissue sections (Liu et al. 2025). More generally, this suggests that tissue context can provide a shared reference through which heterogeneous molecular modalities are related within a common representation.

These principles may be relevant to future AI Virtual Cells and biological world models, which aim to integrate information across molecular modalities and biological scales into representations of cellular state (Bunne et al. 2024; Noutahi et al. 2025; Chen et al. 2026). Recent formulations of Virtual Cell world models further emphasize generative latent prediction, in which state evolution is modeled in latent space while generative decoders ground those states in observable, multimodal biological outcomes (Xing et al. 2026). This provides a complementary extension of the latent-prediction principle underlying SpatialJEPA: spatial context could help shape a richer representation of cellular state, while generative components could ultimately relate such representations back to molecular, structural, or other measurable phenotypes. SpatialJEPA contributes to this direction at the level of state representation rather than dynamics or generation; it does not model perturbation-conditioned transitions or constitute a complete biological world model. Instead, our results suggest one way in which tissue context could inform the latent state representations upon which richer Virtual Cell models are built.

Several limitations qualify these findings. First, spatial placement requires alignment to a biologically comparable source rather than direct coordinate prediction. The held-out Multiome benchmark uses graph-informed teacher domains to assess transfer of spatial organization more directly than the published modality-specific P22 clusters, although agreement with these domains primarily measures preservation of teacher-learned structure. Because dissociated Multiome nuclei lack independently known tissue coordinates, this benchmark cannot establish the accuracy of their anatomical placement; direct validation would require target molecular profiles linked to measured spatial locations. The separate position-recovery benchmark also uses source spots seen during training (Supplemental Section S2). Second, transfer of spatial structure is not uniform. In held-out RNA–MSI sections, broad striatal organization was preserved, but the finer lesion-associated dopamine contrast transferred less consistently (Supplemental Section S6). Student–teacher neighborhood agreement also varies across tissue domains, with larger differences in contiguous regions such as white matter and striatum (Supplemental Fig. S3d). Third, metabolites were not directly measured in the target RNA–ATAC or human caudate-head nuclei. Their associated weights were estimated through pairing with measured source profiles and therefore represent source-referenced associations rather than direct metabolic measurements in the targets. Finally, the caudate head exhibits relatively selective AD-associated changes compared with affected cortical regions (Kana et al. 2026), and the observed astrocyte C1 association with AD neuropathologic change did not survive correction for multiple testing. Further evaluation across independent tissues, disease contexts, and matched target metabolomics will be needed to establish the generality of these findings.

In conclusion, SpatialJEPA demonstrates that spatial graph context can be distilled into transferable latent representations, enabling tissue-informed analyses of cells and nuclei that lack spatial coordinates. By transferring aspects of tissue organization to dissociated profiles and relating their transcriptional and regulatory programs to metabolites measured in spatial sources, the framework extends spatial multiomics beyond reference mapping to representation-level supervision. This offers a path toward spatially grounded models of cellular state that can be applied to large, non-spatial single-cell atlases.

## 4 Methods

### 4.1 Datasets and preprocessing

#### Spatial RNA–ATAC source

The spatial RNA–ATAC source dataset is the postnatal day 22 (P22) mouse brain dataset of Zhang et al. (2023) (NCBI Gene Expression Omnibus [GEO] accession GSE205055). It jointly profiles chromatin accessibility and gene expression while preserving spot coordinates across a coronal section, and provides ‘RNA_cluster’ and ‘ATAC_cluster’ annotations used as reference biological cluster labels.

#### Dissociated RNA–ATAC target

The non-spatial target dataset is an RNA–ATAC 10x Genomics mouse brain Multiome sample (10x Genomics 2023) with paired RNA and ATAC profiles, but no spatial coordinates. It was obtained from the 10x Genomics dataset repository (https://www.10xgenomics.com/datasets/mouse-brain-nuclei-isolated-with-chromium-nuclei-isolation-kit-saltyez-protocol-and-10x-complex-tissue-dp-ct-sorted-and-ct-unsorted-1-standard). This sample carries no author-provided cell-type annotation, so CellTypist (Domínguez Conde et al. 2022) was applied to the target RNA view using the pre-trained Mouse_Whole_Brain model. These predicted labels were used only for evaluation and were not provided to any model during training.

#### Spatial transcriptome–metabolome data

Spatial RNA–MSI data combine 10x Visium spatial transcriptomics with MALDI mass spectrometry imaging (MSI) in mouse and human brain tissue (Vicari et al. 2024). Processed spatial transcriptomics and MSI data were obtained from Mendeley Data (doi:10.17632/w7nw4km7xd.1); the spatial transcriptomics data are also deposited in GEO (GSE232910) and raw MALDI-MSI data in SciLifeLab Figshare (doi:10.17044/scilifelab.22770161).

Mouse analyses used section V11L12-109_B1 (animal mPD3), from a unilateral 6-hydroxydopamine (6-OHDA) Parkinson’s disease model in which dopaminergic input is depleted in one striatal hemisphere; per-spot annotations distinguish striatal from non-striatal tissue. The human SpatialMETA model used for SEA-AD transfer was trained on three sections from a single Parkinson’s disease donor (V11T17-102_A1, _C1 and _D1; donor hPD1), which were horizontally integrated as described in Supplemental Methods (Section S9.6).

Reprocessing of these sections and co-registration of their MSI and Visium measurements are described in Supplemental Methods (Section S9.1).

#### Human cohorts

SEA-AD caudate head (CaH) single-nucleus RNA-seq and Xenium data were obtained from the SEA-AD resource (Hawrylycz et al. 2024; Kana et al. 2026) through the AWS Open Data Registry (s3://sea-ad-single-cell-profiling/Caudate_Nucleus/ and s3://sea-ad-spatial-transcriptomics/caudate_nucleus/). The CaH cohort comprises 42 donors and 714,534 nuclei with harmonized cell-subclass annotations.

#### RNA–ATAC feature harmonization and graph construction

Source and target features were placed on a common index before training, with genes matched by symbol and chromatin-accessibility peaks matched by genomic interval overlap. A candidate gene–peak graph was constructed by linking each gene to accessible peaks within 150 kb of its promoter. Candidate edges were assigned the MultiGATE distance-decay prior (Miao et al. 2025). Genes were restricted to those identified as variable by SCTransform in both source and target datasets, and peaks to those linked *in-cis* to a retained gene, yielding identical RNA and ATAC feature spaces across domains. Consequently, downstream gene–peak analyses operate on this restricted shared feature set rather than on the full transcriptome. For spatial samples, the tissue graph connected spots within a radius of 40 in the array coordinate frame supplied with the dataset; graph-masked and dissociated inputs used self-only connectivity instead.

#### RNA–MSI preprocessing and feature selection

Following spatial alignment, RNA and MSI measurements were represented as paired modalities over the same set of Visium spots. To ensure compatibility with downstream transfer datasets, the RNA candidate feature space was restricted using target-derived rankings. Within each candidate pool, spatially variable RNA and MSI features were selected using univariate Moran’s *I*. This yielded 1,806 RNA genes and 800 MSI features for the mouse model, and 1,057 RNA genes and 790 MSI features for the human model. Thus, target data informed feature selection, but encoder parameters were trained exclusively on spatial source data. Further annotation and preprocessing details are given in Supplemental Methods (Section S9.2).

### 4.2 SpatialJEPA model and training

#### Graph-context distillation

SpatialJEPA augments a spatial multiomic backbone with a teacher–student objective in which teacher and student receive the same molecular profiles but different graphs (Fig. 1a): the full spatial graph *G_s_* for the teacher and a self-only identity graph *G*_id_ for the student,

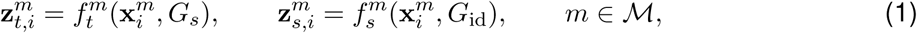

where *i* indexes samples (cells, nuclei, or spots) and *M* is the backbone-specific set of distilled representations. The teacher, with its own parameters, is optimized under the backbone objective, and the student is trained to predict its latent representations with a stop-gradient mean-squared-error objective,

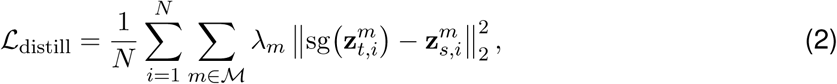

where sg(*·*) is the stop-gradient operator and *λ_m_* weights each representation. Training alternates: at each iteration, the teacher is updated with the spatial graph, and its latents are then held fixed as targets for a student update with the identity graph (Supplemental Algorithm S1). The framework therefore requires only that the backbone can operate with either graph, and the trained student needs no coordinates at inference.

#### MultiGATE backbone

For spatial RNA–ATAC data, the backbone is MultiGATE (Miao et al. 2025) (Fig. 1b), a graph-attention autoencoder that encodes RNA and ATAC separately using a spatial spot–spot graph and a gene–peak graph, reconstructs each modality, and aligns them with a symmetric contrastive objective on matched profiles (full objective and training settings in Supplemental Methods, Section S9.4). The distilled set comprised the projected, *ℓ*_2_-normalized RNA and ATAC embeddings used in the contrastive objective, *M* = *{***f**, **g***}*, with *λ_m_* = 1; training was full-batch, with one student step per teacher step. A non-spatial MultiGATE ablation was trained on the same profiles with the same objective and schedule, but with the identity graph and no teacher supervision, matching the student’s backbone and inference-time graph without spatial information.

#### SpatialMETA backbone

For spatial transcriptome–metabolome data, the backbone is Spatial-META (Tian et al. 2025) (Fig. 1b), a conditional variational autoencoder for cross-modal and cross-sample integration of spatial transcriptomics (ST) and spatial metabolomics (SM). The architecture and full set of equations are given in Supplemental Methods (Section S9.5).

#### Spatial and cross-modal extensions of SpatialMETA

SpatialMETA’s original encoder ignores spatial neighborhoods. To enable distillation, we replaced the fully connected encoder layer of each modality with a graph-convolution layer (Kipf and Welling 2017), leaving the rest of the model unchanged; the teacher graph connected each spot to its six nearest spatial neighbors (symmetrized), and the student’s identity graph reduces the convolution to a per-spot linear map. We also added to the teacher’s backbone an ST*→*SM cross-reconstruction term, *L^st→sm^_recon_*, which applies the SM decoder to the ST-specific latent mean ***µ***_st_, training the RNA-derived latent to retain metabolite information without new parameters (Supplemental Methods, Section S9.5). Original loss weights were retained for the existing terms (Supplemental Table S1).

#### SpatialMETA distillation and training

Teacher and student used this modified backbone on the same RNA–MSI profiles; the non-spatial ablation used the unmodified SpatialMETA on the same data and schedule, without graph encoders, teacher supervision, or the ST*→*SM term. The distilled set comprised the joint posterior mean ***µ***_joint_ (*λ* = 1) and the ST- and SM-specific means ***µ***_st_ and ***µ***_sm_ (*λ* = 0.5 each), each *z*-scored per dimension within the mini-batch before Eq. 2. Optimization, training schedules, and teacher early stopping are described in Supplemental Methods (Section S9.6).

### 4.3 Zero-shot transfer and post-hoc decoding

#### Zero-shot transfer and optimal-transport alignment

After source training, the student is applied to a non-spatial target without retraining, using the identity graph. Here, “zero-shot” means that the encoder is neither trained nor fine-tuned on target data: target data informed only the shared input feature space, for both backbones (*RNA–ATAC feature harmonization and graph construction*; *RNA–MSI preprocessing and feature selection*), and the post-hoc steps (OT alignment, clustering, and PLS decoders) are fitted on the target. For MultiGATE, source and target features are harmonized before training (*RNA–ATAC feature harmonization and graph construction*). For SpatialMETA, only the target’s RNA is used, including for the RNA–ATAC targets; target genes are matched to the source model, and missing genes and the absent MSI input are set to zero. To place an RNA–ATAC target within the source reference, target and source embeddings were aligned by entropic OT (Peyré and Cuturi 2020; Gossi et al. 2023; Damodaran et al. 2018), separately for RNA and ATAC (Euclidean cost, *ε* = 0.001, balanced uniform marginals). Given the coupling ***γ*** *∈* R*^NT ×NS^* between target embeddings *Z^T^_i_* and source embeddings *Z^S^_j_*, each target embedding was mapped by barycentric projection,

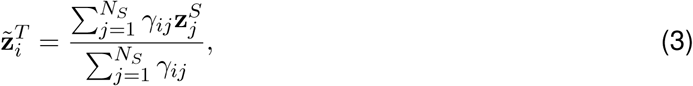

with uniform marginals *Σ^NS^_j=1_ γ_ij_ = 1/N_T_* for each target profile *i* and *Σ^NS^_j=1_ γ_ij_ = 1/N_S_* for each source profile *j* (balanced OT). Equivalently, Eq. 3 can be written in matrix form as **Z**^~^*^T^* = *N_T_ **γ*****Z***^S^*. Each transported target profile was then assigned the spatial coordinates of its nearest source reference profile in the projected latent space. This assignment is post-hoc retrieval rather than coordinate prediction, and OT is not part of the training objective.

#### Post-hoc PLS decoders

The *post-hoc* PLS decoders are linear models fitted after training, distinct from the backbones’ neural decoders. In the RNA–ATAC analyses, separate PLS models are fitted in source and target for interpretation; for RNA–MSI transfer, a decoder is fitted on each target from its latent to source metabolite profiles paired by OT (*Metabolite weights in targets without MSI*). All were *K*-component PLS regressions (Tsuchiya et al. 2022) (scikit-learn PLSRegression, or an equivalent SIMPLS implementation for metabolites). For column-standardized embeddings **X**_0_ *∈* R*^N×D^* and features **Y**_0_ *∈* R*^N×F^*, PLS extracts *K* components of maximal predictor–response covariance, with rotation **R** *∈* R*^D×K^* and response loadings **Q** *∈* R*^F^ ^×K^*,

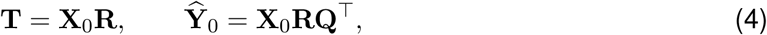

where **T** holds the component scores and **Q** the associated features (derivation in Supplemental Methods, Section S9.7, and Supplemental Fig. S18; analysis-specific settings in *PLS decoding of RNA–ATAC embeddings* and *Metabolite weights in targets without MSI*).

### 4.4 Representation benchmarks

Baselines were the non-spatial ablations described above, MultiVI (Ashuach et al. 2023) trained on the source RNA–ATAC counts and applied zero-shot to the target, and Multigrate (Litinetskaya et al. 2025) trained on the mouse RNA–MSI section. Cross-modal pairing, modality mixing, and conservation of annotated structure were quantified by 1 *−* FOSCTTM, iLISI, and a biological-conservation score from scIB metrics (Luecken et al. 2022; Niu et al. 2025). Spatial-structure metrics were evaluated on the source with ten spatially compact blocks: niche recovery and orthogonal-target recovery are ridge-regression readouts, cross-validated over blocks and scored by Spearman correlation, of each spot’s neighborhood cluster composition from the joint embedding and of the neighbor-averaged ATAC profile from the RNA embedding, and within-cluster coherence is the Spearman correlation between embedding and physical distances within clusters. Held-out Multiome nuclei were coupled to P22 spots by 15-nearest-neighbor or OT coupling, and their Leiden clusters were compared with the teacher’s P22 Leiden domains by ARI. The dopamine benchmark measures, on *k* = 15 latent-neighbor graphs, the gain in intact–lesioned AUROC from neighbor smoothing of measured dopamine, its latent-graph Moran’s *I*, and the enrichment of intact striatum among dopamine-high neighborhoods, with intervals from a within-section spatial-block bootstrap. Full definitions are given in Supplemental Methods (Section S9.9).

### 4.5 Analysis of transferred RNA–ATAC representations

Leiden clusters (*k* = 150, resolution 0.9) were computed separately in source and OT-aligned target from the stacked RNA and ATAC student embeddings, and target clusters were matched one-to-one to source clusters by the Hungarian algorithm. Within matched clusters, genes differentially expressed in both domains (*t*-test, Benjamini–Hochberg *p <* 0.05) were ranked by *−* log *p*_adj_, and source and target ranks were compared by Spearman correlation. Nine-component PLS decoders (Eq. 4) were fitted separately in source and target, from the RNA or ATAC embedding to SCTransform residuals or accessibility; components were sign-aligned across domains, and their gene loadings were tested against six predefined marker modules relative to 2,000 size-matched random gene sets (Benjamini–Hochberg-corrected). Gene–peak links were taken from the student’s MultiGATE gene–peak attention, thresholded by a two-component Gaussian mixture, and each gene’s linked-peak accessibility was summarized as an attention-weighted sum of its peaks. Its spatial coupling with expression was measured by bivariate Moran’s *I* on a six-nearest-neighbor graph of the 8,945 source spots occupied by aligned target nuclei, after averaging target profiles per spot. Full details are given in Supplemental Methods (Section S9.13).

### 4.6 Dopamine-associated programs and human transfer

The reference program (Fig. 4c) was obtained by non-negative matrix factorization (Lee and Seung 1999) of the SCTransform-corrected counts of the mouse RNA–MSI section (ranks 1–29, generalized Kullback–Leibler objective), without using dopamine or any SpatialJEPA output; the component with the highest bivariate Moran’s *I* against measured dopamine on the Visium lattice was retained. Its convergence with the dopamine component (Fig. 4j) was assessed by comparing the C2 weights of the 50 genes most specific to this NMF component with those of the other shared genes (rank-based AUROC), and by a hypergeometric test of the overlap between the two top-50 gene sets. Full details are given in Supplemental Methods (Section S9.14).

#### Shared PLS components

Components were defined once on the source and read out in every dataset through the same latent. Let **Z** be the student’s ST-specific (RNA) latent mean ***µ***_st_ (*D* = 10), computed from RNA alone and *z*-scored within each dataset, and **C** *∈* R*^D×F^* the correlations between the source latent dimensions and the *F* measured, log(1 + *x*)-transformed MSI features. The components **u**_1_, …, **u***_K_* are the leading left singular vectors of **C***/ ‖* **C**| *_F_* (*K* = 6 for the mouse model, 5 for the human model), so neither dopamine nor target data enter the decomposition. Scores are **s***_k_* = **Zu***_k_*, and a feature’s weight is its Pearson correlation with **s***_k_* across the cells of its own dataset. Components were oriented so that source dopamine weighs positively; the dopamine component has the largest absolute dopamine weight (C2 mouse, C1 human). On P22, the teacher’s ***µ***_st_, computed with the P22 spatial graph, was read out instead. Mouse components were fitted on the 1,218 intact-hemisphere spots, because the lesion depletes dopamine without altering MSN identity, a decoupling absent from the targets (the lesion comparisons of Fig. 4d–f use all spots); for the human model, features and latents were centered within each source section.

#### Metabolite weights in targets without MSI

Target cells were paired with source spots by entropic OT between *z*-scored latents (mean-scaled squared-Euclidean cost, balanced marginals; regularization 0.01 for the mouse and 0.001 for the human model, chosen on held-out data), and each cell received the coupling-weighted mean of the paired spots’ standardized MSI (Eq. 3 with metabolite profiles in place of embeddings). A reduced-rank PLS decoder (Eq. 4) was fitted on the target from the latent to the paired MSI, with its rank chosen by five-fold cross-validation (one-standard-error rule). The weight of metabolite *m* on component *k* estimates the cross-loading corr(*m*~, **s***_k_*) (ter Braak 1990) as *R_m_* corr(*m*^, **s***_k_*), where *m*^ is the decoded metabolite and *R_m_* its cross-validated correlation with the paired values (zero if negative), an attenuation factor (Spearman 1904) that down-weights poorly decoded metabolites. For the human model, transfer was run per source section and averaged, using 40,000 subsampled CaH nuclei (Supplemental Fig. S19; random-direction null in Supplemental Fig. S11).

#### Human transfer to SEA-AD caudate head

The human RNA–MSI student was applied to SEA-AD CaH snRNA-seq as described under *Zero-shot transfer and optimal-transport alignment*, after matching each donor’s matrix to the source model’s genes, yielding embeddings of 714,534 nuclei from 42 donors. Five shared components were computed (*Shared PLS components*), with gene weights computed across the nuclei and metabolite weights recovered by section-wise OT pairing (*Metabolite weights in targets without MSI*) rather than from MSI measured in the nuclei. Subclass distributions were ordered by their mean C1 score.

On the SEA-AD CaH Xenium section H21.33.002.CX18.CaH.02.01 (48,717 cells), local oligodendrocyte density was estimated by Gaussian kernel density over oligodendrocyte coordinates (Fig. 5e). Dopamine was carried from the nuclei to Xenium cells by ridge regression on the 241 genes shared by the source model and the Xenium panel (Fig. 5f), C1 was projected through the snRNA-seq C1 weights of the same genes (Fig. 5g), and astrocyte C1 was predicted by ridge regression on the 464 Xenium panel genes, trained on snRNA-seq astrocytes (Fig. 5h). Maps were averaged over 30 spatial neighbors (30 nearest astrocytes for astrocyte C1), and their pairwise spatial associations were quantified by bivariate Moran’s *I* on a symmetrized six-nearest-neighbor graph (Fig. 5i). Full details are given in Supplemental Methods (Section S9.17).

#### C1 within subclasses and AD neuropathology

Six subclasses were tested: D1, D2, and hybrid MSNs, and astrocytes, oligodendrocytes, and microglia. Nucleus-level C1 was averaged per donor with at least 50 nuclei of the subclass (42 donors; 41 for hybrid MSNs) and related to ADNC (Not AD, Low, Intermediate, High) by cumulative-logit ordinal regression adjusting for age at death, sex, and post-mortem interval, with C1 and continuous covariates standardized (odds ratios per standard deviation of C1). The caudate continuous pseudo-progression score (CPS) from combined AT8 and 6E10 features and the AT8- and 6E10-specific CPSs (Kana et al. 2026) were modeled by ordinary least squares, and dementia by logistic regression; the 30 *p*-values were Benjamini–Hochberg-corrected.

### 4.7 Software availability

The SpatialJEPA source code is freely available for academic and nonprofit use under the Apache 2.0 license at GitHub (https://github.com/li-lab-mcgill/SpatialJEPA) and is provided as Supplemental Code.

Models were implemented in PyTorch (Paszke et al. 2019), with graph operations provided by PyTorch Geometric (Fey and Lenssen 2019). Single-cell and multimodal data processing used Scanpy (Wolf et al. 2018) and MuData/muon (Bredikhin et al. 2022), and spatial analyses used Squidpy (Palla et al. 2022). The SpatialMETA teacher, student, and non-spatial models were trained under PyTorch 2.2.2 (CUDA 12.1) with PyTorch Geometric 2.6.1, scikit-learn 1.5.0, anndata 0.10.8, MuData 0.2.4, and muon 0.1.6. The motif scan was run in a separate environment carrying the MOODS and JASPAR dependencies. Optimal-transport couplings for metabolite transfer were computed with POT (Flamary et al. 2021), and per-nucleus motif deviations with pychromVAR (Li 2023), a Python implementation of chromVAR (Schep et al. 2017) within the scverse ecosystem (Virshup et al. 2023). Analyses with distinct dependency requirements were executed in separate conda environments, with complete environment specifications provided alongside the code. Training configurations, losses, run identifiers, and model checkpoints were tracked with MLflow (Zaharia et al. 2018), and downstream analyses were linked to saved run identifiers and checkpoints to preserve the provenance of the model outputs used to generate the reported results.

## Data access

No new sequencing, imaging, or mass spectrometry data were generated in this study. All analyzed datasets are publicly available and were previously published; their sources and accession numbers are given in Methods (*Datasets and preprocessing*): spatial RNA–ATAC data from GEO (GSE205055); 10x Genomics Multiome data from the 10x Genomics dataset repository; spatial RNA–MSI data from Mendeley Data (doi:10.17632/w7nw4km7xd.1), GEO (GSE232910), and SciLifeLab Figshare (doi:10.17044/scilifelab.22770161); and SEA-AD caudate-head single-nucleus RNA-seq and Xenium data from the AWS Open Data Registry.

## Competing interest statement

D.M.K. is a founder of Cervolve Inc. The authors declare no other competing interests.

## Acknowledgments

Y.L. is supported by a Canada Research Chair (Tier 2) in Machine Learning for Genomics and Healthcare (CRC-2021-00547), a Natural Sciences and Engineering Research Council (NSERC) Discovery Grant (RGPIN-2026-07343), and a Canadian Institutes of Health Research (CIHR) Project Grant (PJT-540722). D.M.K. is supported by an NSERC Canada Graduate Scholarship – Doctoral (CGS D). A preliminary version of the spatial RNA–ATAC results was presented as a short paper at CIBB 2026.

## Author contributions

D.M.K. and Y.L. conceived and designed the study. D.M.K. developed and implemented the methods and analysis workflows, performed the analyses, interpreted the results, and wrote the manuscript. Y.L. supervised the project and revised the manuscript.

## Notes

### Summary of Updates

The manuscript has been substantially expanded to demonstrate the generalizability of SpatialJEPA across molecular modalities, computational architectures, species, and datasets. The original spatial RNA-ATAC analyses have been extended through additional benchmarking against non-spatial multiomics integration methods, quantitative assessments of spatial neighborhood recovery, and independent validation using dissociated single-cell multiomics data. These analyses further characterize the extent to which spatial-context distillation preserves tissue organization in molecular representations generated without spatial coordinates. Downstream biological analyses have also been expanded to systematically quantify changes in gene expression and chromatin accessibility coupling following alignment to a spatial reference. SpatialJEPA has been generalized to spatial transcriptomic and metabolomic data through a graph-augmented implementation of SpatialMETA. Using data from a mouse model of Parkinson's disease, new analyses investigate the preservation of metabolically relevant tissue organization and generalization to tissue sections from held-out animals. Further experiments demonstrate how metabolite-associated molecular programs learned from spatial transcriptomic and metabolomic data can be transferred to independent spatial and dissociated RNA-ATAC datasets. These analyses identify transcriptional signatures and cis-regulatory elements associated with a dopamine-related molecular program in the striatum. The revised manuscript also introduces an application to human brain data, whereby a model trained on spatial transcriptomic and metabolomic profiles from the human striatum is applied to over 700,000 nuclei from 42 donors in the SEA-AD atlas. This analysis investigates metabolite-associated transcriptional programs across cell types and their potential relationships with Alzheimer's disease neuropathology. Additional validation experiments assess the reliability of recovering metabolite associations in the absence of directly measured metabolomic profiles in target datasets. The Introduction and Discussion have been revised to reflect the broader methodological scope of SpatialJEPA and its potential applications to the integration of spatial and non-spatial multiomics datasets. The Methods and Supplemental Material have been substantially expanded to document the additional computational architectures, benchmarking experiments, biological analyses, and robustness assessments. Figures have been revised and new figures introduced to present these results.

