## Supplementary Materials for "SpatialJEPA: Graph-context distillation enables spatially grounded single-cell representations across molecular assays"

Dylan Mann-Krzisnik Yue Li

#### Contents

**S1 Integration metrics on the P22 spatial RNA–ATAC source and dissociated target 3**

**S2 Graph-context distillation recovers relative spatial positioning without coordinate inputs 6**

**S3 Position recovery on RNA-side and ATAC-side latents 8**

**S4 Held-out Multiome placement scored against the published annotations 10**

**S5 RNA and ATAC PLS components on the P22 source and the aligned target 10**

**S6 Teacher early stopping and held-out evaluation of the mouse RNA–MSI model 10**

### S1 Integration metrics on the P22 spatial RNA–ATAC source and dissociated target

The main text evaluates graph-context distillation on the P22 mouse brain spatial RNA–ATAC dataset using metrics of spatial structure preservation. Here, we report complementary integration metrics for the same models: 1-FOSCTTM for cross-modal pairing, iLISI for RNA–ATAC modality mixing, and a biological conservation score for preservation of annotated structure (Fig. S1). Higher values indicate better performance for all three metrics.

On the spatial source (Fig. S1a), the `SpatialJEPA` teacher and graph-masked student achieve similar values across all three metrics, indicating that cross-modal integration is largely preserved after graph-context masking. Both outperform the independently trained non-spatial MultiGATE ablation. MultiVI performs similarly to the `SpatialJEPA` models on 1-FOSCTTM and achieves the highest iLISI, while the `SpatialJEPA` models retain higher biological conservation. The source UMAPs of the same representations (Fig. S1c,d) are consistent with these scores: the non-spatial ablation largely separates RNA and ATAC profiles on the source, whereas the teacher and student interleave them and share a similar layout of RNA clusters. On the dissociated target (Fig. S1b), the zero-shot student achieves the highest 1-FOSCTTM and biological conservation, while the OT-aligned student and source-trained MultiVI show stronger modality mixing.

**Cross-modal pairing.** Integration metrics were computed on the modality-specific latent embeddings using the single-cell integration benchmarking (scIB) metrics implementation where applicable (Luecken et al. 2022; Niu et al. 2025). Let $\mathbf{z}_i^R$ and $\mathbf{z}_i^A$ denote the RNA and ATAC embeddings of paired observation $i$, corresponding to a spatial spot in the source or a nucleus in the dissociated target. Cross-modal distances were computed directly in the model-produced embedding spaces using Euclidean distance,

$$D_{ij} = \left\| \mathbf{z}_i^R - \mathbf{z}_j^A \right\|_2 .$$

The fraction of samples closer than the true match (FOSCTTM) is the mean, over observations and both modality directions, of the fraction of cross-modal candidates lying closer than the paired observation:

$$\text{FOSCTTM} = \frac{1}{2N} \sum_{i=1}^{N} \left[ \frac{1}{N} \sum_{j \neq i} \mathbb{I}(D_{ij} < D_{ii}) + \frac{1}{N} \sum_{j \neq i} \mathbb{I}(D_{ji} < D_{ii}) \right] . \tag{S1}$$

We report $1 - \text{FOSCTTM}$ so that larger values indicate more accurate cross-modal pairing.

**Modality mixing.** Modality mixing was measured using the integration local inverse Simpson's index (iLISI) after pooling the RNA and ATAC embeddings and treating modality of origin as the batch label. The scIB Metrics implementation constructs a 90-nearest-neighbor graph for the LISI metrics; with the default settings, this corresponds to a perplexity of 30. For each observation,

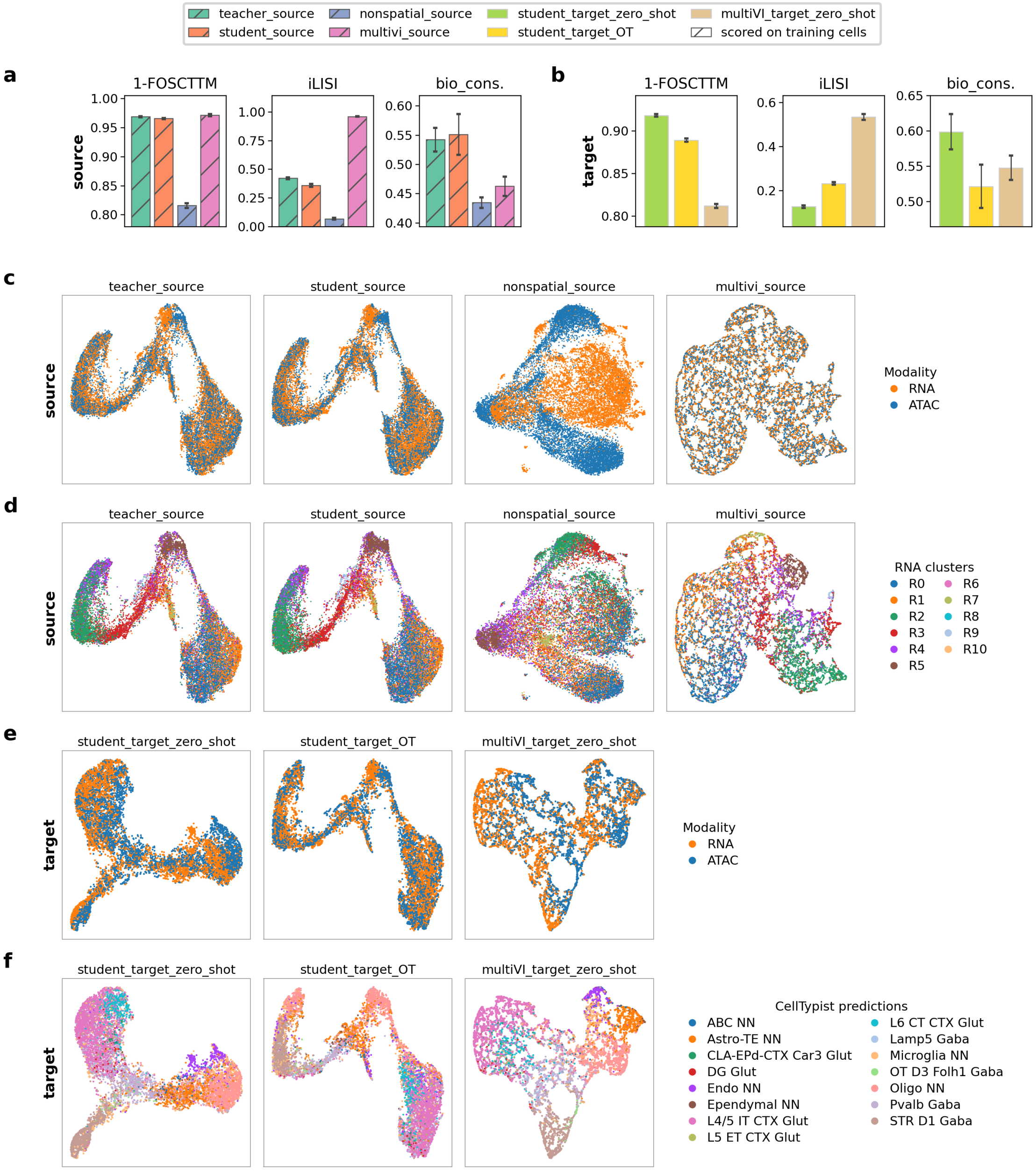

**Figure S1: Standard integration metrics and embeddings for graph-context distillation on the P22 mouse brain spatial RNA–ATAC source and the dissociated 10x Multiome target.** **(a)** Metrics on the spatial source, for the `SpatialJEPA` teacher, the graph-masked student, the non-spatial MultiGATE ablation, and MultiVI. **(b)** Metrics on the dissociated target, for the zero-shot student, the OT-aligned student, and source-trained MultiVI applied zero-shot. Columns are, from left to right, 1-FOSCTTM, iLISI, and a biological conservation score; higher is better for all three and axes are independently scaled. Hatched bars denote evaluation on training cells. Label-dependent metrics use one annotation for both modalities (RNA clusters for the source, CellTypist predictions for the target) and should be compared within a panel. Error bars are 95% intervals, obtained by bootstrap of per-observation scores for 1-FOSCTTM and by repeated subsampling for the other metrics (see *Uncertainty estimation* below). **(c,d)** UMAPs of each model's pooled RNA and ATAC embeddings on the source, colored by modality **(c)** and by the RNA clusters used for the label-dependent metrics **(d)**. **(e,f)** The same for the target, colored by modality **(e)** and by CellTypist predictions **(f)**. Points are drawn in random order. Target UMAPs show the same random 20% of nuclei as Fig. 2d, so the zero-shot student panels reproduce that figure. Source UMAPs use different settings from Fig. 2b,c and therefore differ in layout.

distance-weighted local modality proportions $p_m$ define the local inverse Simpson index

$$\text{LISI} = \left( \sum_m p_m^2 \right)^{-1}. \tag{S2}$$

The median local value is then scaled as

$$\text{iLISI} = \frac{\text{median}(\text{LISI}) - 1}{M - 1}, \tag{S3}$$

where $M = 2$ modalities, yielding a score from 0 for locally separated modalities to 1 for maximal local mixing.

**Shared biological annotation.** All label-dependent metrics were computed on pooled RNA and ATAC embeddings using a shared biological annotation for each paired observation. Using separate RNA and ATAC cluster labels would confound biological conservation with modality identity. We therefore assigned both views the RNA cluster label of their shared source spot and, on the dissociated target, the CellTypist prediction of their shared nucleus (Methods, Section 4.1). Because source and target use different label vocabularies, these metrics are compared across models within each sample rather than between samples.

**Biological conservation.** Biological conservation was evaluated against these shared annotations. Leiden clustering was performed on a 15-nearest-neighbor graph, with the resolution optimized over 10 candidate values, and agreement with the biological annotation was quantified using normalized mutual information (NMI) and the adjusted Rand index (ARI). The label average silhouette width (ASW), denoted $\text{ASW}_{\text{label}}$, was computed from Euclidean distances and rescaled from $[-1, 1]$ to $[0, 1]$. Local preservation of biological labels was quantified using the cell-type local inverse Simpson's index (cLISI). It uses the same local inverse-Simpson calculation as iLISI, but with biological labels and reversed scaling, such that 1 indicates locally label-pure neighborhoods. cLISI complements the modality-mixing metrics by capturing preservation of biological structure. The reported biological-conservation score is the unweighted mean of the four enabled components,

$$B_{\text{bio}} = \frac{\text{NMI} + \text{ARI} + \text{ASW}_{\text{label}} + \text{cLISI}}{4}. \tag{S4}$$

**Uncertainty estimation.** Uncertainty was estimated by resampling, using two different procedures because the two families of metrics decompose differently. FOSCTTM is a mean of per-observation scores, so 95% percentile intervals were estimated by nonparametric bootstrap with replacement (1,000 replicates). The remaining scIB metrics depend on neighborhood graphs and clustering and were therefore recomputed on $R = 20$ independent subsamples containing $\phi = 0.8$ of the observations. For these metrics, uncertainty was estimated from the dispersion across subsamples and rescaled to the full sample size:

$$\mathrm{CI}_{95\%} = \hat{\theta} \pm 1.96 \sqrt{\frac{\phi}{1-\phi}} \ \mathrm{sd}\left(\hat{\theta}_1, \ldots, \hat{\theta}_R\right), \tag{S5}$$

where $\hat{\theta}_r$ denotes the metric recomputed on subsample $r$. The factor $\sqrt{\phi/(1-\phi)}$ is the delete-$d$ jackknife scaling for subsamples drawn without replacement (Shao and Wu 1989): because each subsample retains most observations, subsamples overlap substantially and their dispersion alone would understate the variability of the full-sample estimate. These intervals quantify within-sample uncertainty for the evaluated section and do not represent variation across independent tissue samples.

**In-sample and transfer evaluation.** We distinguish performance measured on the sample used for model fitting from performance after transfer to the dissociated target. Hatched bars in Fig. S1 indicate in-sample performance, where metrics are computed on the same sample used to train the model. Transfer performance is evaluated on the dissociated target, whose observations were not used to train the encoders. This distinction is especially important for $1 -$ FOSCTTM because models trained to align paired RNA and ATAC representations are directly optimized for the correspondence that FOSCTTM measures. High in-sample $1 -$ FOSCTTM therefore shows successful alignment on the training data, whereas target performance provides evidence of transfer.

### S2 Graph-context distillation recovers relative spatial positioning without coordinate inputs

The metrics of main-text Fig. 2e show that, through spatial-context distillation, the `SpatialJEPA` student preserves both molecular organization and neighborhood-level tissue structure without spatial information at inference. To specifically assess recovery of relative spatial position, we repeatedly split the spatial spots in half, using one half as queries and the other as references. For each query spot, we identified its nearest reference in representation space and measured the physical distance between their true tissue locations. To this end, we compared the `SpatialJEPA` teacher and student with a matched non-spatial ablation, MultiVI, and random assignment in a spatial position-recovery benchmark. All models were evaluated using the same position-recovery procedure over 40 random half-splits, with error expressed in spot widths (Fig. S2a; Section S9.10).

The graph-masked `SpatialJEPA` student retained substantial positional information despite operating without a spatial graph at inference, localizing 52% of query spots within two spot widths of their true positions, compared with 69% for the spatial teacher (Fig. S2b). By contrast, the matched non-spatial MultiGATE ablation and MultiVI each localized fewer than 1% of query spots within this tolerance (0.7% and 0.8%), placing both near the random spatial-pairing floor obtained from distances between independently and uniformly sampled pairs of source spots. As a simpler baseline, nearest-neighbor assignment using 30 principal components of the log-normalized expression

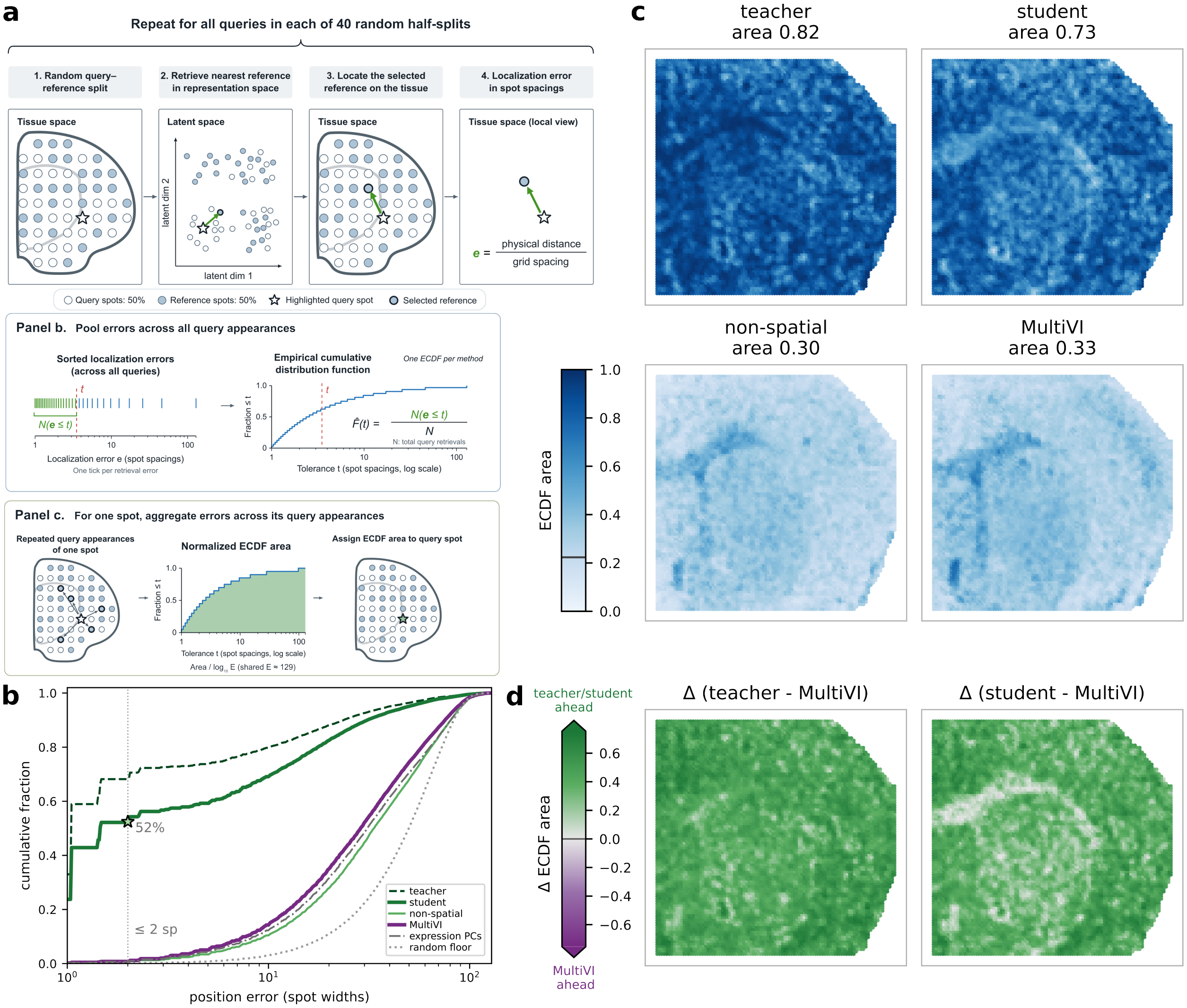

**Figure S2: Graph-context distillation transfers spatial position to an encoder without a spatial graph at inference.** **(a)** Schematic of the position-recovery evaluation. Across 40 random splits, half of the spatial spots serve as queries and the other half as references. For each query, the nearest reference is identified in representation space, and position error is measured as their physical distance in tissue space, normalized by the median nearest-neighbor spot spacing. Errors pooled across all query appearances yield the ECDFs in (b), whereas errors collected across repeated appearances of individual spots yield the per-spot ECDF areas in (c,d). **(b)** Position-error distributions, shown as empirical cumulative distribution functions (ECDFs), with query spots excluded from the reference pool during nearest-neighbor retrieval (all source spots were used to train the encoders). The methods compared are the joint RNA–ATAC latents of the `SpatialJEPA` teacher and student, the non-spatial MultiGATE ablation, and MultiVI, together with 30 expression principal components and a random spatial-pairing baseline. Position error corresponds to the true spatial distance, in spot widths, between each query spot and its nearest reference spot in representation space. The vertical reference line marks the 2-spot width position along the ECDF curves, and the green star marks the 52% of query spots that the student localizes within that tolerance. The x-axis uses a logarithmic scale. **(c)** Per-spot ECDF area, defined as the area under each spot's ECDF across repeated query appearances, bounded by 0 and 1 and shown on a shared scale. The horizontal line in the color bar marks the random-assignment baseline. **(d)** Per-spot difference in ECDF area between the `SpatialJEPA` teacher (left) or student (right) and MultiVI, on a shared color scale. Green indicates better `SpatialJEPA` localization, and purple indicates better MultiVI localization.

matrix localized only 0.7% of query spots within the same tolerance. The student and the non-spatial ablation are matched in architecture, training data, and inference-time information, differing only in whether the representation was distilled from a spatially aware teacher. Their marked difference in position recovery therefore isolates the contribution of spatial-context distillation, indicating that it transfers positional context into the student representation without requiring a spatial graph at inference.

To summarize the recovery of relative spatial position across the full error distribution, we collected, for each spot, the position errors from the repeated splits in which that spot served as a query. These errors were then used to construct a spot-specific ECDF. Each spot was summarized by the area under its ECDF on the logarithmic error axis (ECDF area), scaled from 0 to 1, with larger values indicating more precise localization (Section S9.10). Averaging these per-spot ECDF areas across the tissue gives 0.82 for the teacher and 0.73 for the student, substantially above all baselines and the random-assignment floor of 0.22 (Fig. S2c). Among the baselines, expression principal components (0.32) nearly match MultiVI (0.33), while the non-spatial MultiGATE ablation (0.30) remains close to chance. Thus, positional information is not readily recovered from molecular variation or cross-modal integration alone, whereas spatial-context distillation yields a substantial gain.

At a finer spatial scale, both the `SpatialJEPA` teacher and student retain their advantage over MultiVI in relative spatial recovery, with higher ECDF areas at 98% and 94% of spots, respectively (Fig. S2d). The modalities contribute unequally: evaluated on its RNA-side latent alone, the student reaches 0.57 against the teacher's 0.75, whereas on its ATAC-side latent it reaches 0.80 against 0.82. The student nevertheless outperforms MultiVI across every tissue domain in both modality-specific evaluations (Fig. S3a,b).

Taken together, these results show that spatial-context distillation transfers substantial positional information.

### S3 Position recovery on RNA-side and ATAC-side latents

Fig. S2 scores each model on its joint latent: for the `SpatialJEPA` teacher and student and the non-spatial ablation, the mean of the $\ell_2$-normalized RNA and ATAC embeddings of MultiGATE, and for MultiVI its joint latent. Figure S3 repeats the same estimator on each modality's latent separately and averages the per-spot ECDF areas within the clusters of that modality (`RNA_clusters` and `ATAC_clusters`, shown in Fig. S3c and main-text Fig. 2a). On the RNA-side latents (Fig. S3a), the student reaches an ECDF area of 0.57 against the teacher's 0.75 (30% versus 59% of queries within two spot widths), and leads MultiVI (0.33) in all eleven `RNA_clusters` (+0.06 to +0.35), with the smallest advantage in white matter. On the ATAC-side latents (Fig. S3b), the student comes close to the teacher (ECDF area 0.80 versus 0.82; 65% versus 70% of queries within two spot widths), whereas the non-spatial ablation and MultiVI remain at 0.33, and the student leads MultiVI in all fourteen `ATAC_clusters` (+0.26 to +0.55). The student–teacher gap in spatial-neighborhood

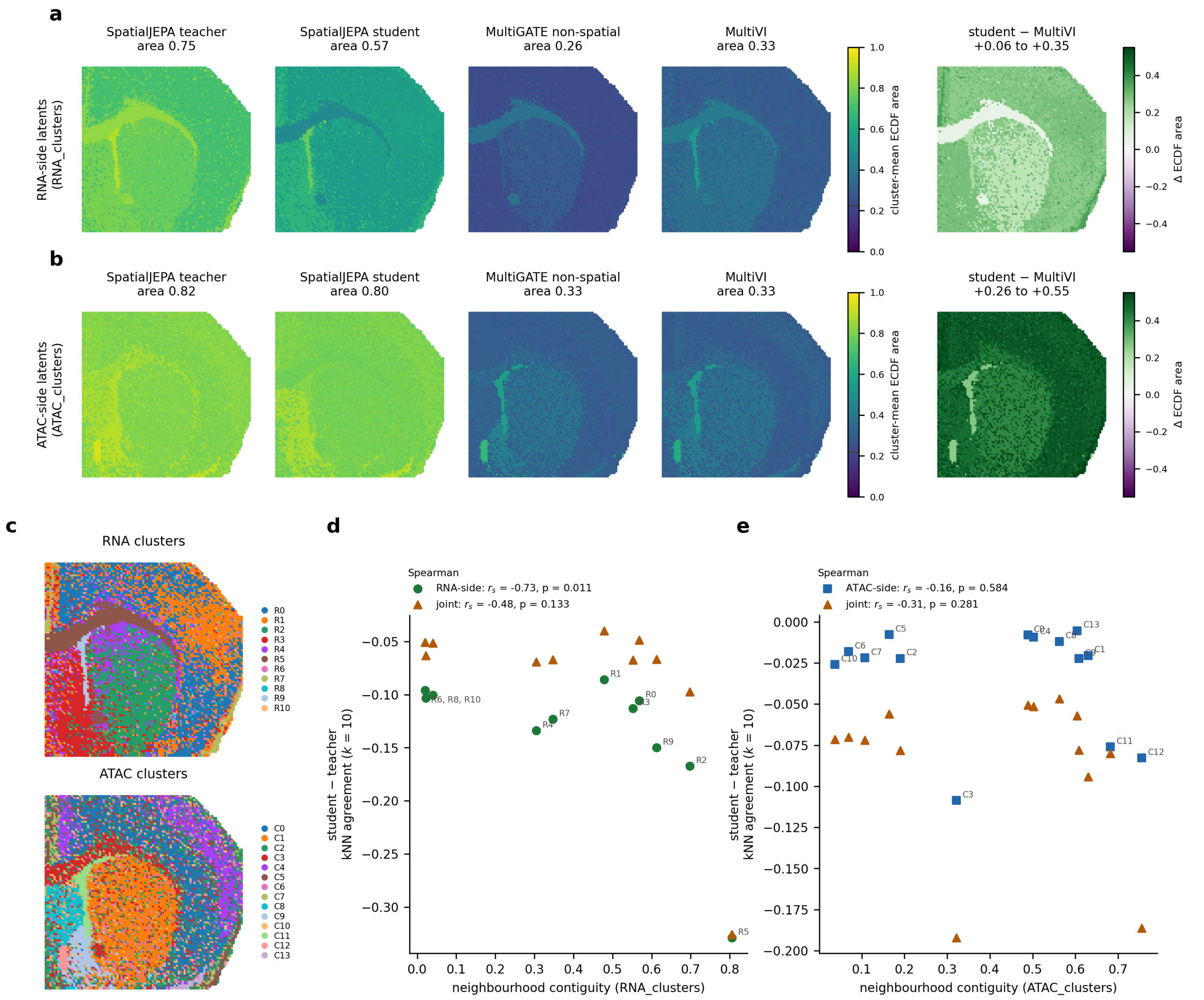

**Figure S3: Position recovery on the RNA-side and ATAC-side latents of the P22 spatial source.** **(a)** RNA-side latents: per-spot ECDF area averaged within each `RNA_clusters` domain for the `SpatialJEPA` teacher and student, the non-spatial MultiGATE ablation, and MultiVI (titles give the mean over spots), and the difference between student and MultiVI (right; green, student ahead). **(b)** The same for the ATAC-side latents, averaged within `ATAC_clusters`. All areas use the logarithmic axis of Fig. S2, from 1 to 129 spot widths; the line on each left color bar marks the random-assignment floor. **(c)** The `RNA_clusters` (top) and `ATAC_clusters` (bottom) annotations on the source section, as in main-text Fig. 2a. **(d)** Neighborhood contiguity of each `RNA_clusters` domain against the student-minus-teacher difference in spatial-neighborhood agreement ($k$-nearest-neighbor overlap, $k = 10$), for the RNA-side and joint latents. **(e)** The same for the `ATAC_clusters` domains, for the ATAC-side and joint latents. Values closer to zero indicate a smaller student–teacher gap; legends give Spearman correlations.

agreement widens with domain contiguity mainly on the RNA side: across `RNA_clusters` the correlation is $r_s = -0.73$ ($p = 0.011$) for the RNA-side latents and $-0.48$ ($p = 0.13$) for the joint latents, with the largest gap in white matter (Fig. S3d), and across `ATAC_clusters` it is $-0.16$ ($p = 0.58$) for the ATAC-side latents and $-0.31$ ($p = 0.28$) for the joint latents (Fig. S3e).

### S4 Held-out Multiome placement scored against the published annotations

Main-text Fig. 2f scores held-out Multiome clusters placed on the P22 spots against the spatial teacher's Leiden domains. To test whether the student's advantage depends on this reference, the same placed clusters were scored against the published P22 RNA and ATAC cluster annotations (Zhang et al. 2023) (Fig. S4). Placement used the pipeline of Fig. 2f, with each model's own Leiden clusters of its Multiome embedding at $K = 10$, the resolution closest to the eleven published RNA clusters (Section S9.9).

Against the published annotations, the models were close: under 15-nearest-neighbor coupling the student scored 0.31 (RNA) and 0.27 (ATAC), against 0.14–0.27 and 0.20–0.26 for the other models, and under optimal transport the student was marginally lowest (Fig. S4). The student's advantage in main-text Fig. 2f is therefore specific to the graph-informed teacher domains. The published RNA and ATAC annotations agree only modestly with each other (ARI 0.23), while the teacher's own P22 domains, without any placement, agree with them at ARI 0.29 (RNA) and 0.21 (ATAC), emphasizing that these references capture partly different partitions of the same tissue. Consistently, each model's own in-sample P22 clusters showed a similar level of agreement with the published annotations (ARI 0.21–0.29 for RNA and 0.21–0.26 for ATAC).

### S5 RNA and ATAC PLS components on the P22 source and the aligned target

Main-text Fig. 3 relates the first three RNA PLS components of the source and the aligned target through spatial maps and gene-set enrichment. Figure S5 shows the corresponding ATAC PLS components, fitted to accessibility and sign-aligned to the RNA components, on the color scale of the RNA maps in main-text Fig. 3c. Figure S6a shows the gene weights behind the RNA gene-set enrichment; of the genes in its six gene sets, 19 are in the 2,212-gene panel. The myelin genes *Mobp*, *Mal*, and *Mag* keep their positive weights on PLS_0 in the target, whereas the striatal genes *Pde10a*, *Rgs9*, and *Gng7*, which also weigh on source PLS_0, lose that weight in the target and carry PLS_1 in both datasets. On PLS_2, 18 of the 19 genes have the same sign in source and target. The marker genes keep their spatial localization after transfer: the myelin genes follow the white-matter tracts and the striatal genes the striatum, in the source and in the aligned target (Fig. S6b; *Mobp* is shown in main-text Fig. 3e).

### S6 Teacher early stopping and held-out evaluation of the mouse RNA–MSI model

The mouse RNA–MSI analyses of the main text are trained and evaluated on one section (V11L12-109_B1). To test the representation on tissue it was not trained on, the same SpatialMETA-backbone

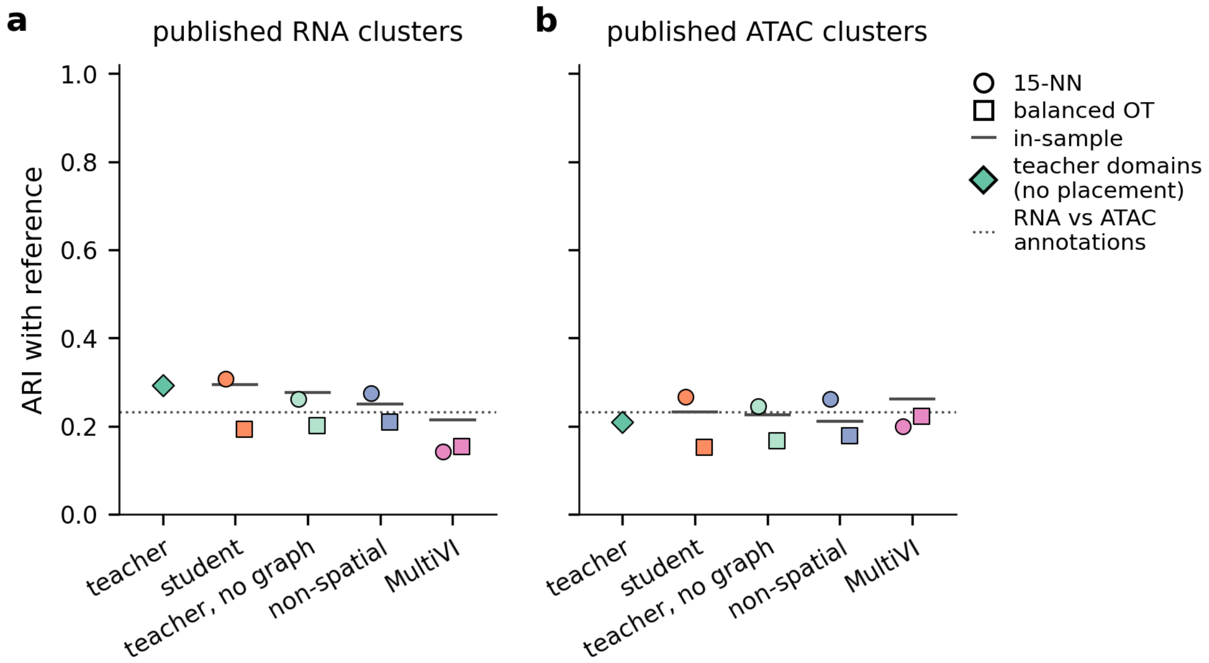

**Figure S4: Held-out Multiome placement scored against the published P22 annotations.** Each model's Leiden clusters of the held-out 10x Multiome nuclei ($K = 10$) were placed on the P22 spots by 15-nearest-neighbor (circles) or balanced optimal-transport (squares) coupling, as in main-text Fig. 2f, and scored by ARI against **(a)** the published RNA clusters and **(b)** the published ATAC clusters of Zhang et al. (2023). Scores against the spatial teacher's domains are in main-text Fig. 2f. Horizontal dashes, each model's own P22 embedding clustered in the same way ($K = 10$) and scored against the same reference (in-sample). Diamonds, the teacher's own P22 domains ($K = 10$) scored directly against the published annotations, without placement; dotted line, agreement between the published RNA and ATAC annotations. "Teacher, no graph": the teacher's encoder applied with an identity graph.

configuration was trained on one striatal section at a time from each of three animals of the 6-OHDA cohort (sections V11L12-109_A1, _B1, and _C1; three seeds each) and scored on the sections of the other two animals. In a second design, models were trained on one half of a section and scored on the other half and on the two other animals. The gene panel, loss weights, and schedule followed the main-text mouse model, and MSI peaks were matched across sections by $m/z$. Held-out sections were encoded from their measured RNA and MSI profiles; the teacher used the held-out section's own spatial graph. Graph-free controls were trained alongside: a student distilled from an identity-graph teacher, that identity-graph model itself, the stock non-spatial SpatialMETA, and ten principal components of the RNA and MSI input (input PCA).

#### S6.1 Teacher early stopping

On held-out spatial blocks of its training section, the teacher's training loss reaches a minimum within a few dozen epochs and then rises (Fig. S7a): for section V11L12-109_B1 the best epoch was 29 of 1,000, and across the 27 benchmark trainings it ranged from 22 to 35 epochs on whole sections and from 26 to 67 on half sections. A teacher trained for the full schedule therefore fits its

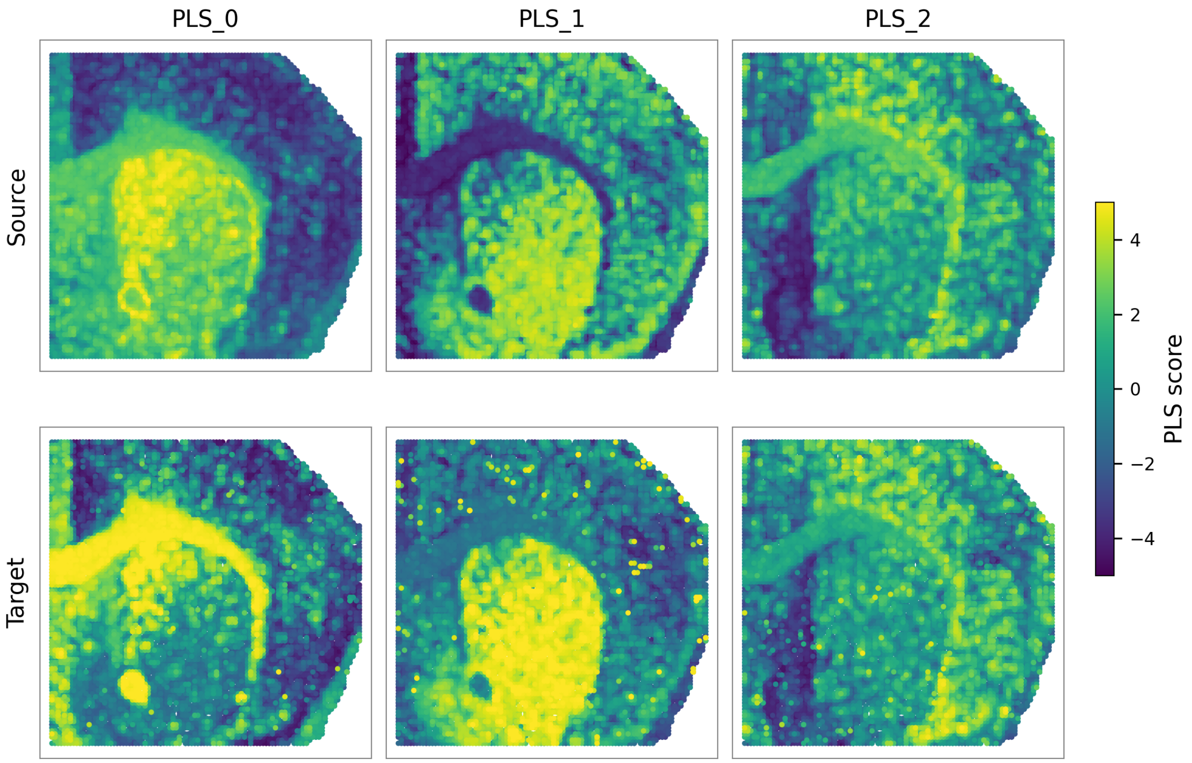

**Figure S5: ATAC PLS components on the P22 spatial source and the aligned target.** Spatial maps of ATAC PLS_0–2 scores in the source (top) and the aligned 10x Multiome target (bottom), from nine-component PLS models fitted from the student embedding to accessibility, independently in each dataset. Signs are aligned to the RNA components, and the color scale is shared with the RNA maps of main-text Fig. 3c.

training spots beyond what generalizes. The mouse model of the main text, and every mouse figure of this supplement, uses an early-stopped teacher (Section S9.6). A selection pass trains the teacher alone on 8 of 10 spatial blocks ($k$-means on spot coordinates), scores the other 2 after every epoch with the training objective at the final KL weight, and stops 100 epochs after the last improvement. The teacher is then trained on all spots for the selected number of epochs, and the student keeps distilling from this frozen teacher for the full 1,000 epochs.

Early stopping makes the teacher reproducible across seeds and training tissue (Fig. S7b). On the same spots, measured by symmetric out-of-fold linear $R^2$ under spatial-block cross-validation, two seeds trained on the same section agree at 0.89 (0.71 for full-schedule teachers); a teacher trained on the scored section and one trained on another animal agree at 0.81 (0.50); and two teachers trained on other animals agree at 0.82 (0.55). The student also follows its teacher more closely on unseen tissue (Fig. S7c), with linear $R^2$ of 0.53 on the held-out half and 0.67 on held-out animals, against 0.42 and 0.51 with a full-schedule teacher. This reaches the agreement of a ridge map fitted per spot on the held-out teacher itself (0.51 and 0.68), the ceiling for a predictor that sees each spot's own profile but not its neighbors.

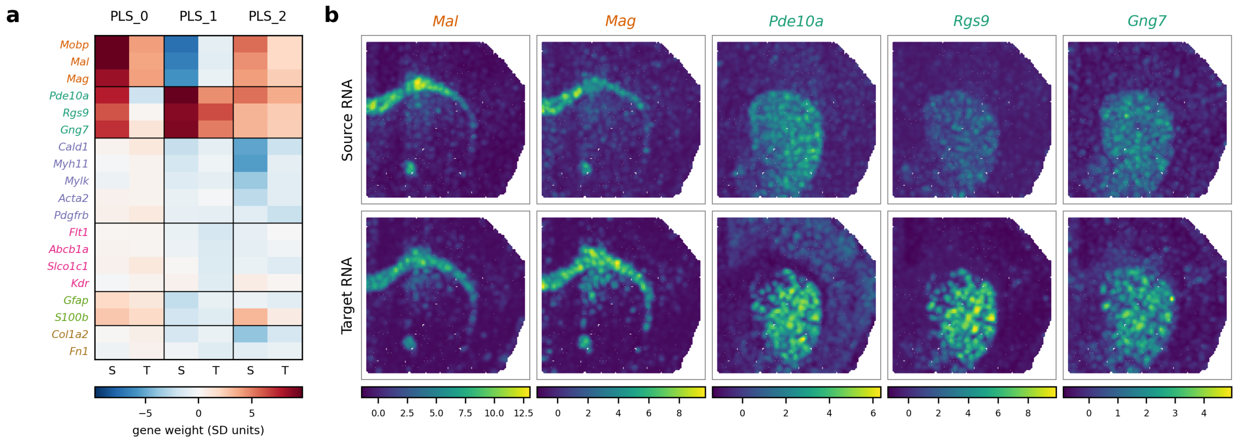

**Figure S6: Gene weights of the first three RNA PLS components and expression of their marker genes in the P22 spatial source and the aligned 10x Multiome target. (a)** Weights of the 19 genes of the six main-text gene sets (Fig. 3d) that are in the 2,212-gene panel, for source (S) and target (T) PLS_0–2. Weights are the PLS response weights used for the gene-set enrichment, scaled to unit standard deviation across genes within each model and component, with signs aligned across the source and target fits as in the main text. Gene names are colored by gene set. **(b)** Expression of the oligodendrocyte/myelin markers *Mal* and *Mag* and the striatal markers *Pde10a*, *Rgs9*, and *Gng7* in the source (top) and the aligned target (bottom), displayed as in main-text Fig. 3e: mean expression per spot, smoothed with a Gaussian kernel of one spot width, with one color scale per gene.

#### S6.2 Cluster agreement on held-out tissue

Representations of held-out tissue were scored by $k$-means clusters ($K = 5$, 10, and 20, averaged) against the early-stopped teacher's clusters of the same spots and against the annotated region $\times$ lesion compartments, and by the Moran's $I$ of the latent on the tissue (Fig. S7d–f). On held-out animals, the student's clusters agree with the teacher's more than those of any graph-free control: ARI 0.34, against 0.29 for the student of an identity-graph teacher, 0.27 for the identity-graph model, 0.26 for non-spatial SpatialMETA, and 0.24 for input PCA, with the student ahead of each control in all six train/test pairs. The same holds on held-out halves (0.27 against 0.20–0.21; six of six halves). The student's clusters also follow the region $\times$ lesion compartments (ARI 0.23, against 0.25 for the teacher and 0.13–0.18 for the controls), and its latent is smoother on the tissue (Moran's $I$ 0.64 against 0.44–0.46, in every held-out unit). The student of the identity-graph teacher differs from the student only in its teacher's graph, so these differences isolate the contribution of spatial distillation. A student of a full-schedule teacher reaches the same cluster agreement (ARI 0.34) with a less smooth latent (Moran's $I$ 0.54).

#### S6.3 Dopamine-lesion metrics on held-out tissue

The dopamine-embedding metrics of main-text Fig. 4d–f transfer less well (Fig. S8). In the training sections, teacher and student have similar structural gains (0.079 and 0.071, averaged over the three training animals). On held-out tissue, the teacher keeps a positive gain (0.049 on held-out halves and 0.055 on held-out animals), whereas the student's gain turns negative (−0.045 and −0.041), as

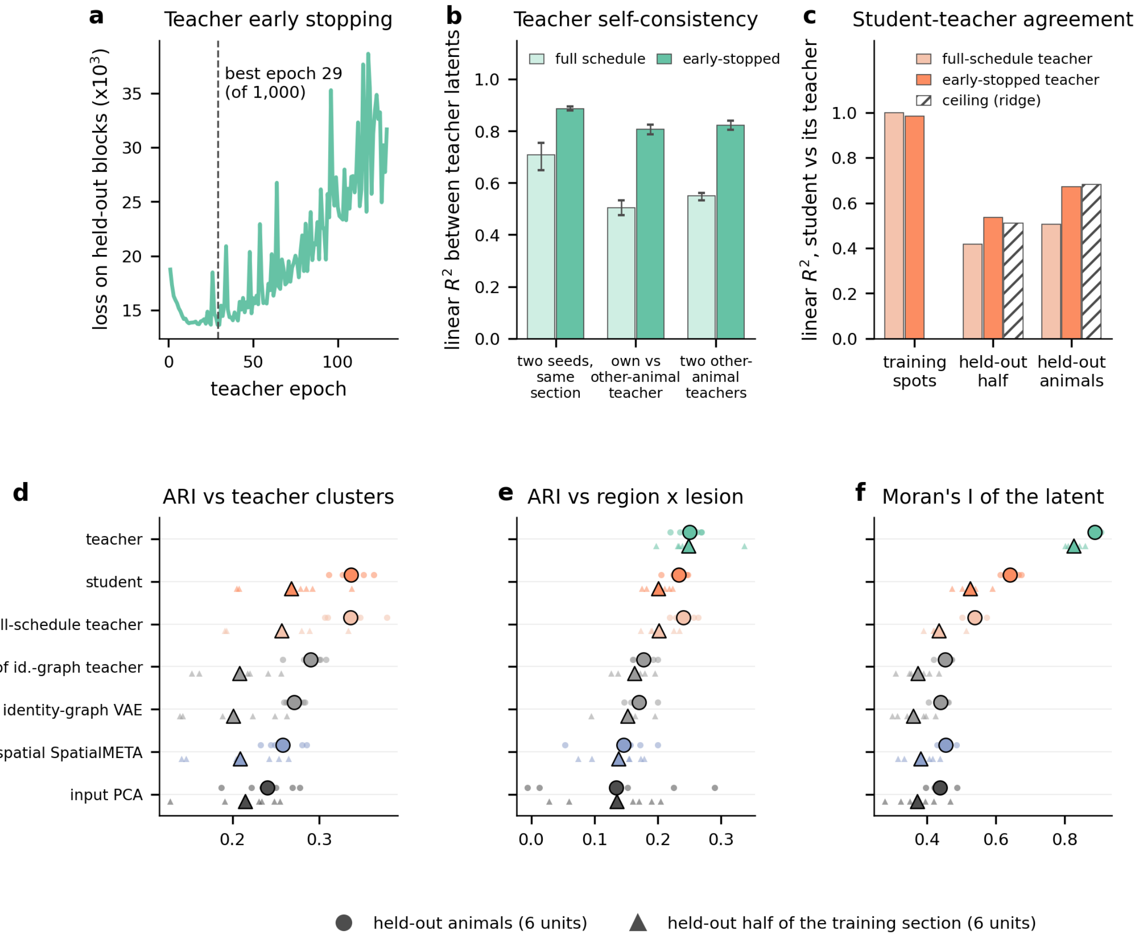

**Figure S7: Teacher early stopping and agreement on held-out mouse striatum.** **(a)** Loss on two held-out spatial blocks of section V11L12-109_B1 during the teacher's early-stopping pass (training objective at the final KL weight); the dashed line marks the selected epoch. **(b)** Agreement between teacher latents on the same spots (symmetric out-of-fold linear $R^2$ under spatial-block cross-validation; mean and range over the three animals), for full-schedule and early-stopped teachers: two seeds trained on the scored section, a teacher trained on the scored section versus one trained on another animal, and two teachers trained on other animals. **(c)** Linear $R^2$ between a student and its teacher on the training spots, on the held-out half of the training section, and on held-out animals, for students of full-schedule and early-stopped teachers; hatched, a ridge map fitted per spot on the held-out teacher (ceiling for a graph-free predictor). **(d–f)** Held-out tissue: ARI of $k$-means clusters ($K = 5$, 10, and 20, averaged) with the early-stopped teacher's clusters of the same spots (d) and with the region × lesion compartments (e), and Moran's $I$ of the latent on the tissue (f). Circles, held-out animals (six train/test pairs); triangles, the held-out half of the training section (six halves); small symbols, individual units (mean over three seeds); large symbols, their mean. The teacher uses the held-out section's spatial graph and is shown as a reference.

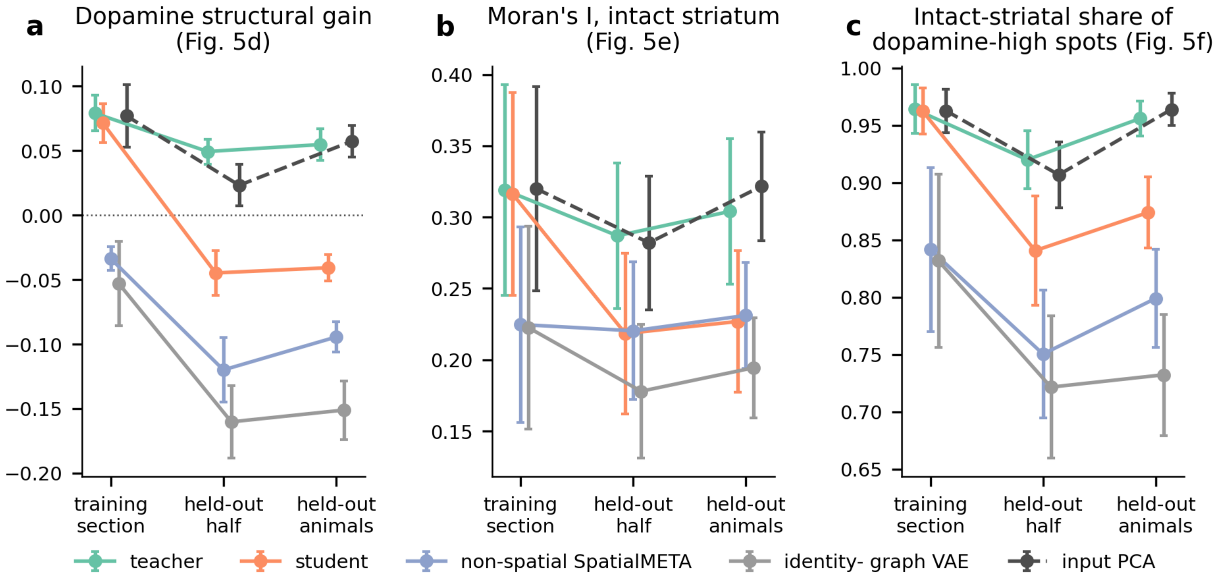

**Figure S8: The dopamine-embedding metrics of main-text Fig. 4d–f on the training section and on held-out tissue.** Structural gain (a), Moran's $I$ of dopamine on the latent graph in intact striatum (b), and the share of dopamine-high spots that lie in intact striatum (c; the log-odds of main-text Fig. 4f saturates on held-out sections), for the early-stopped teacher, its student, non-spatial SpatialMETA, the identity-graph model, and input PCA (dashed). Training section: in-sample, for each of the three training animals; held-out half: the other half of a half-trained section; held-out animals: the two other animals of each whole-section training. Mean and standard error over units (three, six, and six units), each the mean over three seeds.

does that of a student of a full-schedule teacher. The student still exceeds non-spatial SpatialMETA and the identity-graph model in structural gain and intact-striatal specificity in every held-out unit, but not in intact-striatum Moran's $I$, and input PCA matches or exceeds it on all three metrics. The lesion contrast of main-text Fig. 4d–f is therefore carried mainly by the spatial graph: on unseen tissue the graph-free student transfers tissue organization at the level of clusters and compartments (Fig. S7d–f), but not this finer contrast.

### S7 Supporting analyses for the mouse spatial RNA–MSI dopamine program

This section collects the supporting analyses behind the mouse dopamine results of the main text: a spatial sanity check on the teacher- and student-imputed metabolite channel, the dopamine dose-response across latent directions with an RNA-only control, a check of the metabolite-weight procedure over the same directions, the gene-selection step that defines the input to the TF-regulon over-representation test, the cis-regulatory analysis of the dopamine component, and the mapping of motif accessibility onto the source section.

#### S7.1 Teacher- and student-imputed dopamine on the P22 spatial section

The spatial RNA–MSI-trained SpatialMETA teacher carries a neural decoder from RNA to the MSI block, so it can be used to predict `msi:Dopamine` on a spatially resolved section for which no metabolite channel was measured. This teacher-side route consumes the section's spatial graph and is therefore available only for spatially resolved targets. Applying that decoder to the P22 spatial RNA–ATAC section places the imputed dopamine values over the striatum, corresponding to RNA cluster R2 and ATAC cluster C1 (Fig. S9a; clusters in Fig. S3c). The student route needs no spatial graph: a four-component PLS regression fitted on the RNA–MSI source, with the student embedding as predictors and the log-transformed, scaled measured MSI as responses, maps each spot's student embedding to metabolite values. It gives the same striatal pattern with more spot-to-spot variation (Fig. S9b); per spot, the two imputations correlate at $r = 0.56$. This check is qualitative, as dopamine was not measured on this section.

#### S7.2 Dopamine dose-response across latent directions and an RNA-only control

For 5,000 random directions in the 10-dimensional student latent, oriented so that measured source dopamine weighs positively, we computed each direction's dopamine weight in the source and the convergence of its target gene weights with the NMF dopamine program (AUROC, as in main-text Fig. 4j). Directions carrying more source dopamine converge more strongly with the NMF program in both targets (teacher on P22: $r = 0.75$; student on Multiome: $r = 0.76$), and the dopamine component C2 lies near the upper end of the random directions, above 90% and 95% of them, respectively (Fig. S10, top). The same analysis was repeated on an RNA-only latent, the first ten principal components of the 1,806 genes read by the student, with components defined from the same source metabolites (Fig. S10, bottom). It yields a dopamine component of similar weight (0.59, versus 0.63 for the student) and NMF convergence (AUROC 0.83 on P22 and 0.84 on Multiome). The dopamine axis is therefore anchored in striatal gene expression: measured MSI identifies it in the source and the student carries it into RNA-only targets, but its convergence with the NMF program does not by itself demonstrate information beyond RNA.

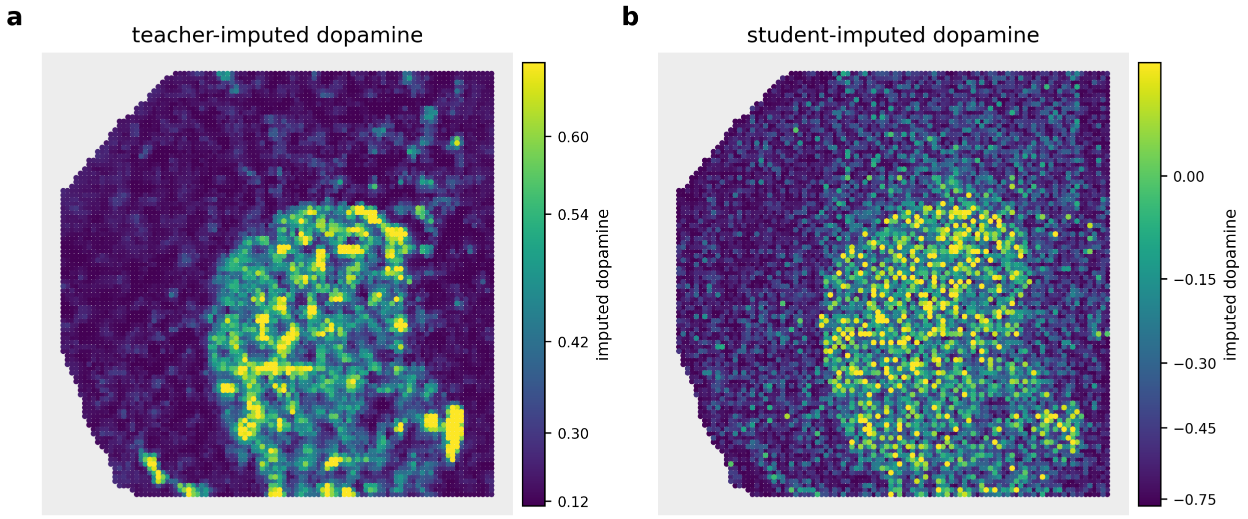

**Figure S9: Imputed dopamine values projected onto P22 spatial RNA–ATAC coordinates.** **(a)** The spatial RNA–MSI-trained SpatialMETA teacher's neural decoder applied to the teacher's RNA embedding of each spot. **(b)** A four-component PLS decoder, fitted on the spatial RNA–MSI source from the student embedding to the log-transformed, scaled measured MSI, applied to the student embedding of each spot. The two decoders output values on different scales (the teacher's own metabolite scale; scaled log-transformed intensities for the PLS decoder), so each panel has its own color bar, clipped at its 2nd and 98th percentiles with a square-law ($\gamma = 2$) color stretch.

#### S7.3 Metabolite weights over random latent directions

Main-text Fig. 4i reports metabolite weights in the Multiome nuclei, which carry no MSI: they are recovered by pairing the nuclei with source spots through optimal transport and fitting a PLS decoder on the target (main-text Methods; Fig. S19c). To check that this recipe returns the source's measured associations, it was applied to the 5,000 random latent directions of Fig. S10 and to the six components, and each direction's recovered dopamine weight was compared with its dopamine weight in the source (Fig. S11, top). Recovered and source dopamine weights agree across random directions ($r = 0.92$ for the teacher on P22 and 0.81 for the student on Multiome), and C2 recovers its source dopamine weight (0.76 and 0.71, against 0.63 in the source). Over all 779 metabolites, the recovered weights of a random direction match its source weights with a median $r$ of 0.95 (teacher, P22) and 0.89 (student, Multiome); for C2 the agreement is 0.97 and 0.94. As a control, the target latent was randomly rotated before pairing, which keeps its distribution but breaks its alignment with the source (Fig. S11, bottom). The agreement then disappears ($r = -0.20$ and $-0.12$ across directions; median all-metabolite $r = -0.29$ and $-0.32$), and the recovered dopamine weight of C2 falls to $-0.18$ and $-0.20$. The recovered weights therefore carry the source's measured metabolite associations onto the target cells through the student's alignment; they are not measurements of metabolites in the targets.

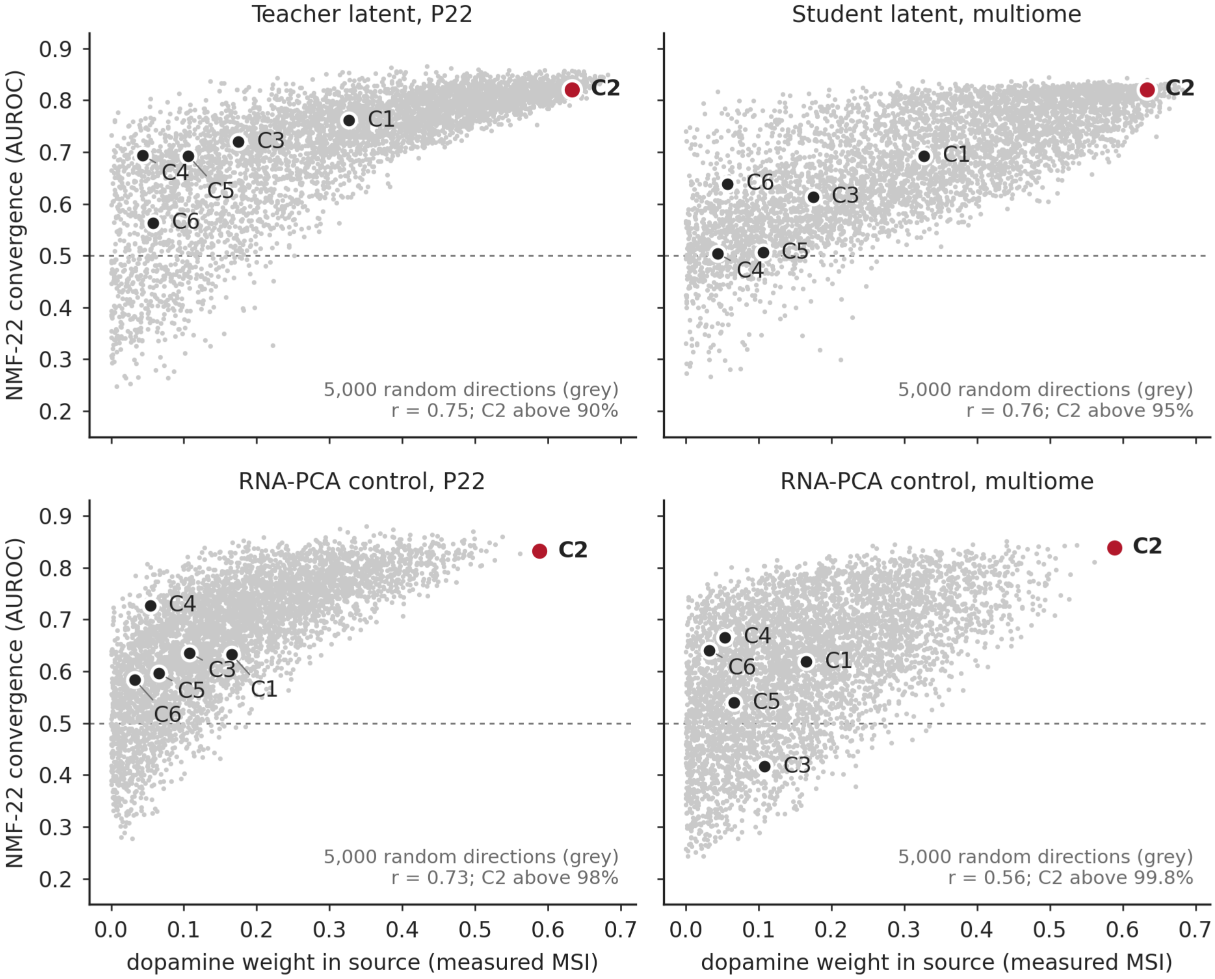

**Figure S10: Source dopamine weight versus NMF-program convergence over random latent directions.** Each gray point is one of 5,000 random latent directions; colored points are the six source-defined components, with the dopamine component C2 in red. The horizontal axis gives the direction's dopamine weight in the source (measured MSI); the vertical axis gives the AUROC of its target gene weights for the 50 most NMF-specific genes. The text in each panel reports the correlation over random directions and the share of random directions below C2. **Top**, `SpatialJEPA` teacher on P22 (left) and student on Multiome (right). **Bottom**, an RNA-only control latent (ten principal components of the student's input genes) analyzed in the same way.

#### S7.4 Spike-and-slab selection of dopamine-associated RNA loadings

The TF-regulon over-representation test of Section S7.5 needs a discrete gene set, whereas the dopamine component assigns a continuous weight to every gene. Rather than impose an arbitrary top-$k$ cutoff, we fitted a two-component Gaussian mixture to the absolute RNA weights of the Multiome student component C2, $|w|$, and treated the low- and high-magnitude components as a background "spike" and a selected "slab" (Fig. S12). The two-component model improves the Bayesian information criterion by $\Delta\text{BIC} = 19{,}472$ over a single-component model. The resulting posterior-probability threshold is $|w| = 0.047$, assigning $n = 3{,}848$ genes to the slab used for the

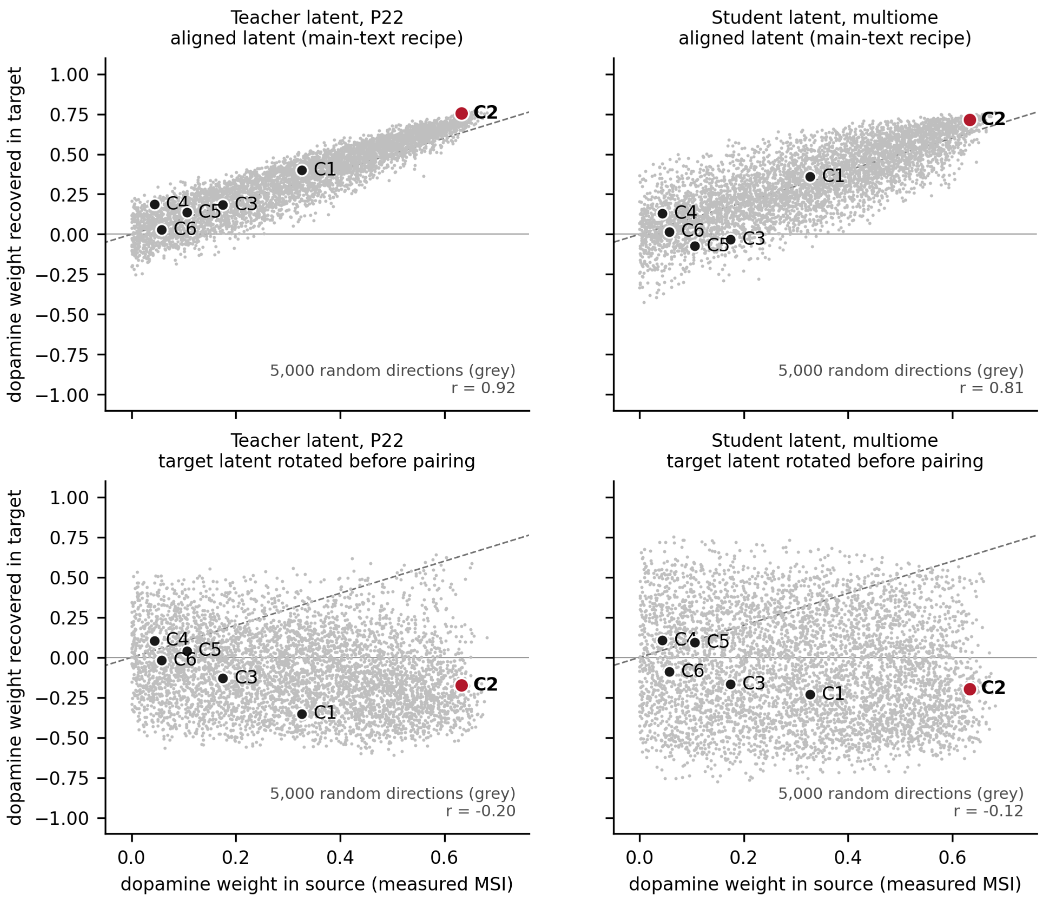

**Figure S11: The main-text metabolite-weight recipe over random latent directions, with a rotated-latent control.** Each gray point is one of the 5,000 random latent directions of Fig. S10; black points are the six source-defined components, with the dopamine component C2 in red. The horizontal axis gives the direction's dopamine weight in the source (measured MSI, intact hemisphere); the vertical axis gives its dopamine weight recovered in the target by optimal-transport pairing and a target-fitted PLS decoder (main-text Methods). The dashed line is the identity, and the text gives the correlation over random directions. **Top**, the recipe as used in the main text, for the `SpatialJEPA` teacher on P22 (left) and the student on Multiome nuclei (right). **Bottom**, the same with the target latent randomly rotated before pairing (20 rotations, each applied to 250 of the random directions; components show the median over the 20 rotations).

regulon test.

#### S7.5 The dopamine-associated component captures cell-type-specific cis-regulatory programs

After establishing biologically plausible and transferable gene signatures related to dopamine and associated metabolites, we focused on the C2 weights for chromatin accessibility. For this, we extracted candidate cell-type-specific cis-regulatory elements (cCREs) from Li et al. (2021) for striatal dopamine receptor type 2-expressing medium spiny neurons (D2-MSNs). We ranked all accessible ATAC peaks from the Multiome dataset by their signed C2 weights, computed after

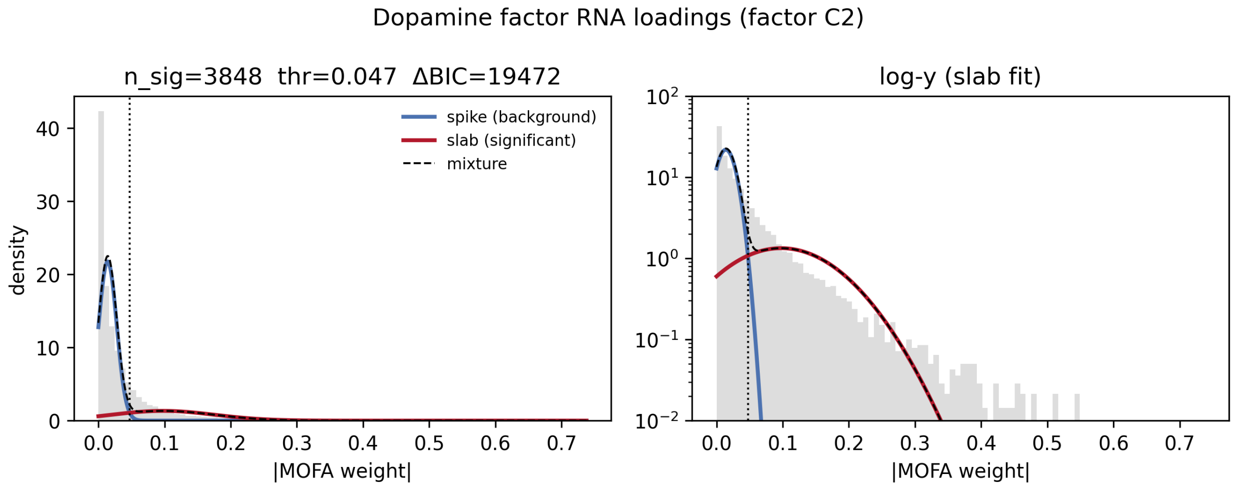

**Figure S12: Spike-and-slab separation of the Multiome student C2 RNA weights.** A two-component Gaussian mixture was fitted to the absolute RNA weights $|w|$ of the Multiome student component C2 to define the gene set used for the TF-regulon over-representation test of Section S7.5. Gray bars show the observed distribution of $|w|$; the blue curve is the low-magnitude "spike" component (background), the red curve the high-magnitude "slab" component (selected genes), and the dashed black curve their mixture. The dotted vertical line marks the resulting posterior-probability threshold at $|w| = 0.047$, above which genes are assigned to the slab. **Left**, linear density scale. **Right**, the same fit on a logarithmic $y$-axis, which resolves the slab across the sparse high-weight tail. The two-component model improved the Bayesian information criterion by $\Delta\text{BIC} = 19{,}472$ over a single component and assigned $n = 3{,}848$ genes to the slab.

adjusting the component score for sequencing depth and per-nucleus GC bias, and performed pre-ranked gene set enrichment analysis (GSEA), using the D2-MSN cCREs as reference (Section S9.16). In terms of normalized enrichment score (NES), D2-MSN cCRE-overlapping peaks were enriched toward the most positive weights (NES = 3.18, FDR $< 0.001$; Fig. S13a). Across the 66,520 ranked peaks, positive weights extended to rank 26,737, and the leading edge comprised 16,775 D2-MSN cCRE peaks.

To identify transcription factors (TFs) associated with these elements, we scanned the leading-edge (foreground) and remaining (background) D2-MSN cCRE sequences for known JASPAR 2024 vertebrate motifs, then tested each motif for enrichment in the foreground relative to the background while controlling for GC content (Section S9.16). In D2-MSN cCREs, 369 motifs were enriched robustly to GC content (FDR $q < 0.05$). Ordered by effect size, they were led by MEIS1 (GC-adjusted odds ratio 1.69), ZEB1, MEIS2, and MEIS3, followed by AP-1 motifs (FOSL1::JUNB, JUND) and PBX3 (Fig. S13b). The D2-MSN motif program is notable for its convergence on MEIS and PBX homeodomain factors: MEIS2 is a key regulator of the differentiation and subtype specification of MSNs derived from the lateral ganglionic eminence (Su et al. 2022). Together, these enrichments support an MSN-lineage regulatory program rather than generic cCRE motif density.

To assess the corresponding RNA-side program, we fitted a two-component Gaussian mixture to the absolute C2 RNA weights and retained genes assigned to the high-magnitude component (Fig. S12). TF-regulon over-representation analysis of this gene set recovered EGR1, SMAD4, and

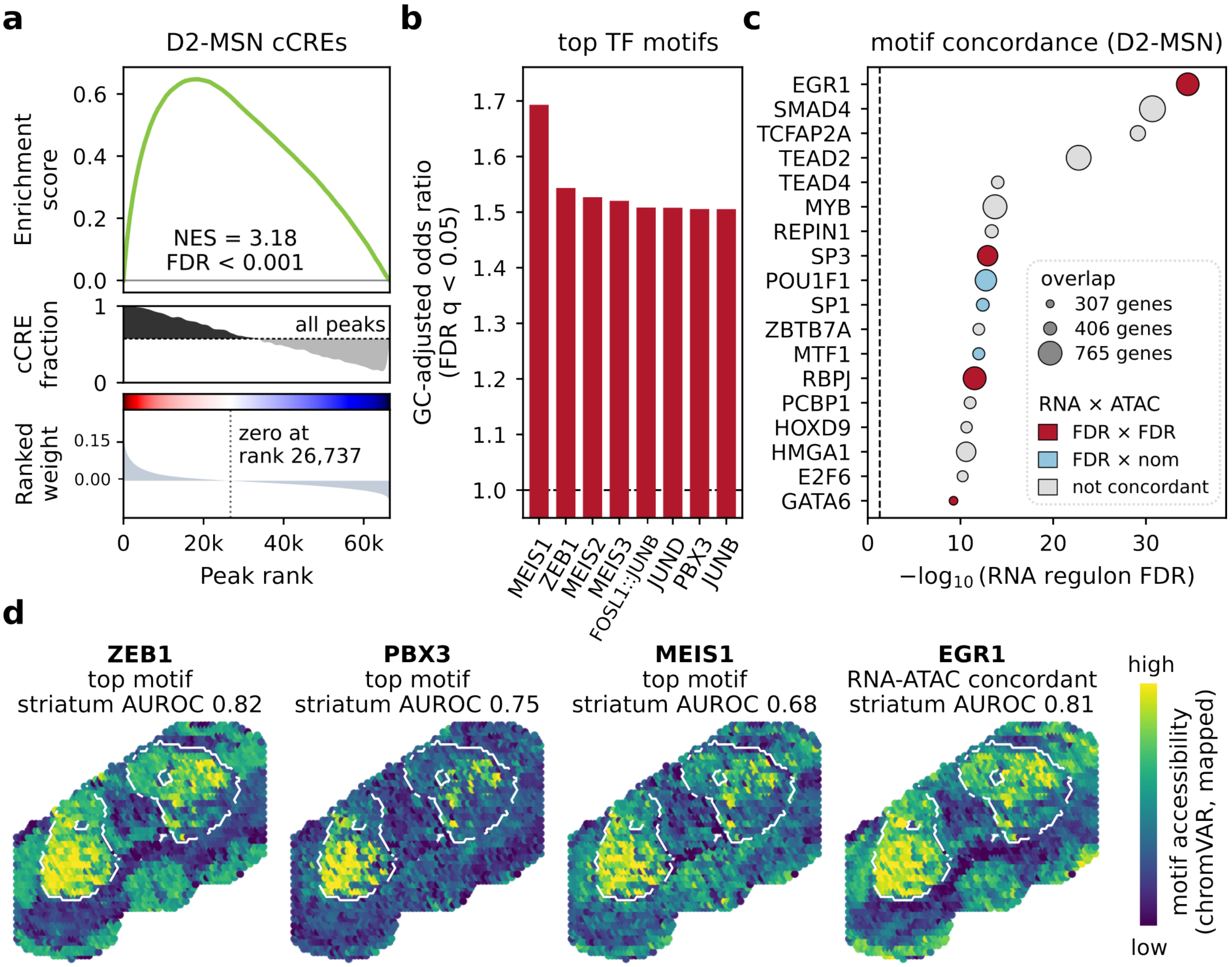

**Figure S13: Regulatory-chromatin analysis linking the dopamine-associated PLS component C2 to D2-MSN chromatin in the dissociated 10x Multiome target.** **(a)** Pre-ranked GSEA of all accessible peaks ranked by signed C2 peak weights, adjusted for sequencing depth and per-nucleus GC bias, against D2-MSN cCREs. **(b)** Top JASPAR 2024 TF motifs enriched in high-weight leading-edge D2-MSN cCREs relative to other D2-MSN cCREs; motifs robust to GC content at FDR $q < 0.05$, ordered by GC-adjusted odds ratio. **(c)** Concordance between RNA-regulon enrichment and independently tested ATAC-motif enrichment in D2-MSN cCREs; the horizontal axis shows RNA-regulon significance, dot size denotes overlapping-gene count, and color identifies the most stringent nominal/FDR combination reached. **(d)** Motif accessibility (chromVAR deviations) of the Multiome nuclei, placed on the spatial RNA–MSI source section by optimal-transport pairing of their RNA-derived student latents with the source spots, for three leading motifs of (b) and the RNA–ATAC-concordant EGR1 of (c). The striatum is outlined; AUROC, striatal versus other spots.

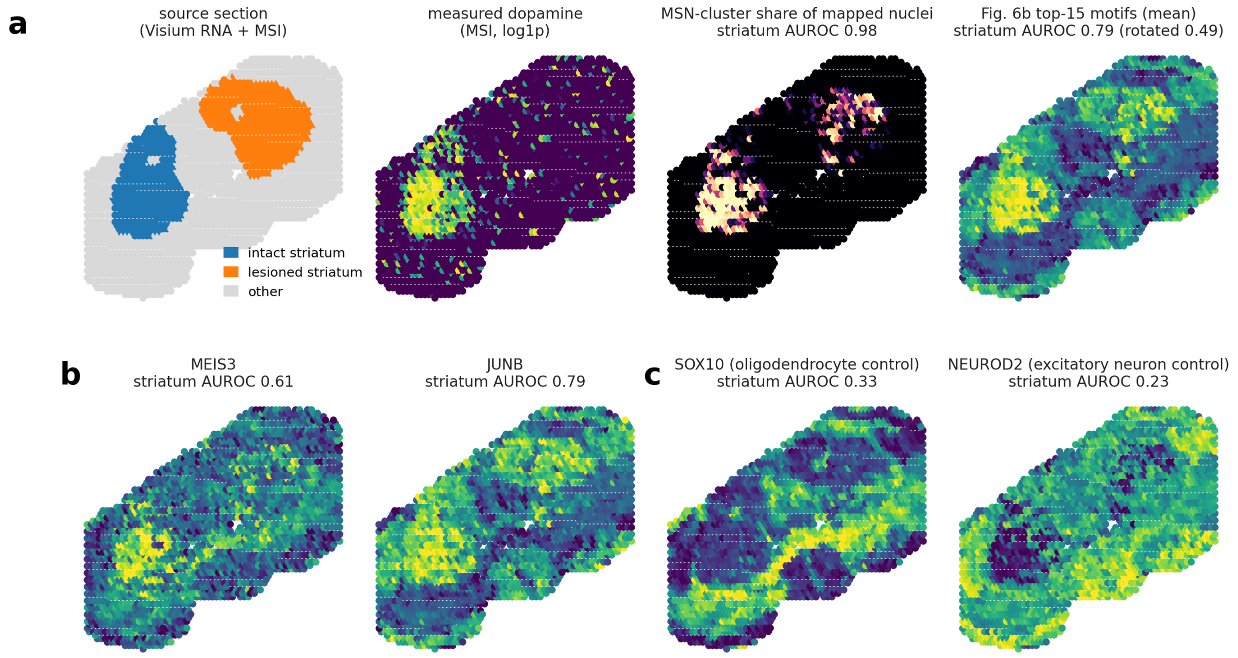

**Figure S14: Motif accessibility of Multiome nuclei mapped onto the spatial source section. (a)** Source section regions, measured dopamine, the share of each spot's paired nuclei from MSN clusters, and the mean accessibility of the 15 leading D2-MSN motifs of Fig. S13b, with the striatum AUROC and its mean over 20 random rotations of the nuclei latent. **(b)** Mapped accessibility of MEIS3 and JUNB, two leading motifs of Fig. S13b that Fig. S13d does not map, with their striatum AUROC. **(c)** Oligodendrocyte (SOX10) and excitatory-neuron (NEUROD2) control motifs.

TFAP2A (listed as TCFAP2A in the regulon library) most strongly, followed by TEAD2, TEAD4, MYB, REPIN1, SP3, POU1F1, and SP1 (all FDR $q < 0.05$). Comparing these RNA signals with the D2-MSN ATAC motifs identified EGR1, SP3, RBPJ, GATA6, and JUND as concordant at FDR level in both views (JUND not shown in Fig. S13c), and POU1F1, SP1, and MTF1 at FDR level on the RNA side with nominal ATAC support (Fig. S13c). The shared factor JUND is well supported: JUND shows robust, region-specific expression in mammalian brain and participates in activity-dependent neuronal transcription (Mellström et al. 1991; Okuno et al. 1997).

Because RNA and chromatin were measured in the same Multiome nuclei, the student representation also allows us to map their chromatin accessibility onto the spatial RNA–MSI source. We paired the nuclei with source spots through optimal transport and mapped per-nucleus motif accessibility onto the corresponding tissue locations (Section S9.16). Accessibility of ZEB1, PBX3, MEIS1, and the RNA–ATAC-concordant EGR1 concentrated in the striatum, particularly in the intact hemisphere (AUROC 0.68–0.82 for striatal versus other spots; Fig. S13d; Fig. S14). As a control, randomly rotating the target latent vectors before pairing reduced the striatal enrichment of the four leading motifs to near chance (mean AUROC 0.48–0.50 over 20 rotations), supporting the contribution of learned source–target alignment to their spatial localization. Together, these results demonstrate that the student representation can position chromatin-accessibility profiles from dissociated nuclei within an anatomically coherent spatial reference, extending the spatial interpretation of regulatory programs beyond datasets with directly measured spatial chromatin accessibility.

Together, the enrichment of MSN-associated cCREs and the concordant RNA–ATAC signals link the dopamine-associated C2 component to a striatal cis-regulatory program in the Multiome target.

##### S7.6 Motif accessibility mapped onto the source section

Fig. S13d places the chromatin of the Multiome nuclei on the spatial source section: the nuclei are paired with the source spots through optimal transport between their RNA-derived student latents, and each spot receives the mean motif accessibility of its paired nuclei. The pairing places nuclei from MSN clusters almost exclusively in the striatum of both hemispheres (AUROC 0.98 for striatal versus other spots; Fig. S14a), and the striatal motif maps follow from this placement. The mean map of the 15 leading GC-robust D2-MSN motifs, of which Fig. S13b shows the top eight, separates striatal from other spots with an AUROC of 0.79, against 0.49 on average when the nuclei latent is randomly rotated before pairing (Fig. S14a). The rotations vary widely, so the 15 motifs exceed the median of the 20 rotations but only four of them (ZEB1, PBX3, JUNB, and JDP2) exceed all 20; 14 exceed at least 17 of them. Two leading motifs not mapped in Fig. S13d, MEIS3 and JUNB, are also enriched in the striatum (AUROC 0.61 and 0.79; Fig. S14b), whereas the oligodendrocyte and excitatory-neuron motifs SOX10 and NEUROD2 are depleted from it (AUROC 0.33 and 0.23; Fig. S14c).

#### S8 Metabolite transfer to the human caudate head

In the caudate-head (CaH) nuclei of main-text Fig. 5, metabolite weights are not measured but recovered: the nuclei are paired with the spots of each source section by optimal transport (OT), and a decoder fitted on the paired MSI gives each metabolite's weight on a component (main-text Methods, Section 4.6). This section checks that recovery on source tissue held out from it, shows which metabolites the transfer assigns to each CaH subclass, and describes a second component, C2, whose metabolite weights are also recovered and which separates astrocytes and ependymal cells from the other subclasses.

##### S8.1 Recovery of C1 metabolite weights on held-out source sections

Each of the three source sections was held out in turn. The components were re-derived from the other two sections, and the held-out section's C1 metabolite weights were computed from its measured MSI and, separately, recovered by OT from the other two sections (Fig. S15a). The components re-derived without a section match those of the full data ($|\cos| \geq 0.98$ for C1). The recovered weights agree with the measured ones at $r = 0.76$, 0.90, and 0.69 across the 787 MSI features for the held-out sections A1, C1, and D1. Dopamine is recovered among the strongest weights where it is measured among them (rank 5 and 1 against measured ranks 4 and 2 for A1 and

D1); when section C1 is held out, dopamine has the strongest measured weight but ranks 17th after recovery, with a recovered weight of 0.68 against 0.79 measured.

Averaged by subclass, the metabolite profiles transferred to the nuclei follow C1 (Fig. S15b). Hybrid MSNs receive the highest dopamine, serotonin, 3-MT, and histidine, followed by D1 MSNs and the Sst Chodl and Vip interneurons, whereas oligodendrocytes receive the lowest of these and the highest values of the features at $m/z$ 570.23 and 572.24, which weigh negatively on C1. Taurine, with a C1 weight near zero, varies little across subclasses. These values are the measured metabolite profiles of the source spots each nucleus is paired with, not measurements in the nuclei: they indicate the tissue context that the transfer assigns to each subclass. Because the decoded metabolites and C1 are read out of the same student latent, metabolites with large C1 weights largely follow C1 across subclasses.

#### S8.2 A second component, C2, separates astrocytes and ependymal cells

The second component (C2) accounts for 11% of the source RNA–MSI correlation structure, against 76% for C1, and is uncorrelated with C1 across CaH nuclei ($r = -0.03$). Its negative pole concentrates ependymal cells and astrocytes (subclass means $-4.2$ and $-2.1$; astrocytes against all other nuclei, AUROC 0.94), with negative RNA weights for *BMPR1B*, *WDR49*, *AQP4*, *GFAP*, and *STON2*; CN ST18 GABAergic neurons, OPCs, and oligodendrocytes score among the highest (Fig. S16a–c).

C2's metabolite weights are recovered in the nuclei beyond chance: the OT-recovered weights agree with the source-measured weights at $r = 0.94$, against a maximum of 0.28 when the CaH latent is randomly rotated before pairing (ten rotations), and at 0.96 and 0.80 in the early-stopped models trained with two other seeds (rotation maxima 0.61 and 0.40). On held-out source sections, recovery reaches $r = 0.63$, 0.62, and 0.68 (Fig. S17a), lower than for C1. The leading individual metabolites are less stable than the component: norepinephrine has one of the largest recovered weights but ranks between 1st and 60th across held-out sections, and between 3rd and 676th in their measured MSI; across seeds, features at $m/z$ 573–574 recur among the strongest positive weights. We therefore read C2 as a recoverable metabolic axis rather than as evidence for specific metabolites. In the nuclei, the features with negative C2 weights ($m/z$ 552.25 and 555.19) are transferred at their highest levels to ependymal cells and astrocytes (Fig. S17b).

Projected onto the CaH Xenium sections with its RNA weights, as C1 in main-text Fig. 5, C2 again places astrocytes and ependymal cells lowest (subclass means correlate with those in the nuclei at Spearman $\rho = 0.64$). Its negative pole follows the local density of ependymal cells, which line the lateral ventricle (Fig. S16d–f). This is clearest on a section with an extended ventricle wall (H20.33.026.CX18.CaH.03.02; bivariate Moran's $I$ $-0.63$ between ependymal density and C2, $-0.27$ unsmoothed), and present on the main-text Fig. 5 section, which holds 46 ependymal cells ($-0.27$, unsmoothed $-0.11$). The same holds for C2 within astrocytes. Because the RNA-weight projection captures variation within astrocytes poorly, astrocyte C2 was carried to the Xenium astrocytes by a ridge map on the 464 panel genes fitted within the snRNA-seq astrocytes (agreement with C2 on

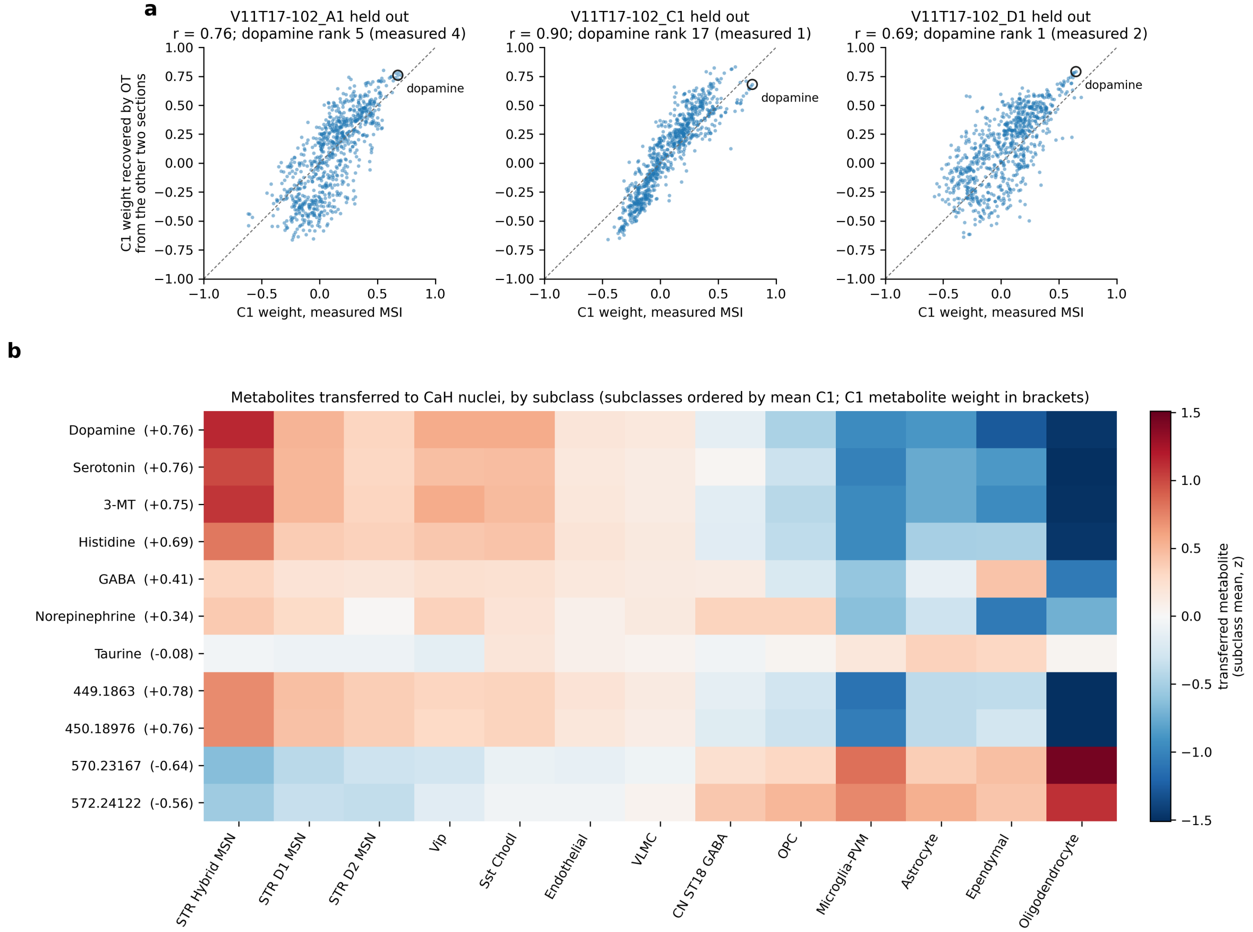

**Figure S15: Recovery of C1 metabolite weights on held-out source sections, and metabolites transferred to CaH nuclei.** **(a)** For each source section held out in turn (A1, C1, D1 of V11T17-102), the C1 weights of the 787 MSI features computed from the held-out section's measured MSI ($x$) and recovered by OT from the other two sections ($y$); components re-derived from the other two sections. Dashed, identity; circled, dopamine; ranks among the 787 features. **(b)** Metabolites transferred to CaH nuclei by OT pairing with the source spots, averaged by subclass (40,000 nuclei, the subsample used for the main-text metabolite weights; subclasses with at least 50 nuclei; $z$-scored per metabolite). Subclasses are ordered by mean C1; each metabolite's C1 weight is given in brackets. Unannotated features are labeled by $m/z$.

held-out donors: $r = 0.61$ per nucleus, 0.78 for donor means), adjusted for transcript counts within each section. Astrocyte C2 is lowest at the ventricle wall (Moran's $I$ with ependymal density $-0.57$ on both sections; unsmoothed $-0.40$ and $-0.23$). Within astrocytes, C2 is largely distinct from C1 in the nuclei ($r = 0.15$), but the two carried scores correlate more on Xenium ($r = 0.47$), as both draw on boundary-associated genes such as *CD44* and *SLC14A1*.

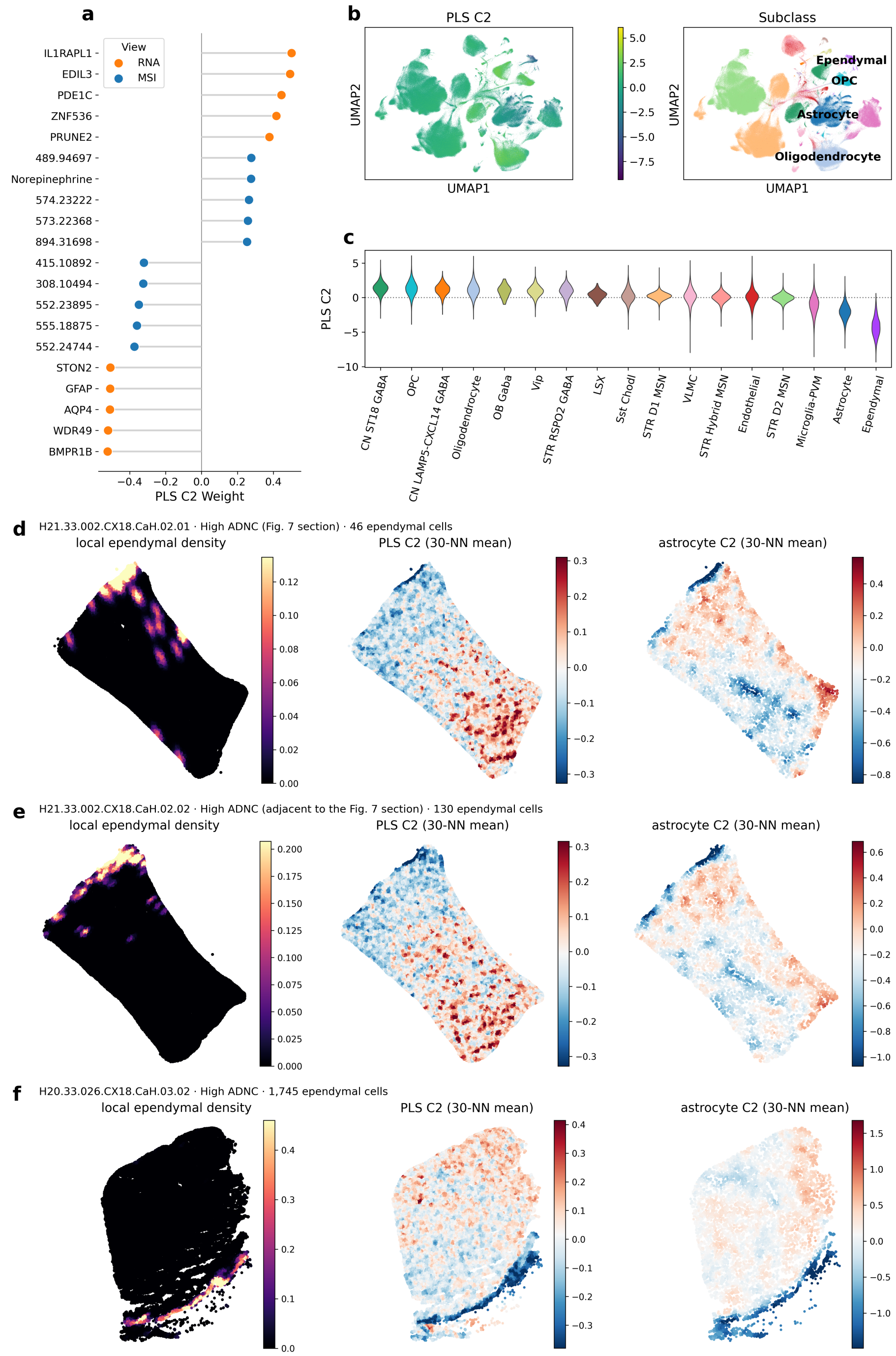

**Figure S16: The second shared component, C2, in CaH nuclei and Xenium sections.** **(a)** Strongest positive and negative C2 weights across RNA and metabolites, with metabolite weights recovered in the nuclei by OT pairing with the source spots. **(b)** RNA UMAPs colored by C2 score (left) and subclass (right); labels mark the subclasses at C2's poles. **(c)** C2 scores by subclass, ordered by descending mean. **(d–f)** Three CaH Xenium sections: the main-text Fig. 5 section (d), the adjacent section of the same tissue block (e), and a section with an extended ventricle wall (f). Left, local ependymal-cell density (kernel density estimate as for oligodendrocytes in main-text Fig. 5); middle, C2 projected onto all cells with its RNA weights, averaged over 30 nearest neighbors; right, C2 within astrocytes, carried from the snRNA-seq astrocytes by a ridge map on the panel genes, adjusted for transcript counts, and averaged over the 30 nearest astrocytes. Color scales are per section.

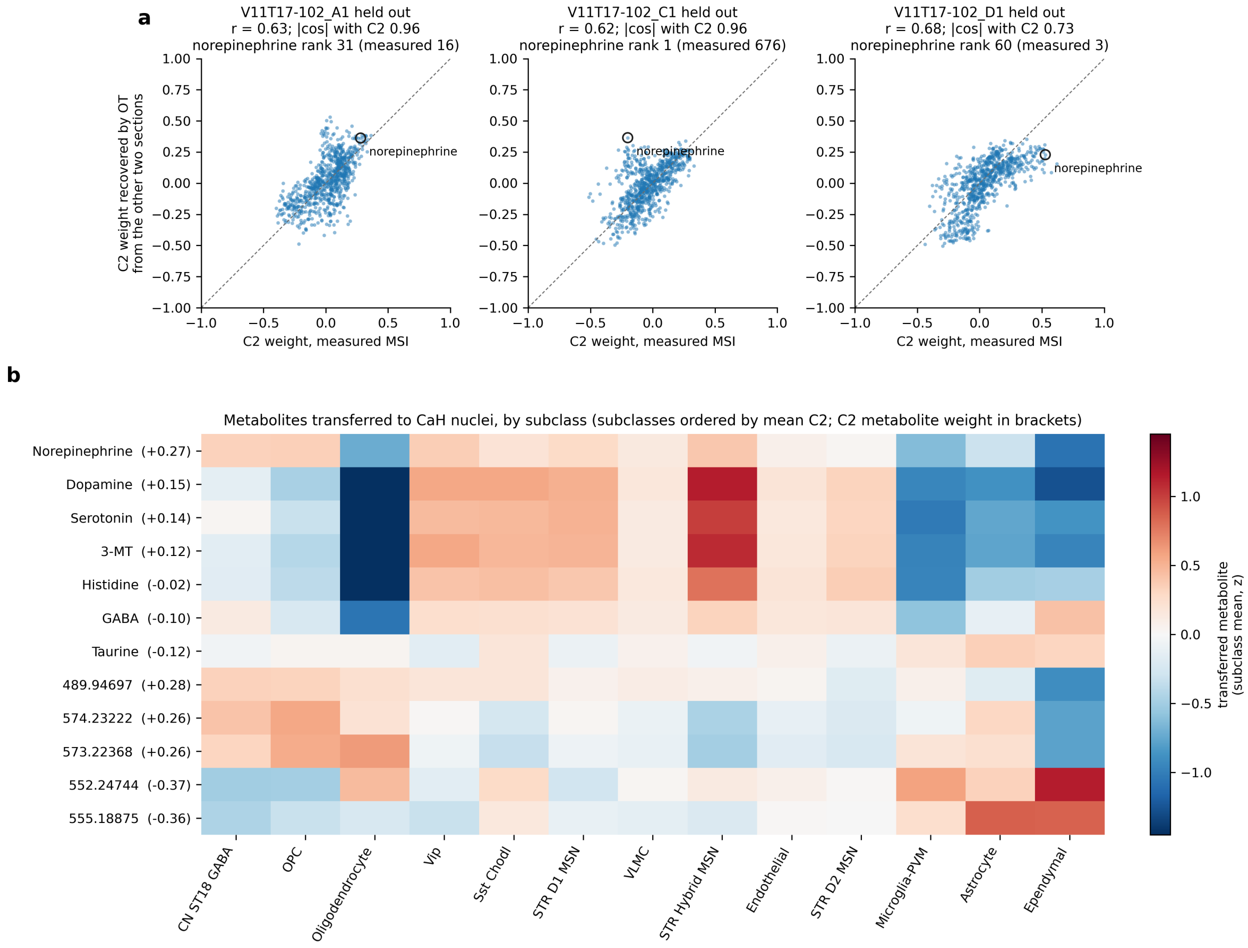

**Figure S17: Recovery of C2 metabolite weights on held-out source sections, and metabolites transferred to CaH nuclei. (a)** As Fig. S15a for C2. The component re-derived without each section is matched to the full-data C2 by $|\cos|$ (given above each panel) and its sign aligned; circled, norepinephrine. **(b)** As Fig. S15b, with the leading C2 features and subclasses ordered by mean C2; each metabolite's C2 weight is given in brackets.

### S9 Supplemental Methods

This section gives the full model objectives and derivations, the complete methods for analyses summarized in the main-text Methods, and the methods for analyses reported only in the Supplemental Material.

#### S9.1 Reprocessing and co-registration of spatial RNA–MSI sections

Mouse and human sections were reprocessed and spatially aligned following the original authors' workflow (Vicari et al. 2024). Visium libraries were processed with Space Ranger to generate gene-expression matrices registered to the paired H&E image, while MALDI-MSI intensities were represented as per-pixel feature matrices. For each section, the MSI pixel grid was manually co-registered to the H&E/Visium coordinate frame using the authors' alignment parameters and `ManualAlignImages` from STUtility (Bergenstråhle et al. 2020), allowing rotation, translation, anisotropic scaling and, where necessary, axis mirroring. MSI pixels were then matched to their nearest Visium spot by $k$-nearest-neighbor search and retained when the alignment distance was below 35 units, yielding section-level objects with spatially matched transcriptomic and metabolomic measurements. FMP-10-derived neurotransmitter features were annotated using the authors' MS/MS-validated target panel.

All sections analyzed here were derivatized with the FMP-10 matrix, which includes an MS/MS-validated neurotransmitter panel containing dopamine and other annotated neurotransmitters, alongside additional unlabeled metabolite features. Section V11T17-102_B1 was excluded because of a large signal near the center of the section that was considered artifactual.

#### S9.2 Dataset annotation and preprocessing details

**CellTypist annotation of the dissociated target** Predictions were refined by CellTypist's majority-voting procedure using a Leiden over-clustering of the target RNA neighbor graph at resolution 0.5, and the resulting per-nucleus label was copied to the paired ATAC view.

**Gene–peak distance prior** Candidate edges were assigned the MultiGATE distance-decay prior (Miao et al. 2025),

$$w(d) = \left(\frac{d + 1000}{1000}\right)^{-0.75}, \tag{S6}$$

where $d$ is the promoter distance in base pairs. This weighting assigns a value of 1 at the promoter, 0.21 at 10 kb, and 0.03 at the 150 kb boundary. The resulting gene–peak graph defines a fixed set of candidate regulatory interactions and associated prior weights; neither the candidate edges nor these distance-based weights are learned.

**RNA–MSI preprocessing** Heat-shock, mitochondrial, ribosomal, and DNAJ-family genes were removed, and RNA profiles were processed with `SCTransform` where available. MSI intensities were

**Algorithm S1** `SpatialJEPA` graph-context distillation

**Require:** Molecular profiles $\mathbf{x}$, spatial graph $G_s$, identity graph $G_{\text{id}}$
**Require:** Teacher parameters $\theta_t$, student parameters $\theta_s$
1: **for** each training epoch **do**
2:     **for** each batch $\mathcal{B}$ **do**
3:         $\min_{\theta_t} \mathcal{L}_{\text{backbone}}(\theta_t; G_s)$ ▷ update teacher
4:         $\mathbf{z}_t \leftarrow f_t(\mathbf{x}_{\mathcal{B}}, G_s)$ ▷ freeze teacher targets
5:         $\min_{\theta_s} \mathcal{L}_{\text{distill}}(\theta_s; G_{\text{id}}, \text{sg}(\mathbf{z}_t))$ ▷ update student; main-text Eq. 2
6:     **end for**
7: **end for**

library-size normalized, and FMP-10 neurotransmitter features were annotated using the authors' MS/MS-validated target panel; other $m/z$ features remained unlabeled. For the mouse model, the candidate pool comprised the top 2,000 highly variable genes from the P22 spatial RNA–ATAC dataset, ranked by normalized dispersion. For the human model, it comprised the top 2,000 genes from a ranking prioritizing the 442 protein-coding genes measured by CaH Xenium, followed by CaH snRNA-seq genes ranked by variance. After feature selection, RNA was represented using `SCTransform`-corrected counts where available, while MSI features were re-normalized, $\log(1+x)$-transformed, and scaled before model training. Spots with no RNA counts across the retained genes were excluded.

##### S9.3 Alternating teacher–student training

Training alternates between teacher and student updates. At each iteration, the teacher is first optimized under the backbone-specific objective using the spatial graph; its updated latent representations are then held fixed as targets for the student update under the identity graph (Algorithm S1). Unlike the exponential moving-average teacher used in many JEPA implementations (Assran et al. 2023), the `SpatialJEPA` teacher is optimized directly under its spatial multiomic objective and frozen only during the corresponding student update.

##### S9.4 MultiGATE backbone objective

The MultiGATE teacher retains the two-part objective of the original model (Miao et al. 2025), comprising reconstruction of the two modalities and a symmetric CLIP loss:

$$\mathcal{L}_{\text{backbone}}^{\text{MultiGATE}} = \mathcal{L}_{\text{recon}} + \mathcal{L}_{\text{CLIP}}, \tag{S7}$$

where

$$\mathcal{L}_{\text{recon}} = \left\|\mathbf{X}_1 - \hat{\mathbf{X}}_1\right\|_F + \left\|\mathbf{X}_2 - \hat{\mathbf{X}}_2\right\|_F, \tag{S8}$$

and

$$\begin{aligned}\mathcal{L}_{\mathrm{CLIP}} = & -\frac{1}{2N}\sum_{i=1}^{N}\log\frac{\exp[\mathrm{sim}(\mathbf{f}_i,\mathbf{g}_i)\exp(\mathrm{temp})]}{\sum_{j=1}^{N}\exp[\mathrm{sim}(\mathbf{f}_i,\mathbf{g}_j)\exp(\mathrm{temp})]} \\ & -\frac{1}{2N}\sum_{i=1}^{N}\log\frac{\exp[\mathrm{sim}(\mathbf{g}_i,\mathbf{f}_i)\exp(\mathrm{temp})]}{\sum_{j=1}^{N}\exp[\mathrm{sim}(\mathbf{g}_i,\mathbf{f}_j)\exp(\mathrm{temp})]}. \end{aligned} \tag{S9}$$

The reconstruction loss in Eq. S8 is distinct from the mean-squared-error distillation loss in main-text Eq. 2, which instead compares the teacher and student latent representations. Here, $\mathbf{X}_1$ and $\mathbf{X}_2$ denote the RNA and ATAC input matrices and $\hat{\mathbf{X}}_1$ and $\hat{\mathbf{X}}_2$ their reconstructed counterparts. $\mathbf{f}_i$ and $\mathbf{g}_i$ are the projected, $\ell_2$-normalized RNA and ATAC embeddings of spot $i$, $\mathrm{sim}(\cdot,\cdot)$ denotes cosine similarity, and temp is the learned log-temperature parameter initialized to 1. Matched RNA and ATAC profiles from the same spot define the positive pairs. The published MultiGATE reconstruction loss is written as a sum of per-spot Euclidean norms; our PyTorch implementation computes the corresponding reconstruction error as one Frobenius norm over the full RNA matrix and one over the full ATAC matrix. No additional likelihood or cross-modal imputation term is included in the MultiGATE backbone objective.

We implemented MultiGATE in PyTorch while retaining its spatial and cross-modal modeling objectives. Both encoders use hidden dimensions $512 \to 30$ per modality with non-linear activations and were trained full-batch for 3,000 epochs with Adam (learning rate $10^{-4}$, weight decay $10^{-4}$, gradient-norm clipping at 5).

##### S9.5 SpatialMETA backbone objective and the ST→SM extension

For spatial transcriptome–metabolome data, the `SpatialJEPA` backbone is SpatialMETA (Tian et al. 2025), a conditional variational autoencoder for cross-modal and cross-sample integration of spatial transcriptomics (ST) and spatial metabolomics (SM). For each spot, ST and SM encoders (each a single fully connected layer with a 128-dimensional output in the original architecture) map the transcriptomic and metabolomic feature vectors to modality-specific hidden representations. These are concatenated and mapped to a 10-dimensional joint variational posterior with mean $\boldsymbol{\mu}_{\mathrm{joint}}$, from which the joint latent representation $\mathbf{z}_{\mathrm{joint}}$ is sampled, whereas a shared linear map applied to each encoder representation gives deterministic 10-dimensional ST- and SM-specific latent means, $\boldsymbol{\mu}_{\mathrm{st}}$ and $\boldsymbol{\mu}_{\mathrm{sm}}$. Two-layer modality-specific decoders, conditioned on a one-hot section indicator for multi-section training, reconstruct transcriptomic counts under a zero-inflated negative-binomial likelihood and normalized, $\log(1+x)$-transformed, and scaled MSI intensities under a Gaussian likelihood, either from $\mathbf{z}_{\mathrm{joint}}$ or from the corresponding modality-specific latent mean. The original objective combines these four weighted reconstruction terms with a Kullback–Leibler (KL) divergence of the joint posterior from a standard normal prior, weighted by $\beta$, and, for horizontal integration across tissue sections, a multiscale-kernel maximum mean discrepancy (MMD) penalty that aligns the joint latent distributions of different sections.

For each spatial spot $i$, $\mathbf{X}_i^{\mathrm{st}}$ and $\mathbf{X}_i^{\mathrm{sm}}$ denote its transcriptomic and metabolomic feature vectors, respectively. The ST and SM encoders first map the two input modalities to modality-specific hidden representations,

$$\mathbf{q}_{\text{st},i} = \mathcal{F}^{\text{st}}_{\text{encoder}}\left(\mathbf{X}^{\text{st}}_i\right), \qquad \mathbf{q}_{\text{sm},i} = \mathcal{F}^{\text{sm}}_{\text{encoder}}\left(\mathbf{X}^{\text{sm}}_i\right). \tag{S10}$$

In the original architecture, each encoder comprises a single fully-connected layer with a 128-dimensional hidden representation. The two representations are concatenated,

$$\mathbf{q}_{\text{joint},i} = \left[\mathbf{q}_{\text{sm},i}; \mathbf{q}_{\text{st},i}\right], \tag{S11}$$

and mapped to the parameters of the joint variational posterior,

$$\boldsymbol{\mu}_{\text{joint},i} = \mathcal{F}^{\text{joint}}_{\text{mean}}\left(\mathbf{q}_{\text{joint},i}\right), \qquad \boldsymbol{\sigma}^2_{\text{joint},i} = \mathcal{F}^{\text{joint}}_{\text{var}}\left(\mathbf{q}_{\text{joint},i}\right), \tag{S12}$$

from which the joint latent representation is sampled by reparameterization,

$$\mathbf{z}_{\text{joint},i} \sim \mathcal{N}\left(\boldsymbol{\mu}_{\text{joint},i}, \operatorname{diag}\boldsymbol{\sigma}^2_{\text{joint},i}\right). \tag{S13}$$

The deterministic modality-specific latent means are obtained by applying a shared linear map to their respective encoder representations,

$$\boldsymbol{\mu}_{\text{st},i} = \mathcal{F}^{\text{single}}_{\text{mean}}\left(\mathbf{q}_{\text{st},i}\right), \qquad \boldsymbol{\mu}_{\text{sm},i} = \mathcal{F}^{\text{single}}_{\text{mean}}\left(\mathbf{q}_{\text{sm},i}\right). \tag{S14}$$

All three latent representations are 10-dimensional; only the joint representation is stochastic.

Each modality is decoded through a two-layer neural network. For the ST decoder, $\mathbf{r}^{\text{st}}_{\text{mean},i}$ denotes the library-size-scaled mean expression, $\mathbf{r}^{\text{st}}_{\text{var},i}$ parameterizes the negative-binomial dispersion through $\exp\left(\mathbf{r}^{\text{st}}_{\text{var},i}\right)$, and $\mathbf{r}^{\text{st}}_{\text{gate},i}$ contains the logits of the zero-inflation probabilities. The decoder maps a latent representation and, for multi-section training, its tissue-section indicator $\mathbf{B}_i$ to these quantities,

$$\mathcal{F}^{\text{st}}_{\text{decoder}}\left(\mathbf{u}_i, \mathbf{B}_i\right) \to \left(\mathbf{r}^{\text{st}}_{\text{mean},i}, \mathbf{r}^{\text{st}}_{\text{var},i}, \mathbf{r}^{\text{st}}_{\text{gate},i}\right), \tag{S15}$$

whereas the SM decoder produces the mean and variance parameters of the metabolomic observation model,

$$\mathcal{F}^{\text{sm}}_{\text{decoder}}\left(\mathbf{u}_i, \mathbf{B}_i\right) \to \left(\mathbf{r}^{\text{sm}}_{\text{mean},i}, \mathbf{r}^{\text{sm}}_{\text{var},i}\right). \tag{S16}$$

Here, $\mathbf{u}_i$ denotes the 10-dimensional latent representation supplied to the decoder: either the joint latent representation $\mathbf{z}_{\text{joint},i}$ or the corresponding modality-specific latent mean $\boldsymbol{\mu}_{\text{st},i}$ or $\boldsymbol{\mu}_{\text{sm},i}$. The latent vector is first projected to a 128-unit hidden layer with batch normalization (`BatchNorm1d`) and ReLU activation, followed by modality-specific output heads. For multi-section training, the one-hot tissue-section indicator $\mathbf{B}_i$ is concatenated to the latent representation before decoding.

Transcriptomic counts are modeled with a zero-inflated negative-binomial likelihood,

$$\mathcal{L}_{\text{recon}}^{\text{st}} = -\frac{1}{N} \sum_{i=1}^{N} \log p_{\text{ZINB}} \left( \mathbf{X}_i^{\text{st}} \mid \mathbf{r}_{\text{mean},i}^{\text{st}}, \mathbf{r}_{\text{var},i}^{\text{st}}, \mathbf{r}_{\text{gate},i}^{\text{st}} \right), \tag{S17}$$

whereas normalized, $\log(1+x)$-transformed, and scaled MSI intensities are modeled with a Gaussian likelihood,

$$\mathcal{L}_{\text{recon}}^{\text{sm}} = -\frac{1}{N} \sum_{i=1}^{N} \log p_{\mathcal{N}} \left( \mathbf{X}_i^{\text{sm}} \mid \mathbf{r}_{\text{mean},i}^{\text{sm}}, \mathbf{r}_{\text{var},i}^{\text{sm}} \right). \tag{S18}$$

Following the notation of SpatialMETA (Tian et al. 2025), the superscript cor denotes reconstruction from the corresponding modality-specific latent mean. For ST, this gives

$$\begin{aligned} \left( \mathbf{r}_{\text{mean},i}^{\text{st,cor}}, \mathbf{r}_{\text{var},i}^{\text{st,cor}}, \mathbf{r}_{\text{gate},i}^{\text{st,cor}} \right) &= \mathcal{F}_{\text{decoder}}^{\text{st}} \left( \boldsymbol{\mu}_{\text{st},i}, \mathbf{B}_i \right), \\ \mathcal{L}_{\text{recon}}^{\text{st,cor}} &= -\frac{1}{N} \sum_{i=1}^{N} \log p_{\text{ZINB}} \left( \mathbf{X}_i^{\text{st}} \mid \mathbf{r}_{\text{mean},i}^{\text{st,cor}}, \mathbf{r}_{\text{var},i}^{\text{st,cor}}, \mathbf{r}_{\text{gate},i}^{\text{st,cor}} \right). \end{aligned} \tag{S19}$$

Analogously, the corresponding SM reconstruction is

$$\begin{aligned} \left( \mathbf{r}_{\text{mean},i}^{\text{sm,cor}}, \mathbf{r}_{\text{var},i}^{\text{sm,cor}} \right) &= \mathcal{F}_{\text{decoder}}^{\text{sm}} \left( \boldsymbol{\mu}_{\text{sm},i}, \mathbf{B}_i \right), \\ \mathcal{L}_{\text{recon}}^{\text{sm,cor}} &= -\frac{1}{N} \sum_{i=1}^{N} \log p_{\mathcal{N}} \left( \mathbf{X}_i^{\text{sm}} \mid \mathbf{r}_{\text{mean},i}^{\text{sm,cor}}, \mathbf{r}_{\text{var},i}^{\text{sm,cor}} \right). \end{aligned} \tag{S20}$$

These corresponding reconstruction terms constrain the modality-specific latent means to remain directly decodable through the same modality-specific decoders used for the joint representation.

The variational posterior is regularized toward an isotropic standard normal prior through

$$\begin{aligned} \mathcal{L}_{\text{KL}} &= \frac{1}{N} \sum_{i=1}^{N} D_{\text{KL}} \left[ \mathcal{N} \left( \boldsymbol{\mu}_{\text{joint},i}, \operatorname{diag} \boldsymbol{\sigma}_{\text{joint},i}^2 \right) \middle\| \mathcal{N} \left( \mathbf{0}, \mathbf{I} \right) \right] \\ &= \frac{1}{2N} \sum_{i=1}^{N} \sum_{d=1}^{D} \left[ \mu_{\text{joint},id}^2 + \sigma_{\text{joint},id}^2 - \log \sigma_{\text{joint},id}^2 - 1 \right], \end{aligned} \tag{S21}$$

where $D$ is the latent dimensionality. This term limits departures of the joint posterior from the prior and promotes a smooth, regularized latent space.

For horizontal integration, SpatialMETA additionally matches the distributions of the joint posterior means across tissue sections:

$$\begin{aligned}\mathcal{L}_{\mathrm{MMD}} = \sum_{\substack{b,b'\in\mathcal{S}\\ b<b'}} \Bigg[ & \mathop{\mathbb{E}}_{i,j\sim\mathrm{Unif}(\mathcal{I}_b)} k\Big(\boldsymbol{\mu}_{\mathrm{joint},i}, \boldsymbol{\mu}_{\mathrm{joint},j}\Big) \\ & + \mathop{\mathbb{E}}_{i,j\sim\mathrm{Unif}(\mathcal{I}_{b'})} k\Big(\boldsymbol{\mu}_{\mathrm{joint},i}, \boldsymbol{\mu}_{\mathrm{joint},j}\Big) \\ & - 2 \mathop{\mathbb{E}}_{\substack{i\sim\mathrm{Unif}(\mathcal{I}_b)\\ j\sim\mathrm{Unif}(\mathcal{I}_{b'})}} k\Big(\boldsymbol{\mu}_{\mathrm{joint},i}, \boldsymbol{\mu}_{\mathrm{joint},j}\Big) \Bigg], \end{aligned} \tag{S22}$$

$$k(\mathbf{u},\mathbf{v}) = \sum_{\alpha\in\mathcal{A}} \exp\left(-\frac{\|\mathbf{u}-\mathbf{v}\|_2^2}{2\alpha}\right).$$

Here, $\mathcal{S}$ is the set of tissue sections, $\mathcal{I}_b$ contains the spots belonging to section $b$, and $\mathcal{A}$ is the set of bandwidths used by SpatialMETA's multiscale Gaussian kernel. The MMD term discourages section-specific shifts in the joint latent distribution; it is zero when training on a single section.

The original SpatialMETA objective can be written as

$$\begin{aligned}\mathcal{L}^{\mathrm{SpatialMETA}}_{\mathrm{original}} &= w_{\mathrm{st}}\,\mathcal{L}^{\mathrm{st}}_{\mathrm{recon}} + w_{\mathrm{sm}}\,\mathcal{L}^{\mathrm{sm}}_{\mathrm{recon}} \\ &\quad + w_{\mathrm{st,cor}}\,\mathcal{L}^{\mathrm{st,cor}}_{\mathrm{recon}} + w_{\mathrm{sm,cor}}\,\mathcal{L}^{\mathrm{sm,cor}}_{\mathrm{recon}} + \beta\,\mathcal{L}_{\mathrm{KL}} + w_{\mathrm{MMD}}\,\mathcal{L}_{\mathrm{MMD}}. \end{aligned} \tag{S23}$$

Here, in Eq. S23, $\mathcal{L}^{\mathrm{st}}_{\mathrm{recon}}$ and $\mathcal{L}^{\mathrm{sm}}_{\mathrm{recon}}$ are the ST and SM reconstruction losses from the joint latent representation $\mathbf{z}_{\mathrm{joint}}$, whereas $\mathcal{L}^{\mathrm{st,cor}}_{\mathrm{recon}}$ and $\mathcal{L}^{\mathrm{sm,cor}}_{\mathrm{recon}}$ reconstruct ST and SM from their respective modality-specific latent means, $\boldsymbol{\mu}_{\mathrm{st}}$ and $\boldsymbol{\mu}_{\mathrm{sm}}$. $\mathcal{L}_{\mathrm{KL}}$ regularizes the joint variational posterior toward a standard normal prior, and $\mathcal{L}_{\mathrm{MMD}}$ aligns the joint latent distributions across tissue sections during horizontal integration. Each $w$ is the weight of its corresponding reconstruction or alignment term, while $\beta$ weights the KL-divergence term.

We additionally introduced an ST→SM cross-reconstruction path in the SpatialMETA backbone used for the `SpatialJEPA` teacher. In the original SpatialMETA formulation, the SM decoder reconstructs metabolite measurements from either the joint latent representation or the SM-specific latent representation. We apply the same decoder to the ST-specific latent mean $\boldsymbol{\mu}_{\mathrm{st},i}$:

$$\mathcal{L}^{\mathrm{st}\to\mathrm{sm}}_{\mathrm{recon}} = -\frac{1}{N}\sum_{i=1}^{N} \log p_{\mathcal{N}}\left(\mathbf{X}^{\mathrm{sm}}_i \mid \mathbf{r}^{\mathrm{st}\to\mathrm{sm}}_{\mathrm{mean},i}, \mathbf{r}^{\mathrm{st}\to\mathrm{sm}}_{\mathrm{var},i}\right), \tag{S24}$$

where

$$\left(\mathbf{r}^{\mathrm{st}\to\mathrm{sm}}_{\mathrm{mean},i}, \mathbf{r}^{\mathrm{st}\to\mathrm{sm}}_{\mathrm{var},i}\right) = \mathcal{F}^{\mathrm{sm}}_{\mathrm{decoder}}\left(\boldsymbol{\mu}_{\mathrm{st},i}, \mathbf{B}_i\right). \tag{S25}$$

This term introduces no additional decoder parameters: it reuses the same SM decoder and Gaussian likelihood as the original SpatialMETA reconstruction terms, but uses the ST-specific latent representation as the input. It therefore explicitly trains the RNA-derived latent representation to retain information predictive of the metabolomic modality.

**Table S1:** Loss weights used for SpatialMETA teacher training. The weights for the original SpatialMETA loss terms follow the published single-section and horizontal-integration presets (Tian et al. 2025). The added ST→SM cross-reconstruction term uses the same weight as the existing SM reconstruction terms in each training regime.

| Term | Weight | Mouse (1 section) | Human (3 sections) |
|---|---|---|---|
| ST reconstruction from joint latent | $w_{\mathrm{st}}$ | 5 | 8 |
| SM reconstruction from joint latent | $w_{\mathrm{sm}}$ | 1 | 2 |
| ST reconstruction from ST latent | $w_{\mathrm{st,cor}}$ | 5 | 8 |
| SM reconstruction from SM latent | $w_{\mathrm{sm,cor}}$ | 1 | 2 |
| ST→SM reconstruction | $w_{\mathrm{st}\to\mathrm{sm}}$ | 1 | 2 |
| KL divergence | $\beta$ | 0.5 | 1 |
| Cross-section MMD | $w_{\mathrm{MMD}}$ | — | 10 |

The final SpatialMETA backbone objective adds the ST→SM cross-reconstruction loss from Eq. S24 to the original SpatialMETA loss:

$$\mathcal{L}_{\mathrm{backbone}}^{\mathrm{SpatialMETA}} = \mathcal{L}_{\mathrm{original}}^{\mathrm{SpatialMETA}} + w_{\mathrm{st}\to\mathrm{sm}}\,\mathcal{L}_{\mathrm{recon}}^{\mathrm{st}\to\mathrm{sm}}. \tag{S26}$$

SpatialMETA loss weights for the mouse and human models, including the added ST→SM cross-reconstruction term, are summarized in Table S1. The added ST→SM term uses the same weight as the existing SM reconstruction terms: 1 for the single-section mouse model and 2 for the multi-section human model.

#### S9.6 SpatialMETA training schedules and teacher early stopping

Training used mini-batches of 128 spots, with one student step following each teacher batch. The teacher was optimized with AdamW (weight decay $10^{-6}$) and the student with Adam (learning rate $10^{-3}$, weight decay $10^{-4}$). The KL weight $\beta$ was warmed up linearly from zero over the first $\min(400, T)$ epochs of a $T$-epoch run.

For training across multiple tissue sections, graphs were constructed separately within each slide capture area and combined block-diagonally, preventing edges between spots from different sections. Because the encoder operates on mini-batches of 128 nodes, the normalized neighborhood aggregation $\hat{\mathbf{A}}\mathbf{X}$ of the encoder input was precomputed once over the full graph and gathered for the spots of each batch, so that mini-batch outputs equal those of the full-graph convolution (Fey and Lenssen 2019).

The mouse model (tissue section V11L12-109_B1) used a teacher learning rate of $10^{-3}$, decayed by a factor of ten every 200 epochs for both teacher and student, the single-section loss weights in Table S1, and a 400-epoch KL warm-up. The teacher was early-stopped. A selection run trained the teacher alone on 8 of 10 spatially contiguous blocks of the section ($k$-means on spot coordinates), evaluated the training objective at the final KL weight on the remaining two blocks after every epoch, and stopped 100 epochs after its last improvement. The teacher was then trained on all spots for the selected number of epochs (29), keeping the warm-up schedule of the full run, and the student continued distilling from the frozen teacher for $T = 1{,}000$ epochs (Section S6).

The human model (tissue sections V11T17-102_A1, _C1 and _D1) was trained jointly at a constant teacher learning rate of $10^{-5}$, with a 400-epoch KL warm-up. Multi-section training used section-conditioned decoders, the block-diagonal teacher graph, and the cross-section MMD penalty. The teacher was early-stopped as for the mouse model, with 2 of 10 spatially contiguous blocks held out in each section. The selection run stopped at epoch 370, the teacher was trained on all spots for the selected 270 epochs, and the student continued distilling from the frozen teacher for $T = 1{,}000$ epochs. Loss weights for the multi-section model are reported in Table S1.

##### S9.7 Post-hoc PLS decoders

In both the RNA–ATAC and RNA–MSI settings, PLS is fitted by the covariance criterion described below and is not part of the backbone reconstruction or distillation objectives. The ST→SM cross-reconstruction term encourages the RNA-derived representation to retain metabolite-related information during training, whereas the post-hoc PLS decoder maps that representation to individual metabolite features.

All post-hoc decoders in the main text were fitted as $K$-component PLS regressions (Tsuchiya et al. 2022). Let $\mathbf{X} \in \mathbb{R}^{N\times D}$ denote the $D$-dimensional embeddings of $N$ observations and let $\mathbf{Y} \in \mathbb{R}^{N\times F}$ denote the corresponding $F$ measured molecular features. Before fitting, the PLS implementation centers and scales each column of both matrices, giving

$$\mathbf{X}_0 = (\mathbf{X} - \mathbf{1}_N\bar{\mathbf{x}}^\top)\mathbf{S}_X^{-1}, \qquad \mathbf{Y}_0 = (\mathbf{Y} - \mathbf{1}_N\bar{\mathbf{y}}^\top)\mathbf{S}_Y^{-1}, \tag{S27}$$

where $\bar{\mathbf{x}}$ and $\bar{\mathbf{y}}$ are the column means and $\mathbf{S}_X$ and $\mathbf{S}_Y$ are diagonal matrices of column standard deviations.

PLS regression performs supervised dimensionality reduction by extracting components that maximize predictor–response covariance rather than predictor variance alone. Let $\mathbf{X}_{k-1}$ and $\mathbf{Y}_{k-1}$ denote the residual predictor and response matrices after extraction of the first $k-1$ components. The $k$th predictor weight is chosen as

$$\mathbf{w}_k = \underset{\|\mathbf{w}\|_2=1}{\arg\max} \left\| \mathbf{Y}_{k-1}^\top \mathbf{X}_{k-1}\mathbf{w} \right\|_2^2, \qquad \mathbf{t}_k = \mathbf{X}_{k-1}\mathbf{w}_k, \tag{S28}$$

where $\mathbf{t}_k$ is the predictor score for component $k$. The corresponding predictor and response loadings are obtained by least squares,

$$\mathbf{p}_k = \frac{\mathbf{X}_{k-1}^\top \mathbf{t}_k}{\mathbf{t}_k^\top \mathbf{t}_k}, \qquad \mathbf{q}_k = \frac{\mathbf{Y}_{k-1}^\top \mathbf{t}_k}{\mathbf{t}_k^\top \mathbf{t}_k}, \tag{S29}$$

after which both matrices are deflated as

$$\mathbf{X}_k = \mathbf{X}_{k-1} - \mathbf{t}_k\mathbf{p}_k^\top, \qquad \mathbf{Y}_k = \mathbf{Y}_{k-1} - \mathbf{t}_k\mathbf{q}_k^\top. \tag{S30}$$

Repeating this procedure yields $K$ mutually orthogonal predictor-score vectors and a supervised rank-$K$ representation whose components are selected for covariance with the molecular response.

For a fitted $K$-component model, let $\mathbf{W}, \mathbf{P} \in \mathbb{R}^{D\times K}$ denote the predictor weights and loadings, respectively. Because the columns of $\mathbf{W}$ are learned on successively deflated predictor matrices, they do not directly define the projection from the original standardized matrix $\mathbf{X}_0$ to all component scores. This projection is instead given by the PLS rotation matrix

$$\mathbf{R} = \mathbf{W}(\mathbf{P}^\top \mathbf{W})^{-1} \in \mathbb{R}^{D\times K}, \tag{S31}$$

which projects each embedding onto its $K$ PLS component scores,

$$\mathbf{T} = \mathbf{X}_0 \mathbf{R} \tag{S32}$$

If $\mathbf{Q} \in \mathbb{R}^{F\times K}$ contains the corresponding response-side feature loadings, the centered predicted feature matrix is

$$\widehat{\mathbf{Y}}_0 = \mathbf{X}_0 \mathbf{R} \mathbf{Q}^\top = \mathbf{X}_0 \mathbf{B}, \qquad \mathbf{B} = \mathbf{R}\mathbf{Q}^\top. \tag{S33}$$

Thus, each decoder first projects the $D$-dimensional embedding onto $K$ component scores and then expands those scores onto the $F$ measured molecular features. Finally, the predictions are translated back to the response scale as $\widehat{\mathbf{Y}} = \mathbf{1}_N \bar{\mathbf{y}}^\top + \widehat{\mathbf{Y}}_0 \mathbf{S}_Y$. The columns of $\mathbf{T}$ are the per-observation PLS component scores, whereas the columns of $\mathbf{Q}$ describe the molecular features associated with each component.

The RNA–ATAC analyses use separate post-hoc PLS regressions to relate the fixed student embeddings to measured RNA or ATAC features (Fig. S18). The resulting component scores and feature loadings are used to compare molecular structure across source and target representations, with RNA and ATAC decoded independently.

##### S9.8 Teacher-side metabolite prediction for spatial targets

For the spatial P22 RNA–ATAC target, we generated MSI predictions directly with the trained SpatialMETA teacher. The teacher decodes the ST-specific latent mean $\boldsymbol{\mu}_{\mathrm{st}}$ through the SM decoder, using the ST→SM prediction path supervised during training by the cross-reconstruction term $\mathcal{L}^{\mathrm{st}\to\mathrm{sm}}_{\mathrm{recon}}$. Predicted metabolite values were taken from the Gaussian mean output of the decoder. Because this route requires the spatial graph, teacher-side metabolite prediction is restricted to spatially resolved targets.

##### S9.9 Representation benchmarks

**Baselines and ablations** We evaluated `SpatialJEPA` against both matched internal ablations and external multimodal-integration baselines. The non-spatial ablations described in main-text Methods (*MultiGATE backbone* and *SpatialMETA distillation and training*) are the most direct: they retain the same backbone, training data, and optimization schedule as the corresponding

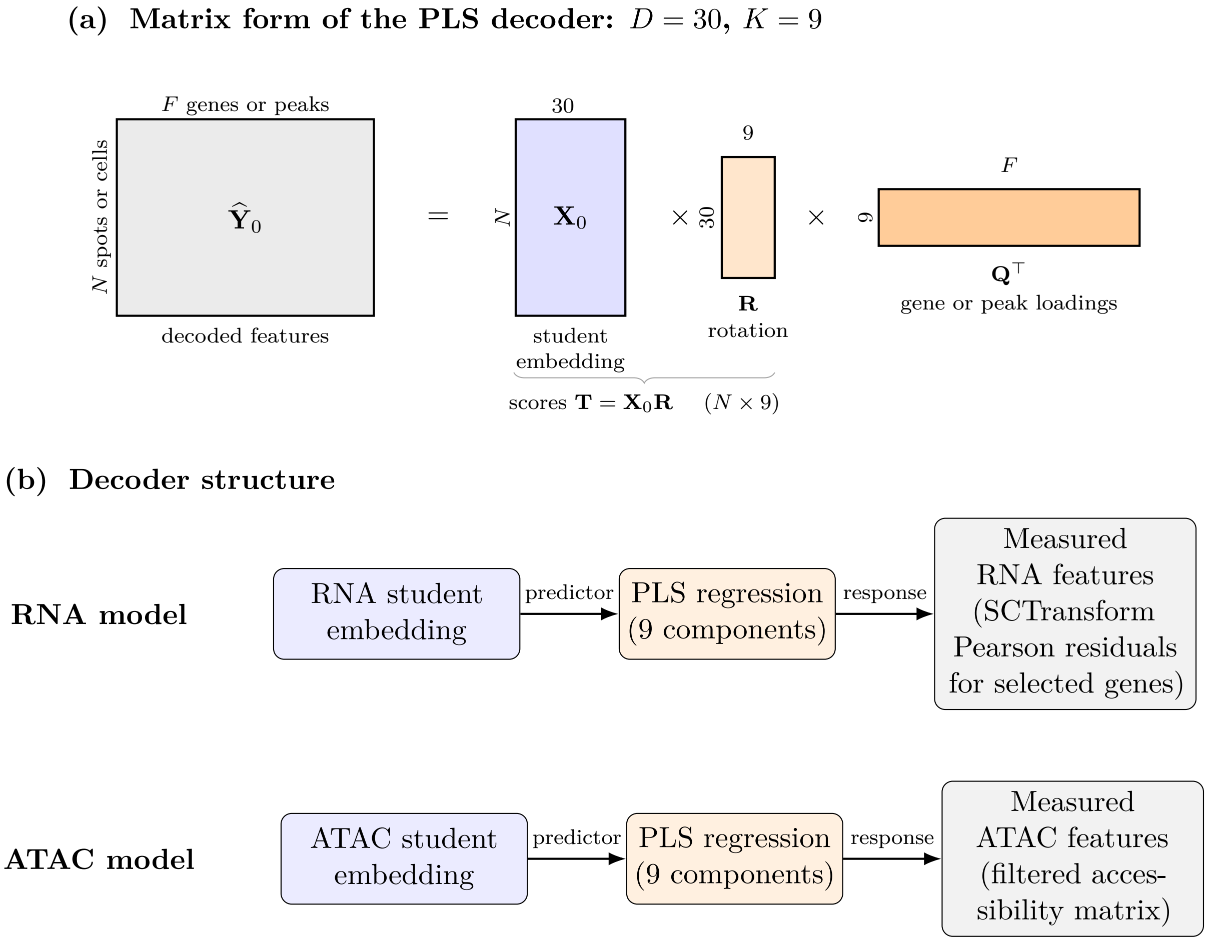

**Figure S18:** Post-hoc PLS decoders used for interpretation of RNA–ATAC embeddings. Separate nine-component PLS regressions were fitted for RNA and ATAC after graph-context distillation, with the corresponding student embedding used as predictor and the measured molecular features used as response. For RNA, the response matrix consisted of SCTransform Pearson residuals for the selected genes; for ATAC, it consisted of the filtered accessibility matrix. These linear decoders are fitted after distillation and are distinct from the backbone decoders used during model training. **(a)** Shared matrix form of the RNA and ATAC PLS decoders. The centered predicted feature matrix is factorized as $\widehat{\mathbf{Y}}_0 = \mathbf{X}_0\mathbf{R}\mathbf{Q}^\top$: the thirty-dimensional student embedding $\mathbf{X}_0$ is projected through the PLS rotation $\mathbf{R}$ onto nine component scores $\mathbf{T} = \mathbf{X}_0\mathbf{R}$, which are then mapped onto the $F$ retained genes or peaks through the response loadings $\mathbf{Q}^\top$. Each row of $\mathbf{X}_0$ is the embedding of one spot or cell, standardized column-wise as in Eq. S27. **(b)** Modality-specific decoder structure. RNA and ATAC are decoded independently, with the corresponding student embedding used to predict the measured features of that modality. The metabolite decoder of Fig. S19c, fitted on each RNA-only target, has the same matrix form.

`SpatialJEPA` models, but remove the spatial graph and teacher supervision.

For the RNA–ATAC experiments, we used MultiVI (Ashuach et al. 2023) as an external baseline. A MultiVI model was trained on raw source counts using the `scvi-tools` default architecture and latent width for 3,000 epochs, matching the corresponding MultiGATE models. From this model, we extracted the joint, RNA-only, and ATAC-only posterior-mean representations. Architecture and optimization were left at the published `scvi-tools` defaults. To provide a zero-shot comparison on the target, the source-trained MultiVI model was also applied directly to the target without retraining; this is possible because source and target already share a common feature index.

For the RNA–MSI experiments, we used Multigrate (Litinetskaya et al. 2025), trained on the same 2,526 registered spots from section V11L12-109_B1 with a negative-binomial likelihood for raw RNA counts and a mean-squared-error likelihood for continuous MSI intensities. The Multigrate model used a 16-dimensional latent space and was trained with 1,689 RNA features and the same 800 MSI features used in the SpatialMETA analysis. The larger RNA feature set reflects an earlier preprocessing stage, whereas the final SpatialMETA feature space was further restricted for compatibility with the downstream transfer datasets as described in main-text Methods (*RNA–MSI preprocessing and feature selection*).

**Conventional integration metrics** Cross-modal pairing, modality mixing, and preservation of annotated biological structure were quantified by $1 - \text{FOSCTTM}$ (fraction of samples closer than the true match), the integration local inverse Simpson's index (iLISI), and a composite biological-conservation score, respectively, using the scIB metrics implementation (Luecken et al. 2022; Niu et al. 2025). Metrics were computed on the modality-specific RNA and ATAC embeddings, and label-dependent metrics used a single annotation per paired observation: the RNA cluster of each spot on the spatial source and the CellTypist prediction of each nucleus on the dissociated target (main-text Methods, *Datasets and preprocessing*). Larger values indicate better performance for all three metrics. Metric definitions, uncertainty estimation, and the distinction between in-sample and transfer evaluation are given in Section S1.

**Spatial-structure metrics** Because the integration metrics above do not account for tissue geometry, we evaluated spatial-structure transfer separately using three continuous readouts on the spatial source, where both the teacher and matched non-spatial ablation are available. In these evaluations, spatial coordinates were used to define prediction targets, physical distances, and cross-validation folds, but were not provided as inputs to the downstream readouts. The encoders were trained on all source spots, with the teacher using a coordinate-derived spatial graph. Thus, spatial-block cross-validation held out observations from readout fitting, not encoder training. Before evaluation, each embedding dimension was standardized to place distances and ridge-regression fits on a comparable scale across models. The three metrics were evaluated using ten spatial blocks. The section was partitioned into ten spatially compact blocks by $k$-means clustering of spot coordinates. Spatially separated folds reduce the optimistic bias that can arise when spatially autocorrelated observations are split at random (Roberts et al. 2017; Valavi et al. 2018). Niche recovery and orthogonal-target recovery used spatial-block cross-validation, with each block held out in turn during readout fitting.

*Niche recovery* predicts, from the joint embedding, the cluster composition of each spot's fifteen nearest physical neighbors excluding itself. Representing local tissue context through the composition of spatial neighborhoods is widely used in spatial-omics analysis (Palla et al. 2022; Mason et al. 2024). The one-hot RNA and ATAC cluster vectors of these neighbors were averaged and concatenated into a single multi-output target.

Ridge regression with a fixed penalty of $\alpha = 1$ was fitted on each training fold after standardizing predictor dimensions using that fold alone, and then applied to the held-out block. Predictions were scored by the mean Spearman correlation across output columns, which is robust to the scale and mean shifts induced by spatial-block cross-validation. The same fixed penalty and readout procedure were used for all models rather than tuning the downstream decoder separately for each representation.

*Orthogonal-target recovery* predicts, from the RNA-only embedding, the neighbor-averaged 30-component truncated-SVD representation of ATAC accessibility under the same cross-validation scheme. It therefore tests whether the representation of one modality preserves spatial structure expressed in the other. This follows the broader principle of evaluating spatially informed representations by their ability to recover molecular or neighborhood context not directly supplied to the representation (Haviv et al. 2025).

*Within-cluster coherence* measures whether embedding distances track physical distances among spots assigned to the same transcriptomic or chromatin cluster. Within each spatial block, up to 2,000 spot pairs were randomly sampled from each cluster containing at least eight spots. Spearman correlations between embedding and physical distances were computed per cluster and averaged across both partitions, weighted by cluster size, yielding one score per block. Unlike the other two metrics, coherence requires no fitted readout. Conditioning on cluster reduces confounding by cell identity, as spatially organized cell types can produce apparent neighborhood agreement even without recovery of finer tissue geometry.

For each metric, 95% confidence intervals were obtained by bootstrapping the ten per-block scores over 2,000 replicates, and the full model lineup was re-evaluated after permutation of the spatial coordinates as a negative control. The RNA–MSI analyses used the same definitions on the spatial RNA–MSI section, with MSI cluster labels and neighbor-averaged MSI truncated-SVD components substituted for their ATAC counterparts.

A complementary fourth readout, held-out position recovery, does not fit a predictor and is defined in Section S9.10.

**Placement of held-out Multiome nuclei on the spatial source** For each model, Multiome nuclei were coupled to P22 spots in that model's embedding of both datasets, with the RNA and ATAC embeddings concatenated after scaling each to unit mean-square norm on P22. Two couplings were compared: each spot's 15 nearest nuclei (15-NN), and balanced entropic optimal transport with the Euclidean cost scaled by its mean and $\varepsilon = 0.01$, as in main-text Methods (*Zero-shot transfer and optimal-transport alignment*). Teacher clusters were obtained by Leiden clustering ($K = 5$, 10, and 20 clusters) of the spatial teacher's P22 embedding, on a 150-nearest-neighbor graph with the resolution chosen to give exactly $K$ clusters. Each model's own Leiden clusters of its Multiome embedding, computed in the same way before alignment, were carried to the spots by majority vote (15-NN) or by the largest transported mass (OT), and agreement with the teacher clusters was measured by ARI averaged over $K$. As in-sample references, each model's P22 embedding was clustered in the same way. For main-text Fig. 2g, placed clusters ($K = 10$, 15-NN) were matched one-to-one to the teacher clusters by maximal overlap (Hungarian algorithm). The models were the student; the teacher evaluated with an identity graph; the non-spatial MultiGATE ablation; and P22-trained MultiVI applied zero-shot. The non-spatial ablation and the graph-free teacher were encoded on the Multiome for this analysis; the same encoding reproduces the manuscript's student, teacher, and ablation embeddings to within $10^{-6}$.

**Dopamine-embedding benchmark** To assess whether spatial context improves the representation of biologically structured metabolite variation, we compared how measured dopamine is organized across latent spaces learned from the spatial RNA–MSI source section. Here, measured dopamine was used to assess whether nearby profiles in each latent space had similar dopamine levels, rather than whether dopamine could be predicted from the embedding. We compared the spatial teacher, graph-masked student, and non-spatial SpatialMETA ablation with Multigrate (Litinetskaya et al. 2025) as an external non-spatial multimodal-integration reference. All readouts were computed from $k$-nearest-neighbor graphs constructed in the joint latent spaces. For comparability across models, representations wider than 10 dimensions were first reduced to 10 dimensions by PCA, reducing the 16-dimensional Multigrate representation to the native dimensionality of the SpatialMETA models. Latent dimensions were then standardized to zero mean and unit variance.

Given the strongly asymmetric dopamine signal between intact and lesioned striatum in the unilateral lesion model, we evaluated three complementary measures of dopamine organization using $k = 15$, consistent with representation-learning benchmarks that assess biologically meaningful neighborhood structure (Luecken et al. 2022; Zhong et al. 2024; Zhu et al. 2026).

For each spot $i$, let $\bar{d}_i$ denote the mean measured dopamine of its $k$ nearest neighbors in latent space. The kNN graph was constructed separately for each analysis: using striatal spots for structural gain, each tissue subset for Moran's $I$, and the complete section for intact-striatal specificity. *Structural gain* is the change in intact-versus-lesioned discrimination when measured dopamine $\mathbf{d}$ is replaced by its neighbor-smoothed value $\bar{\mathbf{d}}$, $\Delta\text{AUROC} = \text{AUROC}(\bar{\mathbf{d}}, \mathbf{y}) - \text{AUROC}(\mathbf{d}, \mathbf{y})$, where $\mathbf{y}$ indicates intact-striatal membership; positive values indicate that latent-neighborhood smoothing improves the separation of intact and lesioned striatum. *Latent-graph Moran's $I$* measures the autocorrelation of measured dopamine on the latent-neighborhood graph, calculated separately within intact and lesioned striatum and within the corresponding non-striatal regions. *Intact-striatal specificity* is the log-odds of intact-striatal membership among spots in the top decile of neighbor-smoothed dopamine, relative to the section-wide log-odds. Formal definitions, including the pseudocount that bounds saturated bootstrap estimates, are given in Section S9.12.

Because all four model representations were available for only one tissue section, uncertainty was estimated by within-section spatial-block bootstrap rather than by resampling independent sections. For the striatum-restricted structural-gain and Moran's $I$ analyses, striatal spots were partitioned into 25 spatial blocks by $k$-means clustering of their coordinates, and whole blocks were resampled with replacement for 1,000 paired bootstrap replicates. Intact-striatal specificity, which is defined over the complete tissue section, was bootstrapped separately using the same procedure applied to all spots. The same resampled blocks were used across models within each replicate, enabling paired comparisons with the non-spatial SpatialMETA ablation. These intervals therefore quantify spatial variability within the analyzed section and should not be interpreted as estimates of variability across independent tissue sections.

#### S9.10 Spatial position recovery

Position recovery was evaluated on the spatial source section using only spots shared by all methods under comparison. For each of 40 random half-splits, half of the spots were held out as queries and the remainder used as references. Each query was matched to its nearest reference by Euclidean distance in representation space, and localization error was defined as the physical distance between their coordinates, normalized by the median nearest-neighbor spacing of the tissue lattice (one spot width). Per-spot hit rates were defined as the fraction of query appearances in which the inferred position fell within two spot widths of the true position. The random baseline was generated from physical distances between independently and uniformly sampled pairs of source spots, using the same number of distances as the held-out evaluation. Because spots are held out only during evaluation, this analysis measures how much positional information is recoverable from a fitted representation rather than generalization to previously unseen tissue.

Four learned representations were evaluated with the same estimator: MultiVI, the non-spatial MultiGATE ablation, the graph-masked `SpatialJEPA` student with an identity graph at inference, and the spatial `SpatialJEPA` teacher. Each was scored on its joint RNA–ATAC latent: for the three MultiGATE-based models, the mean of the $\ell_2$-normalized RNA and ATAC embeddings, MultiGATE's joint embedding; for MultiVI, its joint latent. Fig. S3 repeats the evaluation on the RNA-side and ATAC-side latents separately. As an additional molecular reference, the same position-recovery estimator was applied to the first 30 principal components of the log-normalized RNA expression matrix, testing how much positional information was recoverable directly from transcriptomic variation. Because the teacher directly aggregates information over spatial neighborhoods, it serves as a spatial-context reference rather than a competing graph-free representation.

To summarize localization across the full error distribution, we constructed for each spot an empirical cumulative distribution function (ECDF) of localization error and calculated the area under this curve on the logarithmic distance axis used in Fig. S2b. For errors $e_1, \ldots, e_Q$ accumulated across the $Q$ query appearances of a spot and the maximum error $E$ observed across all model arms

($E \approx 129$ spot widths here, which also sets the upper limit of the axis in Fig. S2b), the ECDF area is

$$A = \frac{1}{Q}\sum_{i=1}^{Q}\left(1 - \frac{\log_{10}\max(e_i, 1)}{\log_{10} E}\right), \tag{S34}$$

with larger values indicating more precise localization. The same value of $E$ was used for every representation to ensure directly comparable ECDF areas, and a chance level was obtained by applying the same statistic to uniformly random spot pairs.

##### S9.11 Domain-level neighborhood agreement

To examine whether the teacher and student represented local tissue organization similarly across anatomical domains, we compared their latent-space neighborhoods with physical neighborhoods. For each spot, neighborhood agreement was defined as the fraction of its $k$ nearest neighbors in representation space that were also among its $k$ nearest spatial neighbors, with $k = 10$. Agreement was averaged within each of the eleven published RNA-annotated domains, and the student-minus-teacher difference was computed for each domain. Values closer to zero indicate more similar neighborhood agreement between the two models.

We then asked whether variation in this difference was associated with tissue-domain organization. Neighborhood contiguity was defined as the mean fraction of each spot's six nearest spatial neighbors carrying the same domain label. Its association with the domain-level student-minus-teacher difference was assessed by Spearman correlation with a two-sided test.

##### S9.12 Dopamine-embedding benchmark definitions

The three dopamine-embedding readouts of main-text Fig. 4d–f are defined as follows, with $k = 15$ latent nearest neighbors.

Let $y_i \in \{0, 1\}$ indicate intact-striatal membership, $d_i$ the measured dopamine intensity at spot $i$, and $\mathcal{N}_k(i)$ the $k$ nearest neighbors of $i$ in latent space. The kNN graph was constructed separately for each analysis: using striatal spots for structural gain, each tissue subset for Moran's $I$, and the complete section for intact-striatal specificity. We defined the neighbor-smoothed dopamine value as

$$\bar{d}_i = \frac{1}{|\mathcal{N}_k(i)|}\sum_{j \in \mathcal{N}_k(i)} d_j.$$

Structural gain quantifies the change in intact-versus-lesioned discrimination induced by this latent-neighborhood smoothing:

$$\Delta\mathrm{AUROC} = \mathrm{AUROC}\left(\bar{\mathbf{d}}, \mathbf{y}\right) - \mathrm{AUROC}(\mathbf{d}, \mathbf{y})\,. \tag{S35}$$

Positive values indicate that neighborhood smoothing in the latent space improves separation of intact and lesioned striatum relative to the measured dopamine signal alone. Because both terms are computed from the same measured dopamine values and lesion labels, their difference isolates the effect of the latent-neighborhood structure on this discrimination. *Latent-graph Moran's I* measures spatial autocorrelation of the measured dopamine values on the latent-neighborhood graph. It was calculated separately within intact and lesioned striatum and within the corresponding non-striatal regions, with the graph reconstructed independently within each subset. *Intact-striatal specificity* asks whether spots assigned to dopamine-high latent neighborhoods preferentially occur in intact striatum. We defined the dopamine-high set as the top decile of neighbor-smoothed dopamine values,

$$\mathcal{T} = \left\{ i : \bar{d}_i \geq Q_{0.9}\left(\bar{\mathbf{d}}\right) \right\},$$

and let

$$\hat{p} = \frac{|\{i \in \mathcal{T} : y_i = 1\}|}{|\mathcal{T}|}$$

denote the fraction of these spots belonging to intact striatum. Specificity was then reported as the log-odds ratio relative to the section-wide fraction $b$ of spots belonging to intact striatum,

$$\log \frac{\hat{p} + \varepsilon}{1 - \hat{p} + \varepsilon} - \log \frac{b + \varepsilon}{1 - b + \varepsilon}, \tag{S36}$$

where $\varepsilon = 10^{-9}$ prevents undefined values at the boundaries. When bootstrap resamples yield complete intact-striatal enrichment ($\hat{p} = 1$), this pseudocount imposes a finite numerical ceiling on the log-odds, producing saturated upper confidence bounds. These bounds therefore reflect complete enrichment in some resamples rather than precisely estimated effect sizes. Larger values indicate greater enrichment of intact-striatal spots among dopamine-high neighborhoods relative to their section-wide prevalence.

##### S9.13 Analysis of transferred RNA–ATAC representations

**Marker-gene scores and source–target gene-rank concordance** For visualization of canonical tissue programs, marker-gene expression scores were computed separately in source and target using `scanpy.tl.score_genes`. This score compares the mean expression of the specified marker genes with that of expression-matched control genes. We used *Pde10a*, *Rgs9*, and *Gng7* for the striatal score and *Mobp* and *Mal* for the oligodendrocyte/white-matter score.

For main-text Fig. 3a,b, Leiden clusters were computed separately in source and OT-aligned target using the stacked RNA and ATAC student embeddings ($k = 150$, resolution 0.9). We evaluated shared Leiden parameters across a grid of neighborhood sizes ($k \in \{15, 30, 50, 100, 150\}$) and resolutions (0.3–1.0), selecting $k = 150$ and resolution 0.9 based on source–target cluster agreement (adjusted Rand index) among configurations yielding equal numbers of clusters (6–10). Target clusters were matched one-to-one to source clusters by maximizing overlap with nearest-source labels using the Hungarian algorithm (`scipy.optimize.linear_sum_assignment`), without altering cluster membership.

To assess source–target transcriptomic concordance within matched Leiden clusters, genes distinguishing each cluster from all remaining observations were identified independently in source and target using the `scanpy.tl.rank_genes_groups` $t$-test. Genes with Benjamini–Hochberg-adjusted $p < 0.05$ in both domains were retained. For main-text Fig. 3b, genes were ranked by $-\log(p_{\mathrm{adj}})$, such that higher ranks indicate stronger differential-expression significance, and source–target concordance was quantified by Spearman correlation of these ranks within each matched cluster pair.

**PLS decoding of RNA–ATAC embeddings** For the RNA–ATAC analyses, separate post-hoc PLS decoders were fitted for RNA and ATAC (Fig. S18). The RNA decoder mapped the RNA embedding to measured gene expression, whereas the ATAC decoder mapped the ATAC embedding to measured chromatin accessibility. In the notation of Eq. S33, both used $D = 30$ embedding dimensions and retained $K = 9$ PLS components, with $F$ equal to the number of retained genes or peaks. For the spatial source, $\mathbf{X}$ was the corresponding student embedding; for the non-spatial target, $\mathbf{X}$ was the OT-aligned student embedding described in main-text Methods (*Zero-shot transfer and optimal-transport alignment*). The RNA response matrix $\mathbf{Y}$ contained SCTransform Pearson residuals (Hafemeister and Satija 2019) for the selected genes, whereas the ATAC response contained the filtered accessibility values.

The resulting score matrix $\mathbf{T}$ gives the nine component values for each spot or cell, which were used for the spatial maps in main-text Fig. 3c. The corresponding response loadings $\mathbf{Q} \in \mathbb{R}^{F\times K}$ describe the $F$ genes or peaks associated with each of the $K$ components. Source and target PLS models were fitted independently and their components subsequently compared across domains as described under *Cross-domain PLS component matching.* The target-side PLS fitting is a post-hoc analysis of the transferred representation and does not involve retraining the student encoder on the target data.

**Cross-domain PLS component matching** Because the source and target PLS models were fitted independently for RNA and ATAC, component signs were aligned before comparison. We first oriented source RNA PLS 0 toward positive association with the oligodendrocyte/myelin marker program. A common matrix formed by concatenating the source-aligned student embeddings from the four source/target modality views was then passed through each fitted PLS model. For each of the first five components, signs were chosen to maximize the sum of pairwise Pearson correlations between component scores, with the source RNA sign held fixed. Absolute Pearson correlations between source and target response loadings $\mathbf{Q}^S$ and $\mathbf{Q}^T$ (Eq. S33; extracted as `y_weights_`) were computed across shared genes for RNA and shared peaks for ATAC to examine cross-domain correspondences. For spatial visualization and RNA gene-set enrichment (main-text Fig. 3c,d), source and target components were compared at their original indices (PLS 0–4), without reordering.

**Module-loading enrichment** Component loadings were tested against six fixed marker modules of six to seven canonical genes each, specified in advance from established mouse brain cell-type markers and not derived from any model output: oligodendrocyte/myelin (*Mobp*, *Mal*, *Mag*, *Mbp*,

*Plp1*, *Cnp*), striatal MSN signaling (*Pde10a*, *Rgs9*, *Gng7*, *Ppp1r1b*, *Drd1*, *Adora2a*), mural/smooth muscle (*Cald1*, *Myh11*, *Mylk*, *Acta2*, *Tagln*, *Rgs5*, *Pdgfrb*), endothelial/BBB (*Flt1*, *Abcb1a*, *Slco1c1*, *Kdr*, *Cldn5*, *Pecam1*), astrocyte (*Aqp4*, *Slc1a3*, *Aldoc*, *Gfap*, *S100b*, *Glul*), and ECM/adhesion (*Col1a1*, *Col1a2*, *Col3a1*, *Col5a1*, *Dcn*, *Fn1*). For each component, gene loadings were standardized across the full feature set, and the mean standardized loading of each module was compared with 2,000 randomly sampled gene sets of matched size. The resulting null-calibrated score was

$$z = \frac{\bar{\beta}_{\text{obs}} - \mu_{\text{null}}}{\sigma_{\text{null}}}, \tag{S37}$$

with two-sided empirical $p$-values obtained from the permutation distribution. The 60 tests across six modules, five same-index component pairs, and two domains were corrected jointly by the Benjamini–Hochberg procedure, with FDR $< 0.05$ considered significant.

**Gene–peak link extraction and PLS weighting** Gene–peak links were identified from MultiGATE's learned gene–peak attention weights. The candidate feature graph connected selected ATAC peaks lying within 150 kb of a gene promoter, with distance-dependent edge weights. The first-layer gene–peak attention matrix was extracted from the trained student and mapped back to peak intervals and gene symbols. Under the feature-anchored attention parameterization, attention scores depend on learned per-feature parameters and the distance prior rather than on the input profiles. Consequently, the attention scores and retained candidate links are identical in source and target. Because attention coefficients are normalized over each feature's neighbors, the coefficient $a_{pg}$ represents the relative attention assigned from peak $p$ to candidate gene $g$ and should not be interpreted as a signed regulatory effect.

To retain high-attention links, the attention coefficients were thresholded using a two-component Gaussian mixture model, following MultiGATE's documentation. Candidate pairs with attention above the intersection of the fitted component densities were retained. Retained peaks were annotated using GENCODE coordinates and their distance to the corresponding transcription start site. Links within 2 kb were classified as promoter-proximal and the remainder as distal.

For each gene $g$, retained peaks were summarized into an attention-weighted accessibility score,

$$s_{i,g} = \sum_{p \in \mathcal{P}_g} x^{A}_{i,p} \, a_{pg}, \tag{S38}$$

where $\mathcal{P}_g$ is the attention-thresholded set of peaks linked to gene $g$, and $x^{A}_{i,p}$ is the accessibility of peak $p$ in observation $i$. Attention therefore determines both link membership and each peak's contribution to this non-negative summary.

**Spatial coupling of expression and linked-peak accessibility** For each gene, the spatial coupling between its expression and the accessibility of its linked peaks, $s_{\cdot,g}$ (Eq. S38), was compared between source and aligned target data on a common spatial lattice. Because target nuclei inherit source coordinates after alignment and may map many-to-one onto source spots, target RNA and ATAC profiles were averaged within their assigned source spot, separately for each modality, and both domains were restricted to the $n = 8{,}945$ source spots occupied by both target modalities. Let $\mathbf{z}_g^R$ and $\mathbf{z}_g^A$ denote the expression of gene $g$ and its linked-peak accessibility across these spots, each standardized to zero mean and unit variance, and let $W$ be a six-nearest-neighbor spatial-weight matrix whose adjacency was symmetrized and then row-standardized. The bivariate Moran's $I$ (Eq. S40 with row-standardized weights, for which $S_0 = n$) is

$$I_g = \frac{1}{n} (\mathbf{z}_g^R)^\top W \, \mathbf{z}_g^A, \tag{S39}$$

which is positive when spots with high expression of $g$ neighbor spots with high accessibility of its linked peaks. The same $W$ was used for source and target. All 2,163 genes in the shared source–target RNA feature set with at least one thresholded gene–peak link were analyzed (main-text Fig. 3g). Spatial coordinates enter only through $W$ and the shared lattice, not through the expression or accessibility scores.

#### S9.14 NMF dopamine program and its convergence with the dopamine component

**NMF dopamine program** Non-negative matrix factorization (NMF) (Lee and Seung 1999) was applied to the spatial transcriptomics data from the RNA–MSI source section. NMF components were learned from transcriptomic features alone, without using measured dopamine or `SpatialJEPA`-derived quantities.

We fitted NMF to the `SCTransform`-corrected count matrix using all ST features, prior to the target-compatibility and spatial-variability filtering described in main-text Methods (*RNA–MSI preprocessing and feature selection*). Each decomposition produced a spatial score for every component at each spot, together with corresponding gene loadings. We evaluated ranks from 1 to 29 using a generalized Kullback–Leibler divergence objective with multiplicative updates, NNDSVD-a initialization, a maximum of 1,000 iterations, tolerance $10^{-4}$, and a fixed random seed.

For each decomposition, we quantified the spatial association between each component and measured dopamine using bivariate Moran's $I$. For component $k$, let $\tilde{\mathbf{u}}_k$ denote its $z$-scored spatial scores and $\tilde{\mathbf{y}}$ the $z$-scored measured dopamine values, the statistic is

$$I_k = \frac{n}{S_0} \frac{\tilde{\mathbf{u}}_k^\top W \tilde{\mathbf{y}}}{\tilde{\mathbf{u}}_k^\top \tilde{\mathbf{u}}_k} \tag{S40}$$

Here, $n$ is the number of spots, $W$ is the binary adjacency matrix of the Visium hexagonal lattice with six neighbors per spot, and $S_0 = \sum_{ij} w_{ij}$.

For downstream comparisons, we selected the component with the highest bivariate Moran's $I$ across all tested decompositions and used its gene loadings as the reference transcriptomic ranking.

**Convergence of NMF and PLS dopamine programs** Convergence between the NMF-derived dopamine-associated program and the dopamine component was evaluated over genes shared by both analyses. On the NMF side, genes were ranked by their specificity to component 22, defined as the softmax across NMF components of each gene's loading after scaling each component to a maximum of one (temperature 0.1), so that genes loading strongly on all components rank low. On the component side, genes were ranked by their C2 RNA weight. The 50 most NMF-specific genes were compared with the remaining shared genes using a rank-based AUROC, and enrichment of their overlap with the 50 highest-weight C2 genes was tested by a hypergeometric test over the shared-gene universe.

#### S9.15 Shared components and metabolite weights in RNA-only targets

Figure S19 summarizes how the components of main-text Figs. 4–5 are defined on the spatial RNA–MSI source and read out in RNA-only targets, and how metabolite weights are obtained in targets that carry no MSI (main-text Methods). The components are fixed directions in the student latent space, so every dataset is scored on the same axes; target metabolite weights are carried over from the source's measured MSI through optimal-transport pairing, not imputed by a decoder fitted on the source.

#### S9.16 Regulatory chromatin analyses

To test whether the dopamine component preferentially weighted cell-type-specific regulatory elements, ATAC peaks were ranked by their signed C2 weights and analyzed by pre-ranked GSEA with `gseapy` (Fang et al. 2023). We used the `prerank` method with 1,000 permutations, a minimum set size of 1, and a maximum set size equal to the number of ranked peaks, retaining cCRE sets with few represented elements. The gene set comprised peaks overlapping D2-MSN cCREs from the mouse cerebrum atlas (Li et al. 2021). All 66,520 accessible peaks were ranked, without reference to cCRE identity. Peak weights were computed against the C2 score after regressing it, across nuclei, on log ATAC depth and per-nucleus GC bias, the count-weighted mean GC fraction of each nucleus's peak signal.

We next tested TF motif enrichment among the cCRE peaks contributing most strongly to the GSEA signal. The foreground comprised the GSEA leading-edge cCRE peaks, i.e. those ranked at or before the maximum of the running enrichment score, whose absolute weights fell in the higher-mean component of a two-component Gaussian mixture; the remaining D2-MSN cCREs formed the background. Sequences from both sets were scanned on both strands with MOODS using JASPAR 2024 CORE vertebrate position-weight matrices (Khan 2021), converted to log-odds scores with a pseudocount of 0.8 against uniform background base frequencies, and thresholded at a per-match $p$-value of $5 \times 10^{-4}$. Motif scores were binarized to per-peak presence. To control for base composition, each motif was tested by logistic regression of motif presence on foreground membership with GC fraction and its square as covariates (one-sided Wald test), and by one-sided Fisher's exact tests against three backgrounds redrawn from the cell type's cCRE peaks to match the

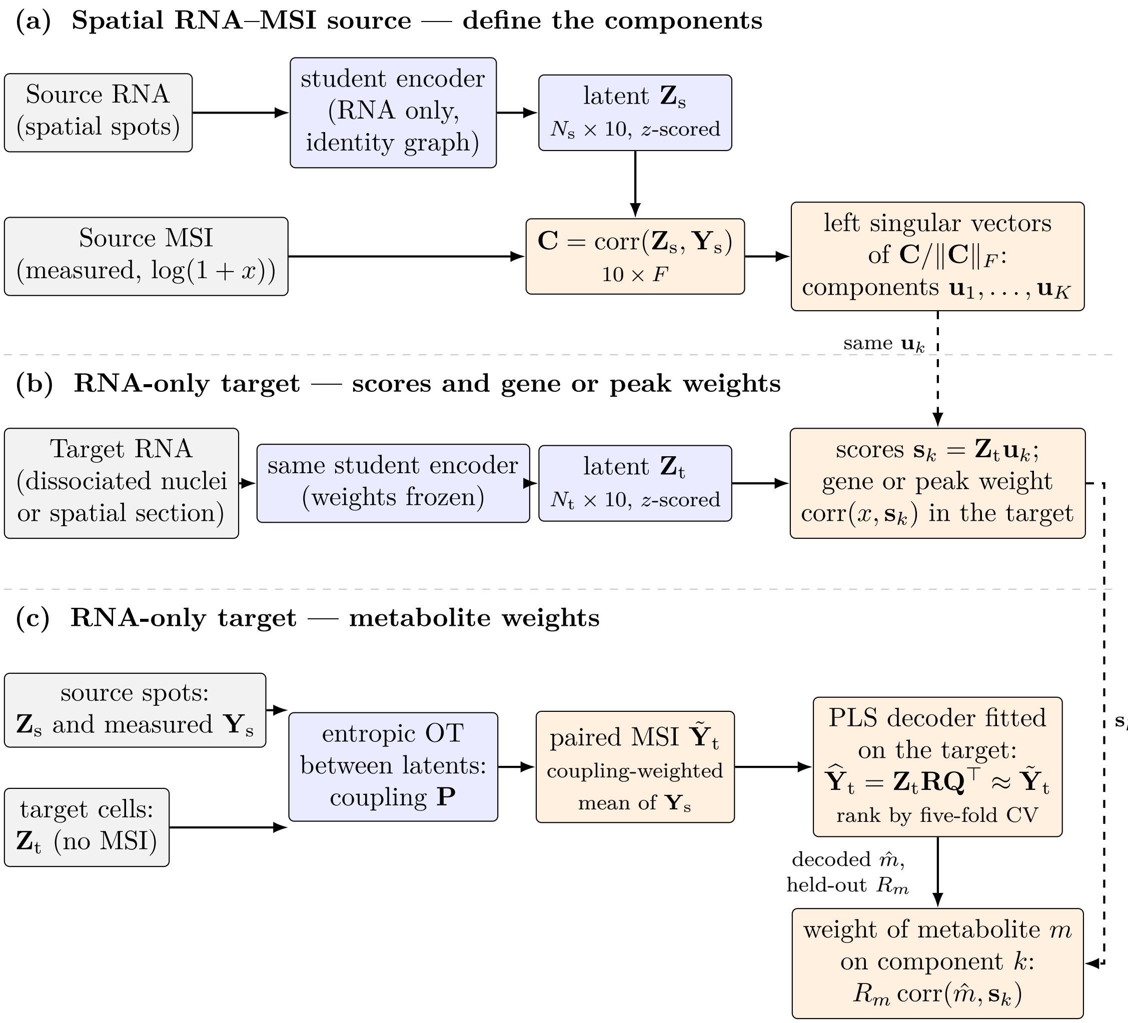

**Figure S19: Shared components and metabolite weights in RNA-only targets.** **(a)** On the spatial RNA–MSI source, the graph-masked student encodes each spot's RNA into a ten-dimensional latent $\mathbf{Z}_s$. The correlations $\mathbf{C}$ between the latent dimensions and the $F$ measured MSI features are decomposed by singular value decomposition; the leading left singular vectors $\mathbf{u}_k$ ($K = 6$ for the mouse model, 5 for the human model) are the components. Dopamine and target data play no part in the decomposition. **(b)** An RNA-only target is encoded by the same frozen student, and each cell is scored on the same directions, $\mathbf{s}_k = \mathbf{Z}_t\mathbf{u}_k$. Gene weights (and, in the 10x Multiome target, peak weights) are correlations with $\mathbf{s}_k$ across the target's own cells. On the P22 spatial target the teacher latent, computed with the P22 spatial graph, is scored on the same components. **(c)** Target cells are paired with source spots by entropic optimal transport between the two latents, and each cell receives the coupling-weighted mean of its paired spots' measured MSI; standardized per metabolite, these paired profiles form $\tilde{\mathbf{Y}}_t$. For the human model, pairing is done separately with each source section and the paired profiles are averaged over sections. A PLS decoder is then fitted on the target, with the standardized target latent $\mathbf{Z}_t$ as predictors and $\tilde{\mathbf{Y}}_t$ as responses. It has the matrix form of Fig. S18a, $\widehat{\mathbf{Y}}_t = \mathbf{Z}_t\mathbf{R}\mathbf{Q}^\top$ (centered), so it maps each cell's latent to a predicted value of every metabolite. The decoded metabolite $\hat{m}$ is the column of $\widehat{\mathbf{Y}}_t$ for metabolite $m$, from the decoder fitted on all target cells. Like the component score $\mathbf{s}_k$, it is a linear function of the latent, and their correlation across target cells gives the unscaled weight. This correlation is multiplied by $R_m$, the decoder's accuracy for metabolite $m$ on cells it was not fitted on. The target cells are split into five folds (spatially compact blocks from $k$-means clustering of spot coordinates on the P22 spatial target; random folds on the Multiome and CaH nuclei). For each candidate rank, the decoder is refitted with each fold held out and predicts that fold. The rank is the smallest whose mean held-out $R^2$, pooled over metabolites, lies within one standard error of the best. $R_m$ is the correlation, across all target cells, between the held-out predictions at that rank and the paired values of metabolite $m$, set to zero when negative. The weight of metabolite $m$ on component $k$ is $R_m\,\mathrm{corr}(\hat{m}, \mathbf{s}_k)$.

foreground GC distribution (three peaks per foreground peak within 0.02-wide GC bins). $P$-values were Benjamini–Hochberg-corrected across motifs, and a motif was considered enriched when the logistic test and at least two of the three matched draws reached $q < 0.05$. Enriched motifs were ordered by their GC-adjusted odds ratio, the exponentiated foreground coefficient of the logistic model. The within-cell-type background controls for differences in baseline motif density among regulatory elements while testing differential enrichment of known motifs.

To obtain a complementary RNA-side regulatory signature, a two-component Gaussian mixture was fitted to the absolute C2 RNA gene weights. Genes with posterior probability $\geq 0.5$ of belonging to the higher-mean component were retained. Evidence for a two-component separation was assessed by the difference in Bayesian information criterion between the one- and two-component models. The retained genes were then tested for TF-regulon over-representation using Enrichr through `gseapy` (Chen et al. 2013), with the `TRANSFAC_and_JASPAR_PWMs` library and all detected RNA features as the background universe.

Cross-modal concordance was assessed by identifying TFs supported by both RNA-regulon enrichment and D2-MSN ATAC-motif enrichment. Each TF was assigned the most stringent combination of nominal ($p < 0.05$) or Benjamini–Hochberg-adjusted ($q < 0.05$) significance reached across the two analyses. To place motif accessibility on the source tissue (Fig. S13d), per-nucleus motif deviations were computed with chromVAR (Schep et al. 2017) as implemented in pychromVAR (Li 2023), over all 66,520 peaks of the 23,275 Multiome nuclei, with JASPAR 2024 CORE vertebrate motif matches at $p < 5 \times 10^{-5}$ and 50 background peak sets matched for GC content and mean accessibility. Deviation $z$-scores were residualized on log ATAC depth. The nuclei were then paired with all 2,526 source spots by the entropic OT used for the metabolite weights (main-text Methods, *Metabolite weights in targets without MSI*), and each spot received the coupling-weighted mean motif accessibility of its paired nuclei. JASPAR versions of the same motif were averaged. Each map was summarized by the AUROC separating striatal from other spots and compared with 20 maps in which the nuclei latent was randomly rotated before pairing.

#### S9.17 SEA-AD Xenium mapping

**SEA-AD Xenium spatial mapping** For spatial visualization, we additionally analyzed SEA-AD CaH Xenium section H21.33.002.CX18.CaH.02.01, containing 48,717 cells. Because oligodendrocytes are concentrated in caudate white matter, their spatial distribution was used as a reference for white-matter-rich tissue. We estimated a continuous oligodendrocyte-density field by fitting a two-dimensional Gaussian kernel density estimator (`scipy.stats.gaussian_kde`) to the coordinates of oligodendrocyte-annotated cells and evaluating the fitted density at every cell coordinate in the section. The KDE bandwidth was set to 0.25 times the default Scott-rule bandwidth to preserve local variation across the gray–white matter boundary. For computational efficiency, up to 5,000 oligodendrocyte coordinates were subsampled using a fixed random seed. Density values were min–max scaled to the interval $[0, 1]$ for visualization (main-text Fig. 5e).

Dopamine was carried to the Xenium cells from the nuclei. A ridge regression, with its penalty selected by efficient leave-one-out cross-validation, was fitted on the 40,000 subsampled nuclei to predict their OT-transferred dopamine from the $z$-scored expression of the 241 genes shared by the source model and the Xenium panel (five-fold cross-validated $r = 0.77$), and applied to the Xenium cells' processed expression. Transferred dopamine was visualized after averaging each cell together with its 30 nearest spatial neighbors to emphasize tissue-scale variation (main-text Fig. 5f). Finally, we projected the Xenium RNA profiles onto C1 using the snRNA-seq C1 gene weights. Xenium transcript counts were normalized to $10^4$ per cell, $\log(1 + x)$-transformed, and $z$-scored per gene within the Xenium section, with values clipped at 10. Of the 1,057 RNA genes of the source model, 241 were measured by Xenium. For each cell $i$, the projected score was calculated as

$$s_i = \frac{\sum_{g \in \mathcal{S}} w_g \tilde{x}_{ig}}{\sum_{g \in \mathcal{S}} |w_g|}, \tag{S41}$$

where $\mathcal{S}$ denotes the 241 shared genes, $w_g$ the snRNA-seq C1 weight of gene $g$, and $\tilde{x}_{ig}$ the processed Xenium expression, $z$-scored per gene within the section. Projected C1 scores were smoothed over the same spatial neighborhoods for visualization (main-text Fig. 5g). Together, these maps place the transferred metabolic and multimodal quantities within the oligodendrocyte-defined tissue context.

**Astrocyte C1 on the Xenium section** To display astrocyte C1 in tissue, astrocyte C1 was transferred to Xenium tissue using ridge regression trained on snRNA-seq astrocytes with the 464 shared Xenium panel genes. Xenium predictions were adjusted for transcript depth within each section and averaged over each astrocyte's 30 nearest astrocytes for visualization (main-text Fig. 5h).

**Spatial cross-correlation among the Xenium maps** To quantify the spatial relationships among the maps in main-text Fig. 5e–g, we computed bivariate Moran's $I$ between each pair using Eq. S40, with $W$ defined as the binary 6-nearest-neighbor adjacency of cell coordinates and symmetrized by taking the union of directed neighbor relations. Because $W$ is symmetric, each pair was reported once. Values were computed both for the maps as plotted and before the 30-neighbor averaging applied to transferred dopamine and C1; the oligodendrocyte-density field was unchanged. Astrocyte C1 (main-text Fig. 5h), defined only for astrocytes, was compared with projected C1 over the section's 6,609 astrocytes, with $W$ the symmetrized 6-nearest-neighbor adjacency among astrocytes and projected C1 taken at the astrocytes' positions; as plotted, both fields are 30-neighbor means.
